# Deciphering the molecular mechanisms underlying anti-pathogenic potential of a polyherbal formulation Enteropan^®^ against multi-drug resistant *Pseudomonas aeruginosa*

**DOI:** 10.1101/2024.01.02.573864

**Authors:** Sweety Parmar, Gemini Gajera, Nidhi Thakkar, Hanmanthrao S Palep, Vijay Kothari

**Author notes:** Contributed equally (Sweety Parmar, Gemini Gajera, Nidhi Thakkar).

## Abstract

Anti-pathogenic potential of a polyherbal formulation Enteropan**^®^** was investigated against a multidrug resistant strain of the bacterium *Pseudomonas aeruginosa*. Enteropan-pre-exposed *P. aeruginosa* displayed reduced (∼70%↓) virulence towards the model host *Caenorhabditis elegans*. Enteropan affected various traits like biofilm formation, protein synthesis and secretion, quorum-modulated pigment production, antibiotic susceptibility, nitrogen metabolism etc. in this pathogen. *P. aeruginosa* could not develop complete resistance to the virulence-attenuating activity of Enteropan even after repeated exposure to this polyherbal formulation. Whole transcriptome analysis showed 17% of *P. aeruginosa* genome to get differentially expressed under influence of Enteropan. Major mechanisms through which Enteropan exerted its anti-virulence activity were found to be generation of nitrosative stress, oxidative stress, envelop stress, quorum modulation, disturbance of protein homeostasis and metal homeostasis. Network analysis of the differently expressed genes resulted in identification of 10 proteins with high network centrality as potential targets from among the down-regulated genes. Differential expression of genes coding for five (*rpoA, tig, rpsB, rpsL*, and *rpsJ*) of these targets was validated through RT-PCR too, and they can further be pursued as potential targets by various drug discovery programmes.

## Introduction

Antibiotic-resistant strains of the gram-negative bacterial pathogen *Pseudomonas aeruginosa* are responsible for considerable morbidity and mortality globally (Kunz Coyne et al., 2022). As per CDC’s Antibiotic Resistance Threat Report (2019), multidrug resistant (MDR) *Psuedomonas aeruginosa* was responsible for an estimated 32,600 hospitalizations, and 2700 deaths in 2017. Healthcare costs in the US alone attributed to these infections were estimated to be 767 million USD. According to a study conducted by Wattal et al. in 2020, India holds a leading position globally with respect to consumption of antibiotics for human use. This heavy antibiotic usage contributes significantly towards antibiotic resistance, resulting in a substantial increase in mortality among newborns who contract sepsis caused by multidrug-resistant pathogens. *P. aeruginosa* displays versatility with respect to types of infections it causes, as it has been involved in pneumonia, urinary tract infections, bloodstream infections, and surgical site infections. Hospitalized patients on ventilators, those with catheters, surgical wounds or burns are particularly at higher risk of contracting *P. aeruginosa* infection. While treating MDR *P. aeruginosa* infections, the choice of available effective antibiotics remains quite narrow, and hence there is an urgent need for discovery and development of novel antibacterial agents/ formulations against this notorious pathogen (Diggle and Whiteley, 2020).

Traditional medicine (TM) formulations can be a potential source of novel leads against bacterial pathogens including *P. aeruginosa*. Since TM often rely on polyherbal formulations (Parasuraman et al., 2014) for treatment, and these polyherbal formulations display ‘multiplicity of targets’ (Makhoba et al., 2020) against susceptible pathogens, investigating the effect of such formulations in the pathogens at whole metabolome or transcriptome level can result in discovery of new molecular targets in pathogens. While dearth of novel cellular and molecular targets has been one of the major hurdles in new antibiotic discovery (Kumar et al., 2021; Ruparel et al., 2023), elucidating the anti-pathogenic potential of polyherbal formulations at molecular level can be quite relevant. Often these polyherbal formulations exhibit anti-virulence effect (Allen et al., 2014) rather than directly inhibiting growth of the target pathogens. They may do so by affecting expression of non-essential genes (i.e., other than housekeeping genes) in the pathogens.

Present study investigated one polyherbal formulation (Enteropan) for its anti-pathogenic potential against a MDR *P. aeruginosa*. This formulation or its component plant extracts are traditionally being prescribed for treatment of irritable bowel syndrome, diarrhea, dysentery, and other gastrointestinal problems (https://www.palepmrf.com/pdf/Enteropan_IBS.pdf).

## Methods

### Test Formulation

Test formulation Enteropan^®^ was procured from Dr. Palep’s Medical Research Foundation Pvt. Ltd, Mumbai. All the nine ingredient plants and their parts whose hydroalcoholic extracts have been mixed to prepare this formulation are listed in Table 1. We obtained the polyherbal mix from the manufacturer in dried powder form without any bulking agent, and mixed 4 g of it in 8 mL of DMSO (Merck). The DMSO-soluble fraction of Enteropan was found to be 71.17% ± 3.07. After separating the insoluble fraction through centrifugation, the remaining DMSO-dissolved fraction was stored under refrigeration.

**Table 1.** Ingredients of Enteropan formulation

| Scientific name | Common English name (Indian name) | Part used | Proportion in the polyherbal mix (mg/ capsule) |
| --- | --- | --- | --- |
| *Aegle marmelos* | Wood apple (Bael) | Leaves | 100 |
| *Myristica fragrans* | Nutmeg (Jaiphal) | Fruit | 15 |
| *Zingiber officinale* | Ginger (Sunthi) | Rhizome | 60 |
| *Aconitum heterophyllum* | Indian Atees (Ativisha) | Bulb | 30 |
| *Coriandrum sativum* | Coriander (Dhanyak) | Seeds | 40 |
| *Cyperus rotundus* | Nutgrass (Nagarmotha) | Rhizome | 25 |
| *Vertiveria zizaniodes* | Khas Khas grass (Usheer) | Roots | 50 |
| *Punica granatum* | Pomegranate (Dadim) | Rind | 50 |
| *Holarrhena antidysentrica* | Star gooseberry (Kutaj) | Bark | 27 |

### Test organisms

*P. aeruginosa* strain was sourced from our internal lab culture collection. Its antibiogram (Table S1) generated through disc diffusion assay revealed it to be resistant to three different classes of antibiotics i.e., co-trimoxazole (combination of trimethoprim and sulfamethoxaole), streptomycin (aminoglycoside), and augmentin (combination of amoxicillin and clavulanic acid). Antibiotic susceptibility of this strain to kanamycin was classified as ‘intermediate’. Pseudomonas broth (20 g/L peptic digest of animal tissue, 10 g/L potassium sulphate, 1.4 g/L magnesium chloride, 3% v/v glycerol, pH 7.0±0.2) or agar (HiMedia) was used to cultivate this bacterium. Inoculum density of this bacterium to be used in all experiments was adjusted at OD_625_ = 0.08-0.10 to achieve equivalence to McFarland turbidity standard 0.5.

*E. coli* OP50 procured from LabTIE B.V. the Netherlands was used as food for *Caenorhabditis elegans*, while maintaining the worm on NGM (Nematode Growing Medium; 3g/L NaCl, 1M CaCl_2_, 1M MgSO_4_, 2.5g/L peptone, 5 mg/mL cholesterol, 1 M phosphate buffer of pH 6, 17g/L agar-agar).

The nematode worm *Caenorhabditis elegans* (N2 Bristol, procured from IIT Gandhinagar) was used as a model host for *P. aeruginosa.* The worm was maintained on NGM agar plates. Worm synchronization was done as described in literature (Corsi et al., 2015) and in our previous studies (Patel et al., 2019a, Joshi et al., 2019) too. Prior to all *in vivo* assays, worms were kept without food for two days to make them gnotobiotic.

### In vivo assays

Four different types of *in vivo* assays performed are described below. Each of them involved live-dead counting over a period of five days under a microscope (4X) with halogen light source. On the last day of experiment, when plates could be opened, death was confirmed by touching them with a straight wire, wherein no movement was considered as confirmation of death. In each well, there were 10 worms in M9 buffer (3g/ L KH_2_PO_4_; 6g/L Na_2_HPO_4_; 5 g/L NaCl), which were challenged with *P. aeruginosa* by adding 100 µL (OD_764_= 1.50 ± 0.05) of bacterial culture grown in Pseudomonas broth for 21±1 h at 35±0.5°C.

#### Anti-pathogenic assay

Ten worms of L3-L4 stage contained in 900 µL M9 buffer were challenged with *P. aeruginosa* (100 µL of the culture broth) in absence or presence of Enteropan (50-1000 µg/mL), wherein neither the bacterium nor the worm were pre-exposed to Enteropan. Incubation was done at 22⁰ C for 5 days with live-dead microscopic count once a day.

#### Anti-infective assay

*P. aeruginosa* was grown at 35⁰ C for 20-22 h in Pseudomonas broth with or without Enteropan (5-1000 µg/mL). Post incubation, 100 µL of the culture broth was mixed with 900 µL of M9 buffer containing 10 worms (L3-L4 stage) in a 24-well plate (surface non-treated; HiMedia). This plate was incubated at 22⁰ C for 5 days (Durai *et. al.* 2013; Patel et al., 2020).

#### Prophylactic assay

Gnotobiotic worms were incubated in M9 buffer supplemented with Enteropan (5-1000 µg/mL) for 96 h. Following incubation, these worms were washed with M9 buffer twice, and then challenged with *P. aeruginosa* not pre-exposed to Enteropan (100 µL of the culture broth) in 24-well plates (HiMedia). Worm survival was monitored over a 5-day period under microscope (Patel *et. al.*, 2018).

#### Post-infection assay

Ten worms contained in M9 buffer were first challenged with pathogen, and after allowing *P. aeruginosa* (100 µL of the culture broth) for 3 or 6 h to establish infection, Enteropan (50-1000 µg/mL) was added into the well as a possible post-infection therapy. Survival of worms was observed through a live-dead count under microscope over 5-days (Patel *et. al.*, 2019b).

Appropriate controls were included in all above experiments as relevant:

Sterility Control: Sterile M9 buffer containing neither bacteria nor worms

Survival Control: M9 buffer containing 10 worms (no bacteria added)

Toxicity Control: 10 worms in M9 buffer supplemented with Enteropan

Infection Control: 10 worms in M9 buffer + 100 µL of the *P. aeruginosa* culture broth (OD_764_ = 1.50 ± 0.05). These wells did not contain any plant extract.

Vehicle Control: 0.5%v/v DMSO was used in place of Enteropan.

Positive Control: Standard antibiotics employed as positive controls are detailed in the figure legends.

### In vitro assays

#### Growth and pigment quantification

The broth dilution assay was used to evaluate *P. aeruginosa’s* growth and quorum sensing (QS)-regulated pigment synthesis in the presence or absence of the test formulation. Different concentrations (ranging from 5 to 1000 μg/mL) of Enteropan formulation were used to challenge the organism. The growth media employed was Pseudomonas broth, into which bacterial inoculum set to 0.5 McFarland turbidity standard was added at 10%v/v, followed by incubation at 35° C for 20-22 h, with intermittent shaking. The experiment also contained an appropriate vehicle control with DMSO (0.5%v/v) and an abiotic control with extract and growth medium but no inoculum.

Bacterial growth was measured photometrically at the end of the incubation by measuring the culture density at 764 nm (Agilent Cary 60 UV-visible spectrophotometer) (Joshi et al., 2016). Following this, pigment was extracted and quantified in accordance with the procedure outlined below for each pigment.

One mL of culture broth was mixed in a 2:1 ratio with chloroform (Merck, Mumbai), followed by centrifugation (15,300 g) for 10 min. This resulted in formation of two immiscible layers. OD of the upper aqueous layer containing the yellow-green fluorescent pigment pyoverdine was measured at 405 nm. Pyoverdine Unit was calculated as OD_405_/OD_764_. The lower chloroform layer containing the blue pigment pyocyanin was mixed with 0.1 N HCL (20%v/v; Merck). This caused a change of colour from blue to pink. This was followed by centrifugation (15,300 g) for 10 min, and OD of upper layer acidic liquid containing pyocyanin was quantified at 520 nm. Pyocyanin Unit was calculated as OD_520_/OD_764._

#### Biofilm assays

Biofilm formation is an important virulence trait, and hence effect of Enteropan on biofilm forming ability of *P. aeruginosa*, as well as on pre-formed biofilm was investigated. A flow diagram depicting all four different biofilm assays is included in supplementary file (Figure S2). Biofilm quantification was achieved through crystal violet assay (Hirshfield et al., 2008). Biofilm viability was assessed through MTT assay (Trafny et al., 2013).

For the crystal violet assay, the biofilm containing tubes (after discarding the inside liquid) were washed with phosphate buffer saline (PBS) in order to remove all non-adherent (planktonic) bacteria, and air-dried for 15 min. Then, each of the washed tubes was stained with 1.5 mL of 0.4% aqueous crystal violet (Central Drug House, Delhi) solution for 30 min. Afterwards, each tube was washed twice with 2 mL of sterile distilled water and immediately de-stained with 1.5 mL of 95% ethanol. After 45 min of de-staining, 1 mL of de-staining solution was transferred into separate tubes, and read at 580 nm (Agilent Cary 60 UV-Vis).

For the MTT assay, the biofilm-containing tubes (after discarding the inside liquid) were washed with PBS in order to remove all non-adherent (planktonic) bacteria, and air-dried for 15 min. Then 1.8 mL of minimal media (sucrose 15 g/L, K_2_HPO_4_ 5 g/L, NH_4_Cl 2 g/L, NaCl 1 g/L, MgSO_4_ 0.1 g/L, yeast extract 0.1 g/L, pH 7.4 ± 0.2) was added into each tube, followed by addition of 200 μL of 0.3% MTT [3-(4,5-Dimethylthiazol-2-yl)-2,5-diphenyltetrazolium bromide; HiMedia]. Then after 2 h incubation at 35°C, all liquid content was discarded, and the remaining purple formazan derivatives were dissolved in 2 mL of DMSO and measured at 540 nm.

#### Nitrite estimation

Quantification of nitrite in bacterial culture was achieved through a colorimetric assay using modified Griess reagent (Misko et al., 1993). Supernatant (250 µL) obtained from centrifugation (13,500 g; 25°C; 10 min) of *P. aeruginosa* culture grown in presence or absence of Enteropan was mixed with 250 µL of Griess reagent (1X; Sigma-Aldrich), followed by 15 min incubation in dark at room temperature. Absorbance of the resulting pink colour was measured at 540 nm. This absorbance was plotted on the standard curve prepared using NaNO_2_ (0.43-65 µM) to calculate nitrite concentration. Sodium nitroprusside (Sigma Aldrich) was used as a positive control, as it is known to generate nitrosative stress in bacteria (Barnes et al., 2013). Deionized water was used as negative control.

#### Antibiotic susceptibility test

Antibiogram of *P. aeruginosa*’s overnight grown culture in Pseudomonas broth in presence or absence of Enteropan was generated through disc diffusion assay in accordance to Clinical and Laboratory Standards Institute (CLSI) guidelines (Simner et al., 2022). Cells grown in Pseudomonas broth were separated through centrifugation (13600 g) and washed with phosphate buffer (pH 7.0±0.2) followed by centrifugation. The resulting cell pellet was used to prepare inoculum for subsequent disc diffusion assay by suspending the cells in normal saline and adjusting the OD_625_ between 0.08-0.10. This inoculum (100 µL) was spread onto cation-adjusted Muller-Hinton agar (HiMedia) plates (Borosil; 150 mm) followed by placing the antibiotic discs (Icosa G-I MINUS; HiMedia, Mumbai) on the agar surface. Incubation at 35°C was made for 18±1 h, followed by observation and measurement of zone of inhibition.

#### Protein estimation

Extracellular protein present in bacterial culture (grown in presence or absence of Enteropan) supernatant, and intracellular protein in the cell lysate was quantified through Folin-Lowry method (Lowry et al., 1951; Dulley and Grieve, 1975). After measuring cell density, one mL of *P. aeruginosa* culture was centrifuged (13,600 g), and the resulting supernatant was used for extracellular protein estimation. The remaining cell pellet was subjected to lysis (Mishra et al., 2019) for release of intracellular proteins. Briefly, the cell pellet was washed with phosphate buffer (pH 7.4), and centrifuged (13,600 g). Resulting pellet was resuspended in 1 mL of chilled lysis buffer (0.876 g NaCl, 1 mL of Triton X 100, 0.5 g sodium deoxycholate, 0.1 g sodium dodecyl sulphate, and 0.60 g Tris HCl, in 99 mL of distilled water), and centrifuged (500 rpm) for 30 min at 4°C for agitation purpose. This was followed by further centrifugation (16,000 g at 4°C) for 20 min. Resulting cell lysate (supernatant) was used for protein estimation. Kanamycin (HiMedia; at IC_50_: 200 µg/mL), an aminoglycoside antibiotic known to inhibit bacterial protein synthesis (Suzuki et al., 1970; Ullah and Ali, 2017), was used as a positive control.

### Whole Transcriptome Analysis

To gain insight into the molecular mechanisms by which Enteropan attenuates bacterial virulence, and modulate various traits like pigment production, antibiotic susceptibility, nitrogen metabolism, protein synthesis/ excretion, etc., we compared the gene expression profile of Enteropan-pre-treated *P. aeruginosa* with that of control culture at the whole transcriptome level.

#### RNA extraction

Trizol (Invitrogen Bioservices; 343909) was used to extract RNA from bacterial cells (Jahn et al., 2008). RNA was dissolved in nuclease-free water after precipitation with isopropanol and washing with 75% ethanol. Using the RNA HS assay kit (Thermofisher; Q32851) and adhering to the manufacturer’s instructions, extracted RNA was quantified using a Qubit 4.0 fluorometer (Thermofisher; Q33238). RNA concentration and purity were evaluated using Nanodrop 1000. Finally, RNA was checked on the TapeStation using HS RNA ScreenTape (Agilent) to yield RIN (RNA Integrity Number) values (Table S2).

#### Library preparation

Final libraries were measured using a Qubit 4.0 fluorometer (Thermofisher; Q33238), a DNA HS assay kit (Thermofisher; Q322851), and a Tapestation 4150 (Agilent) using high-sensitivity D1000 screentapes (Agilent; 5067-5582). The acquired sizes of all libraries are reported in Table S3.

#### Genome annotation and functional analysis

FastQC v.0.11.9 (default parameters) was used to undertake a quality assessment of the sample’s raw fastq readings (Andrews, 2010). The reads’ quality was then reevaluated using Fastq v.0.20.1 (Chen et al., 2018) after pre-processing the raw fastq reads with Fastq v.0.20.1.

The *P. aeruginosa* genome (GCA_000006765.1_ASM676v1) was indexed using bowtie2-build (Langmead and Salzberg, 2012) v2.4.2 (default parameters). The processed reads were mapped to the *P. aeruginosa* genome using bowtie2 v2.4.2. Gene counts were determined using feature count v.0.46.1 (Liao et al., 2014) to quantify the aligned reads from the individual samples. Differential expression was estimated using the exact test (parameters: dispersion 0.1) with these gene counts as inputs in edgeR (Robinson et al., 2010). The up-and down-regulated sequences were extracted from the *P. aeruginosa* coding file and annotated using Blast2GO (Conesa and Gotz, 2008) to obtain the Gene Ontology (GO) keywords. These GO terms were used to create gene ontology bar graphs with the wego tool (Ye et al., 2018).

All the raw sequence data has been submitted to the Sequence Read Archive. The relevant accession number is SRX15248092 (https://www.ncbi.nlm.nih.gov/sra/SRX15248092).

### Network analysis

Network analysis was carried out for Enteropan-exposed *P. aeruginosa*’s differentially expressed genes (DEG) fulfilling the dual criteria of log fold-change ≥2 and FDR ≤0.001. List of DEG was fed into the database STRING (v.11.5) (Szklarczyk et al., 2023) for generating the PPI (Protein-Protein Interaction) network. Then the genes were arranged in decreasing order of ‘node degree’ (a measure of connectivity with other genes or proteins), and those above an empirically selected threshold value (19 and 53 for up-and down-regulated genes respectively) were subjected to ranking by cytoHubba (v.3.9.1) plugin (Chin et al., 2014) of Cytoscape (Shannon et al., 2003). Since cytoHubba uses 12 different ranking methods, we considered the DEG being top-ranked by ≥6 different methods (i.e., 50% of the total ranking methods) for further analysis. These top-ranked shortlisted proteins were further subjected to network cluster analysis through STRING, and those which were part of multiple clusters were considered as ‘hubs’ which can be taken up for further confirmation of their targetability. Here ‘hub’ refers to a gene or protein interacting with many other genes/proteins. Hubs thus identified were further subjected to co-occurrence analysis to see whether an anti-virulence agent targeting them is likely to satisfy the criterion of selective toxicity (i.e., targeting the pathogen without harming host). This sequence analysis allowed us to end with a limited number of proteins which satisfied various statistical and biological significance criteria simultaneously i.e. (1) log fold-change ≥2; (2) FDR ≤0.001; (3) relative higher node degree; (4) top-ranking by at least 6 cytoHubba methods; (5) member of more than one local network cluster; (6) high probability of the target being absent from the host.

#### RT-PCR analysis

PCR was used to confirm the differential expression of the possible hubs discovered by network analysis of the DEG (differentially expressed genes) reported from WTA. Primer3Plus (Untergasser et al., 2007) was used to design primers for the target genes (Table 2). These primer sequences were verified for their ability to specifically bind only to the target gene sequence throughout the whole genome file of *P. aeruginosa*. RNA extraction and purity check was executed as described in previous section. The SuperScript^™^ VILO^™^ cDNA Synthesis Kit (Invitrogen Biosciences) was used to generate cDNA. Using gene-specific primers purchased from Sigma-Aldrich, the PCR experiment was carried out employing the temperature profile shown in Table S4. The gene PA3725 (recJ) was kept as an endogenous control. The reaction mix used was FastStart Essential DNA Green Master mix (Roche; 06402712001). Real time PCR assay was performed on Quant studio 5 real time PCR machine (Thermo Fisher Scientific, USA). Sample generation for PCR validation was done independent of that for transcriptome assay.

**Table 2.**
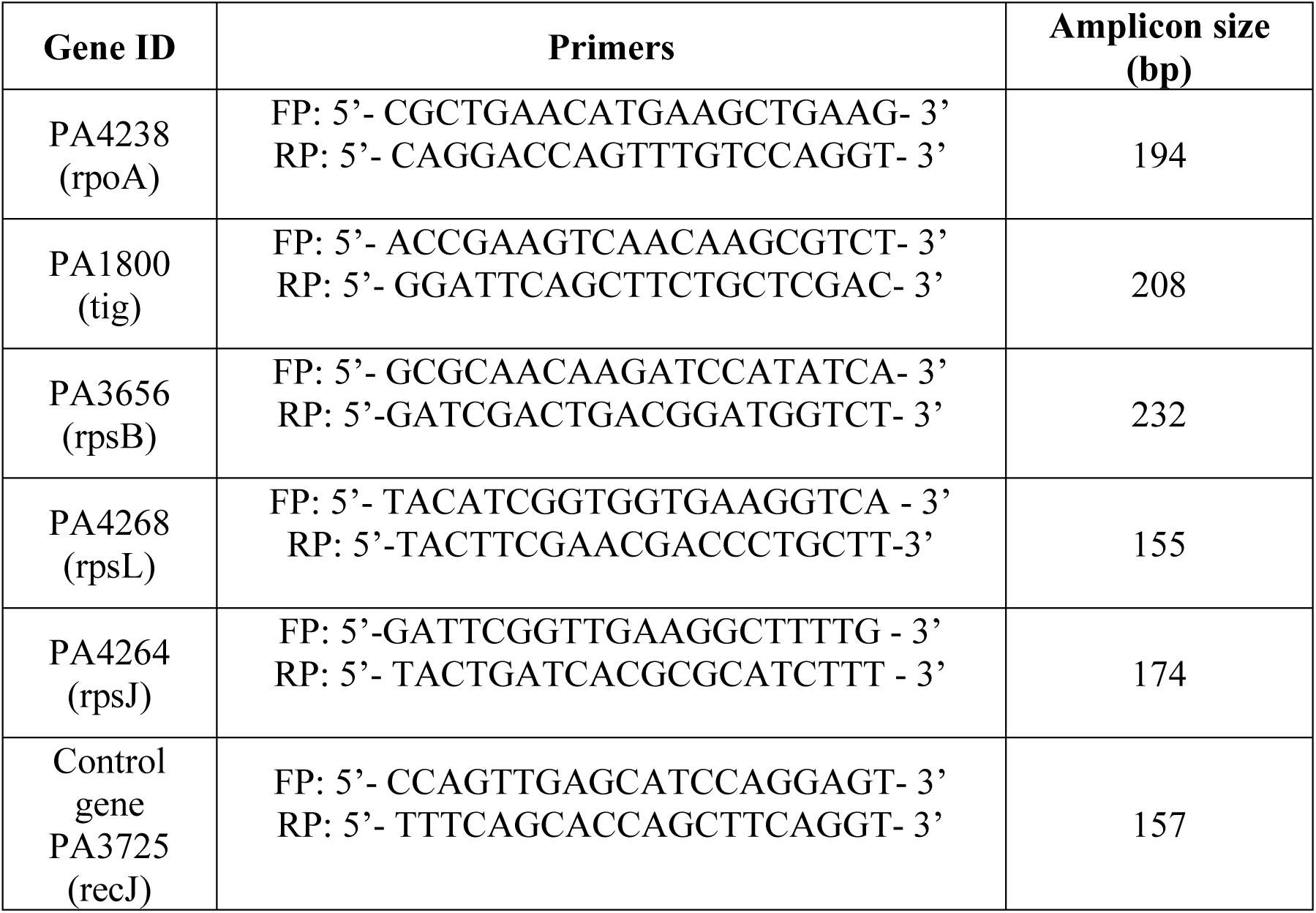
Primer sequences for the target genes

### Statistical analysis

All results reported are means of three or more independent experiments, each performed in triplicate. Statistical significance was assessed through t-test performed using Microsoft Excel^®^, and data with p≤0.05 was considered to be significant.

## Results and Discussion

### In vivo assays

#### *P. aeruginosa* displayed reduced virulence towards *C. elegans* in presence of Enteropan

When *C. elegans* was challenged with *P. aeruginosa* in presence of Enteropan, the bacterium could kill lesser worms than in absence of Enteropan (Figure 1A; Supplementary videos: A-E). The most effective concentration of Enteropan with respect to offering protection to the worm population from bacterial attack was found to be 250 µg/mL. Since higher concentrations offered either at par or lesser protection to the worms, the dose-response relationship here can be said to be non-linear. Irrespective of the magnitude of protection offered to worm population in presence of Enteropan^®^, progeny worms were observed (third day onward) in all experimental and positive control wells, but not in the wells pertaining to vehicle control. It might have occurred that the virulence-attenuated *P. aeruginosa* were used by the worms as food, and that allowed them to reproduce.

**Figure 1.**
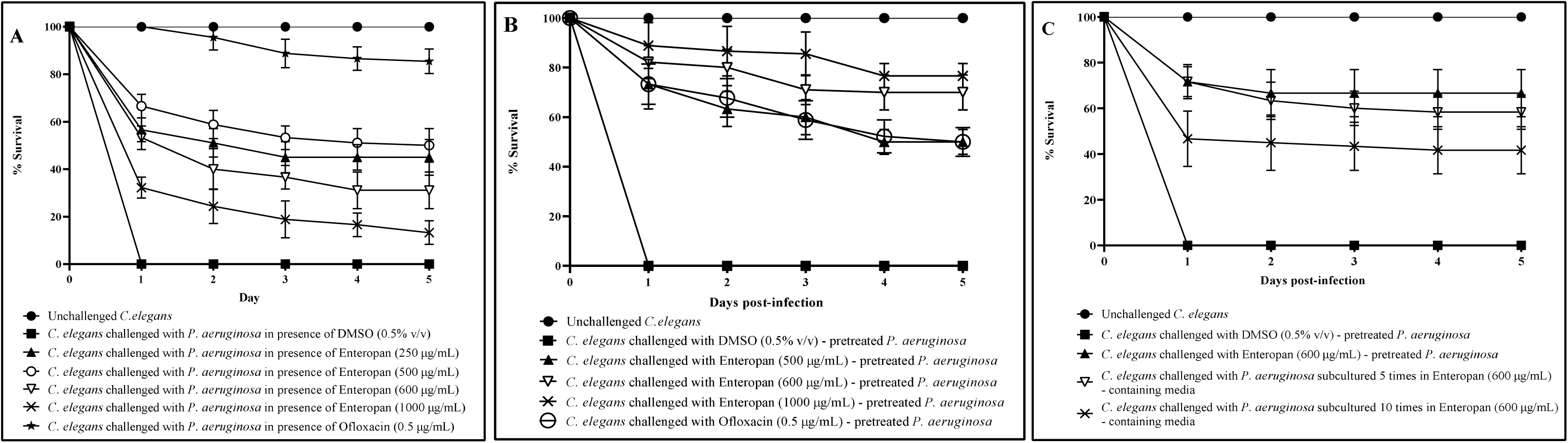
Enteropan attenuates *P. aeruginosa’s* virulence towards the model host *C. elegans.* DMSO present in the ‘vehicle control’ at 0.5%v/v did not affect bacterial virulence. Neither DMSO nor Enteropan showed any toxicity towards the worm population at tested concentrations. To avoid overcrowding in the figures (A-B), we have not shown lines corresponding to concentrations which had no effect on bacterial virulence, and also that for 750 μg/mL, as its virulence-attenuating effect was statistically at part to that of 600 μg/mL. Supplementary videos pertaining to these experiments are available at: osf.io/fnywk. All the % values reported in this figure legend are statistically significant at *p*<0.001. **(A) *P. aeruginosa’s* virulence towards the host worm gets attenuated in presence of Enteropan.** Enteropan conferred a survival benefit on host *C. elegans* at concentrations of 250 μg/mL, 500 μg/mL, 600 μg/mL, 750 μg/mL, and 1000 μg/mL with survival rates of 45% ± 5.47, 50% ± 7.07, 31.11% ± 7.8, 35% ± 8.36, and 13.33% ± 5 respectively. Ofloxacin (0.5 µg/mL) was employed as a positive control and conferred 85.55% ± 5.27 survival benefit on host worm. Progenies (TNTC; too numerous to count) were observed on third day in experimental wells and positive control. See supplementary videos A-E. **(B) Enteropan pre-treatment reduced bacterial virulence towards *C. elegans***. Pre-treatment of bacteria with Enteropan at concentrations of 500 μg/mL, 600 μg/mL, 750 μg/mL, and 1000 μg/mL reduced its virulence towards host worm by 50% ± 5, 70% ± 7.07, 78.88% ± 6, and 76.66% ± 5 respectively as per the 5^th^ day end-point. Ofloxacin (0.5 µg/mL) pre-treatment reduced bacterial virulence towards the host worm by 50% ± 5.77. Progenies were observed on third day in experimental wells corresponding to ≥500 μg/mL Enteropan as well as positive control. See supplementary videos F-I. **(C) *P. aeruginosa* did not develop complete resistance even after repeated exposure to Enteropan**. *P. aeruginosa* obtained after fifth and tenth subculturings in Enteropan (600 μg/mL)-containing media displayed 58.33%±4.04and 43.33%±10.3lesser virulence respectively than extract-non-exposed pathogen.

#### Enteropan pre-treatment reduced bacterial virulence towards *C. elegans*

Enteropan (5-1000 μg/mL) pre-treated *P. aeruginosa* was found to exert lesser virulence against *C. elegans* than its extract not-exposed counterpart. Enteropan concentrations ≤ 250 µg/mL could not compromise *P. aeruginosa*’s ability to kill *C. elegans*, however concentrations ≥ 500 µg/mL did compromise bacterial virulence significantly (Figure 1B; Supplementary videos: F-I). Enteropan pre-treatment of the pathogen at these effective concentrations not only attenuated bacterial virulence, but also supported worm fertility as evidenced by appearance of numerous progenies in the experimental wells by fifth day. The most effective concentration of Enteropan with respect to virulence-attenuating effect was found to be 600 µg/mL, as effect of higher concentrations till 1 mg/mL was statistically not different than that of 600 µg/mL.

After confirming the anti-virulence activity of Enteropan against *P. aeruginosa*, we asked whether this pathogen can develop resistance upon repeatedly getting exposed to the test formulation. To answer this, we subcultured *P. aeruginosa* in Pseudomonas broth supplemented with Enteropan (600 µg/mL) multiple times. Enteropan pre-exposed *P. aeruginosa* thus obtained after fifth and tenth such subculturing were allowed to attack *C. elegans* in M9 buffer (containing no Enteropan). No resistance seemed to have evolved in *P. aeruginosa* till fifth subculturing, however ten subculturings in Enteropan-supplemented media seemed to allow the pathogen to overcome this formulation’s anti-virulence effect marginally (23.3%; Figure 1C). This inability of the pathogen to develop complete resistance against Enteropan might be attributable to the polyherbal nature of the formulation. As the polyherbal formulations can have multiple bioactive compounds in them, they may exert a multiplicity of targets against the susceptible pathogen. To develop resistance in this scenario, the pathogen would be required to develop multiple simultaneous mutations, and that is a quite less probable event biologically as well statistically.

#### Enteropan offered prophylactic protection to *C. elegans*

To investigate whether Enteropan pre-feeding can offer any prophylactic benefit to worm population in face of subsequent pathogen challenge, we allowed *P. aeruginosa* to attack worms pre-fed with Enteropan (5-1000 µg/mL). Enteropan at ≥ 50 µg/mL did confer prophylactic benefit on worm population. While concentrations till 500 µg/mL supported worm survival (13-26%) till 48 h post-pathogen challenge (Figure 2A), higher concentrations supported worm survival (20-23%) till fifth day (Figure 2B; Supplementary videos: J-M). Considering the first-day end point (i.e., by the time the pathogen killed 100% worms in control wells), certain Enteropan concentrations (500-750 µg/mL) performed at par to the positive control ofloxacin.

**Figure 2.**
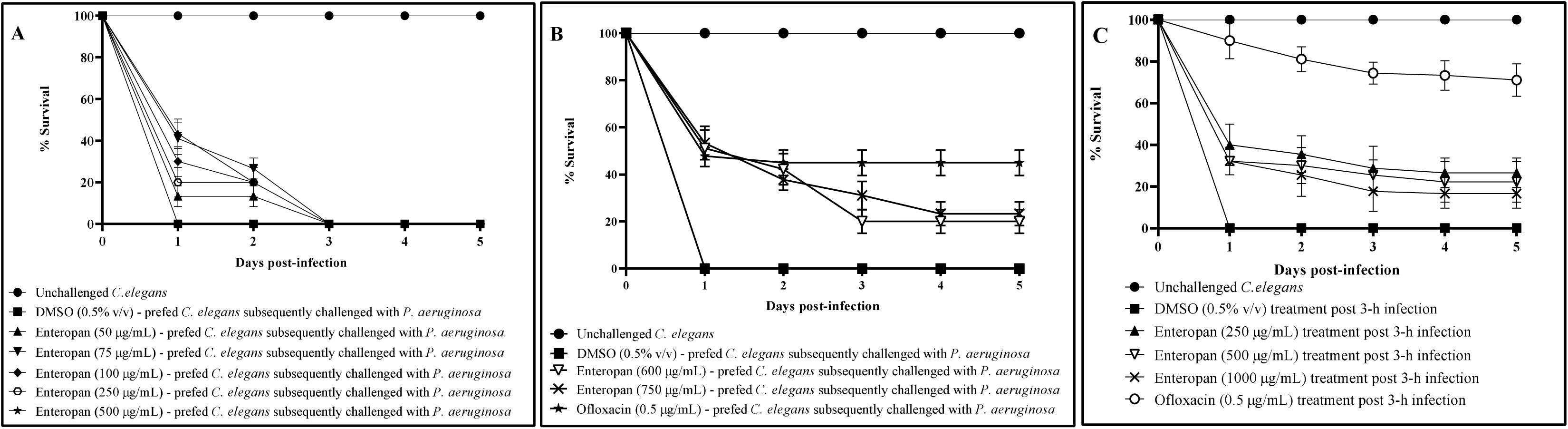
Prophylactic and post-infection therapeutic potential of Enteropan. All the % values reported in this figure legend are statistically significant at *p*<0.001. **(A-B) Enteropan offered prophylactic protection to the worm population against subsequent bacterial challenge.** Worms pre-fed with Enteropan concentrations 50 μg/mL, 75 μg/mL, 100 μg/mL, 250 μg/mL, and 500 μg/mL, registered 13.3% ±5, 26.6% ±5, 20% ±8.66, 17% ± 6.66 and 20% ±8.66 better survival, respectively till the 2^nd^ day in face of subsequent pathogen challenge. Worms pre-fed with higher concentrations of Enteropan at 600 μg/mL, 750 μg/mL, and 1000 μg/mL registered 51.11% ± 7.8, 53.33 ± 7.07 and 43.33% ± 8.66 survival respectively till the end of first day of pathogen challenge. Magnitude of the prophylactic benefit as per fifth day point was 20% ± 5, 23.33 ± 5 and 20% ± 7.07 respectively. Pre-feeding the worms with DMSO (0.5%v/v) did not alter their susceptibility to subsequent bacterial challenge. Ofloxacin (0.5 µg/mL) employed as a positive control conferred 44.9% ± 5.49 prophylactic benefit on the host worms. See supplementary videos J-M. **(C) Enteropan could partially rescue worm population when used as a post-infection therapy**. When pre-infected worms were exposed to Enteropan at 250 μg/mL, 500 μg/mL, 600 μg/mL, 750 μg/mL, and 1000 μg/mL, they scored 26.66% ± 7.0, 22.22% ± 9.71, 11.1% ± 3.33, 13.3% ± 6, and 16.66% ± 7.07 better survival respectively, than control worms. Ofloxacin (0.5 µg/mL) employed as positive control, 3 h post-infection, rescued 71.11% ± 7.81 worms. DMSO (0.5%v/v) did not confer any survival benefit when added post-infection. See supplementary videos N-R.

#### Enteropan is effective as a post-infection therapy

To investigate whether Enteropan is effective as a post-infection therapy, we added Enteropan (50-1000 μg/mL) after 3 or 6 h of mixing bacteria with the worms. While Enteropan addition post 6 h of bacterial attack on worms could not rescue the host (Figure S1), its addition post 3 h of bacterial attack could rescue 10- 27% of the worms (Figure 2C; Supplementary videos: N-R). As a post-infection therapy, Enteropan’s performance did not improve with increase in concentration.

To have a mechanistic insight into the Enteropan-*P. aeruginosa* interaction, we checked effect of Enteropan on various virulence traits of this pathogen *in vitro*, and also compared the gene expression profile of the Enteropan-treated *P. aeruginosa* with that of extract-non-exposed control at the whole transcriptome level.

### *In vitro* experiments

#### Enteropan forced overproduction of quorum sensing (QS)-regulated pigments without affecting the bacterial growth heavily

Enteropan till 100 µg/ml had no effect on *P. aeruginosa* growth. Though 250 µg/ml onwards it had some growth inhibitory effect, magnitude of this inhibitory effect did not increase much with increase in concentration (Figure 3A). Except 5 µg/ml, Enteropan at all tested concentrations enhanced production of QS-regulated pigments (pyoverdine and pyocyanin). Hence concentrations 25-100 µg/ml can be said to have pure quorum-modulatory effect on *P. aeruginosa*. Though in general Enteropan’s effect on pigment production seemed to be dose-dependent, bacterium’s response at 600 µg/ml seemed to deviate from this pattern in case of both the pigments.

**Figure 3.**
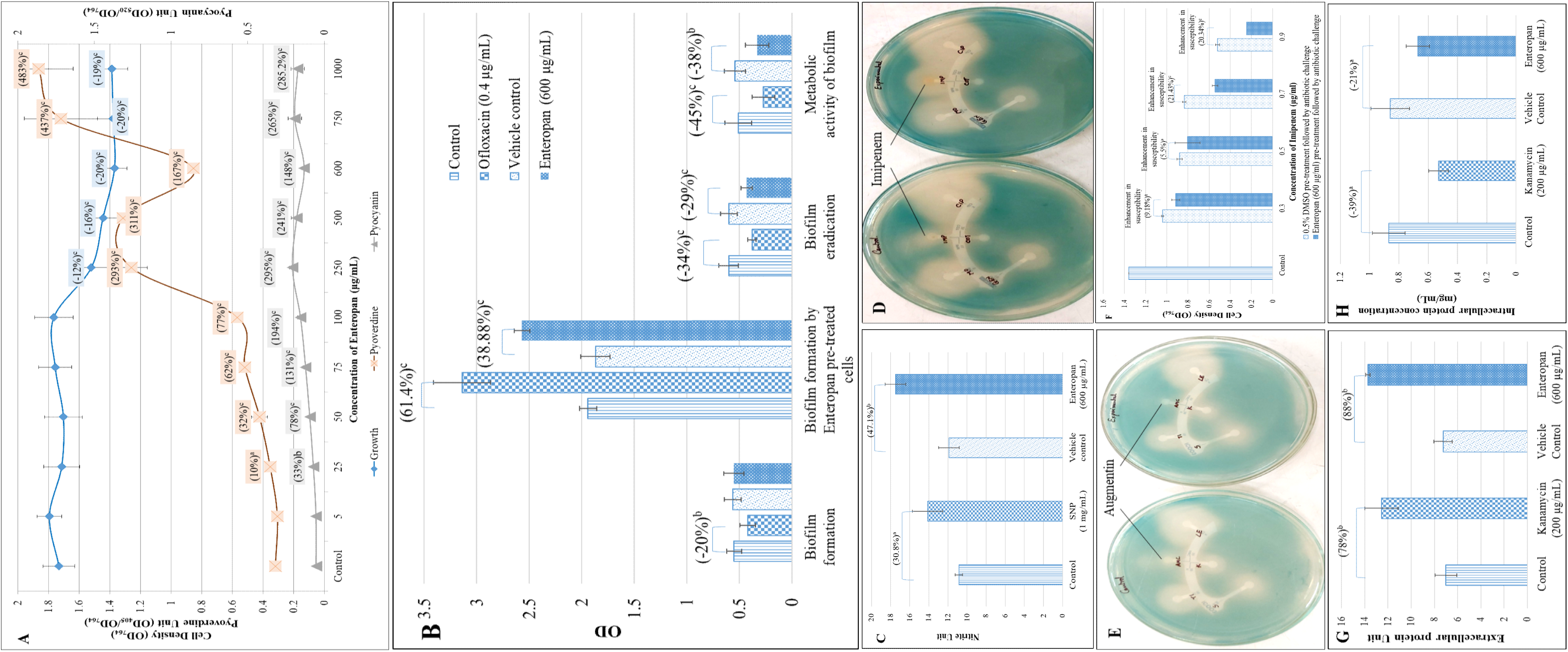
Enteropan’s effect on various phenotypic traits of *P. aeruginosa* revealed through different *in vitro* assays. **(A) Enteropan enhances production of quorum-regulated pigments in *P. aeruginosa,* while exhibiting a mild growth inhibitory effect**. Bacterial growth was measured as OD764. Pyoverdine Unit and Pyocyanin Unit was calculated as the ratio OD405/OD764 and OD520/OD764 (an indication of pyoverdine and pyocyanin production per unit of growth) respectively; ‘Control’ shown in this figure is the vehicle control (0.5%v/v DMSO), which affected neither growth nor pigment production. Ofloxacin (0.5 µg/mL) inhibited growth by 65.6% ±5.32, while inhibiting pigment production completely. **(B) Enteropan’s effect on *P. aeruginosa’s* biofilm formation capability and on pre-formed biofilm.** While P. aeruginosa’s biofilm formation ability remained unaffected in presence of Enteropan, Enteropan-pre-exposed cells subsequently allowed to form biofilm on glass surface accumulated higher biomass. Enteropan when added onto pre-formed biofilm, it could eradicate the biofilm partially, and also reduced the biofilm metabolic activity notably. DMSO (0.5% v/v) used as vehicle control did not affect biofilm of the *P. aeruginosa* in any of these four assays. **(C) *P. aeruginosa* culture accumulated higher extracellular nitrite in presence of Enteropan.** While nitrite concentration in vehicle control (*P. aeruginosa* supplemented with 0.5%v/v DMSO) was at par to that without DMSO, Enteropan caused nitrite concentration in *P. aeruginosa* culture supernatant to rise. Sodium nitroprusside used as positive control caused 30.8% higher nitrite build up in *P. aeruginosa* culture. Nitrite Unit (i.e., Nitrite concentration: Cell density ratio was calculated to nullify any effect of cell density on nitrite production). **(D - E) Enteropan-pretreated cells responded to certain antibiotics differently.** Enteropan-pre-exposed *P. aeruginosa* experienced an increased or decreased susceptibility to imipenem and augmentin respectively, as revealed in disc diffusion assay. **(F) Enteropan pre-treatment enhanced *P. aeruginosa*’s susceptibility to imipenem**, **as revealed in the broth dilution assay. (G) Increased extracellular protein content in *P. aeruginosa* culture grown in presence of Enteropan.** Protein Unit was calculated as ratio of OD750/OD764 (an indication of protein production per unit of growth). **(H) Reduced intracellular protein content in *P. aeruginosa* grown in presence of Enteropan.** Protein content reported in mg/mL are cell-density neutralized values, wherein OD764 was adjusted to 1.00 prior to cell lysis. Kanamycin employed as a positive control at its sub-MIC level also generated response similar to that of Enteropan from bacterial culture with respect to extracellular and intracellular protein content. DMSO (0.5% v/v) used as ‘vehicle control’ affected neither extracellular nor intracellular protein content.

#### Enteropan had a moderately negative effect on pre-formed biofilm

Though Enteropan’s presence could not compromise *P. aeruginosa*’s ability to form biofilm, when Enteropan was added onto the pre-formed biofilm, it could eradicate the biofilm partly and also reduced the metabolic activity within the biofilm (Figure 3B).

#### Enteropan disturbed nitrogen metabolism in *P. aeruginosa*

Since multiple genes associated with detoxification of reactive nitrogen species were up-regulated (transcriptome data described later), we hypothesized that Enteropan-treated *P. aeruginosa’s* ability to overcome nitrosative stress is compromised. To check this hypothesis, we quantified nitrite concentration in extract-treated *P. aeruginosa* culture supernatant, wherein it was found to have 47.10% higher nitrite concentration as compared to control (Figure 3C). This higher accumulation of nitrite can be taken as an indication of compromised denitrification efficiency, since nitrite is an important intermediate in denitrification pathway ahead of the toxic nitric oxide (Borrero et al., 2017). Nitrosative stress can impact the overall bacterial fitness negatively in multiple ways (Chautrand et al., 2022). Reactive nitrogen species can damage biomolecules like DNA, lipids, and proteins. Resistance to nitrosative stresses is of crucial importance towards the survival of bacteria in the environment as well as inside the host. In gram-negative bacteria, several mechanisms protecting against oxidative and nitrosative stresses are present in the envelope. Excessive nitrosative stress can disturb the envelope homeostasis, and this in fact is reflected in the transcriptome of Enteropan-exposed *P. aeruginosa*, wherein 32 cell envelope (cell wall and lipopolysaccharide) associated genes are getting expressed differently.

#### Enteropan modulated *P. aeruginosa*’s susceptibility to imipenem and augmentin

When Enteropan pre-treated *P. aeruginosa* cells were subsequently challenged with different antibiotics in a disc diffusion assay, these cells exhibited marginal increase in their susceptibility to imipenem; however their susceptibility to augmentin disappeared following Enteropan pre-treatment (Table S5; Figure 3D and 3E). This effect of Enteropan pre-treatment on imipenem susceptibility was also confirmed in liquid culture, wherein Enteropan-pre-treated cells were observed to exhibit upto 21.43% higher susceptibility to imipenem (Figure 3F). Imipenem belongs to the carbapenem class of beta-lactams (Mansour et al., 2021), and carbapenem resistance among *P. aeruginosa* isolates are being viewed as a serious problem (Souza et al., 2021). Since this class of antibiotics are looked as a last resort for treatment of MDR *P. aeruginosa* (Pragasam et al., 2016), resistance modifiers capable of making this bacterium more susceptible to them can be of help in extending the lifespan of these antibiotics by allowing their use at lower doses. However as seen with augmentin in this study, effect of herbals on antibiotic susceptibility of pathogen may not always be favourable.

#### Enteropan alters protein synthesis and secretion in *P. aeruginosa*

Extracellular protein concentration (after nullifying cell density) in *P. aeruginosa* culture supernatant in presence of Enteropan was found to be 1.89-fold higher than that in absence of Enteropan (Figure 3G). Cell density neutralized-intracellular protein concentration of *P. aeruginosa* cells grown in presence of Enteropan was found to be 1.40-fold lower than cells grown in absence of Enteropan (Figure 3H). It seems that Enteropan exerted an inhibitory effect on protein synthesis in *P. aeruginosa*, and promoted protein export. This might have caused even some of the essential proteins to leave the cell. The increased export of proteins by Enteropan-treated cells may be assumed to have originated from overexpression of efflux pump/transport machinery (as suggested by the transcriptome data too described later), and a compromised cell envelope integrity suggested by differential expression of 32 genes involved in cell wall or lipopolysaccharide (LPS) synthesis. Kanamycin, a known inhibitor of protein synthesis in bacteria was employed as a positive control in this assay. Kanamycin belongs to the aminoglycoside group of antibiotics, which at sub-MIC level caused *P. aeruginosa* culture supernatants to have 1.79-fold higher extracellular protein. Such increase in extracellular protein concentration in *P. aeruginosa* exposed to sub-MIC level of kanamycin was also reported by Takahashi et al. (2016).

#### Enteropan treatment causes large scale differential gene expression in *P. aeruginosa*

A whole transcriptome level comparison of the gene expression profile of Enteropan (600 µg/ml)-treated *P. aeruginosa* with that of control revealed a total of 952 genes getting expressed differentially (log fold change ≥ 2 and FDR ≤ 0.001). This amounted to differential expression of 17% of genome, wherein 616 genes were up-regulated (Table S6) and 336 were down-regulated (Table S9). Corresponding volcano plot (Figure S3) is given in supplementary file. A function-wise categorization of all the differentially expressed genes (DEG) is presented in Figure 4. While all DEG pertaining to cell division were down-regulated, and majority of DEG pertaining to translation too; majority of DEG associated with efflux pump/transport were up-regulated. Overexpression of efflux machinery is known to compromise bacterial fitness by causing physiological dysregulation (Sun et al., 2014). Owing to the important physiological roles of efflux pumps in various functions such as intercellular communication, bacterial pathogenicity and virulence, and biofilm formation, expression of majority of them is subject to tight control by different transcriptional regulators. Any mischief with this regulation leading to overexpression of the efflux function may result in leaking of even essential items. An empirical look at the list of DEG suggested that Enteropan attenuated virulence of *P. aeruginosa* by causing dysregulation of metal homeostasis, nitrogen metabolism, transcription, amino acid and protein synthesis, carbon metabolism, motility, efflux, etc. Results of various *in vitro* assays presented in preceding section corroborates well with the transcriptome data.

**Figure 4.**
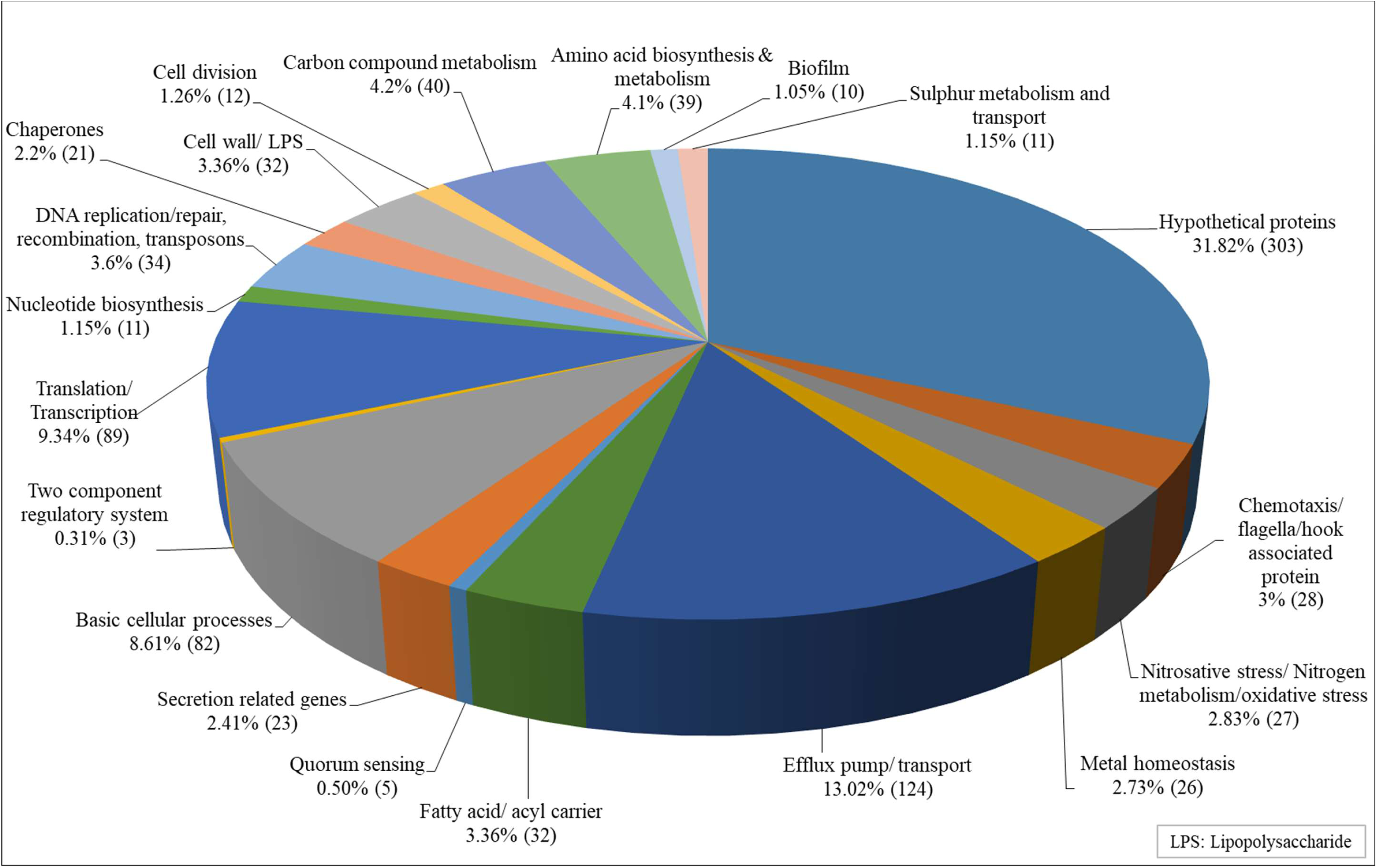
Function-wise categorization of the differentially expressed genes in Enteropan-treated *P. aeruginosa.* Percent values reported are calculated considering the total number of differently expressed genes as 100 percent. Values in parenthesis are number of DEG belonging to that particular category.

#### Network analysis of DEG in Enteropan*-*exposed *P. aeruginosa*

We created Protein-Protein Interaction (PPI) network for up-and down-regulated genes separately. PPI network for up-regulated genes generated through STRING is presented in Figure 5A, which shows 610 nodes connected through 2272 edges with an average node degree of 7.45. Since the number of edges in this PPI network is 2.06-fold higher than expected (1101) with a PPI enrichment p-value <1.0e-16, this network can be said to possess significantly more interactions among the member proteins that what can be expected for a random set of proteins of the identical sample size and degree distribution. Such enrichment can be taken as an indication of the member proteins being at least partially biologically connected. When we arranged all the upregulated DEGs in decreasing order of node degree, 572 nodes were found to have a non-zero score (Table S7), and we selected top 52 genes with a node degree ≥19 for further ranking by different CytoHubba methods. Then we looked for genes which appeared among top-6 ranked candidates by ≥6 cytoHubba methods (Table S8), and 6 such genes were further checked for interactions among themselves by cluster analysis (Figure 5B), whose overexpression can be hypothesized to disturb pathogen physiology. Interaction map of these 6 potential hubs showed them to be strongly networked as the average node degree score was 5. This network possessed 15 edges as against expected (zero) for any such random set of proteins. The PPI network showed 5 of these 6 potential hubs to be part of a single local network cluster (Figure 5B). Cooccurrence analysis showed all of these 6 hubs being absent from humans (Table 3) and hence agonists of these hubs may be expected to target pathogen selectively without interfering with host system functioning.

**Figure 5.**
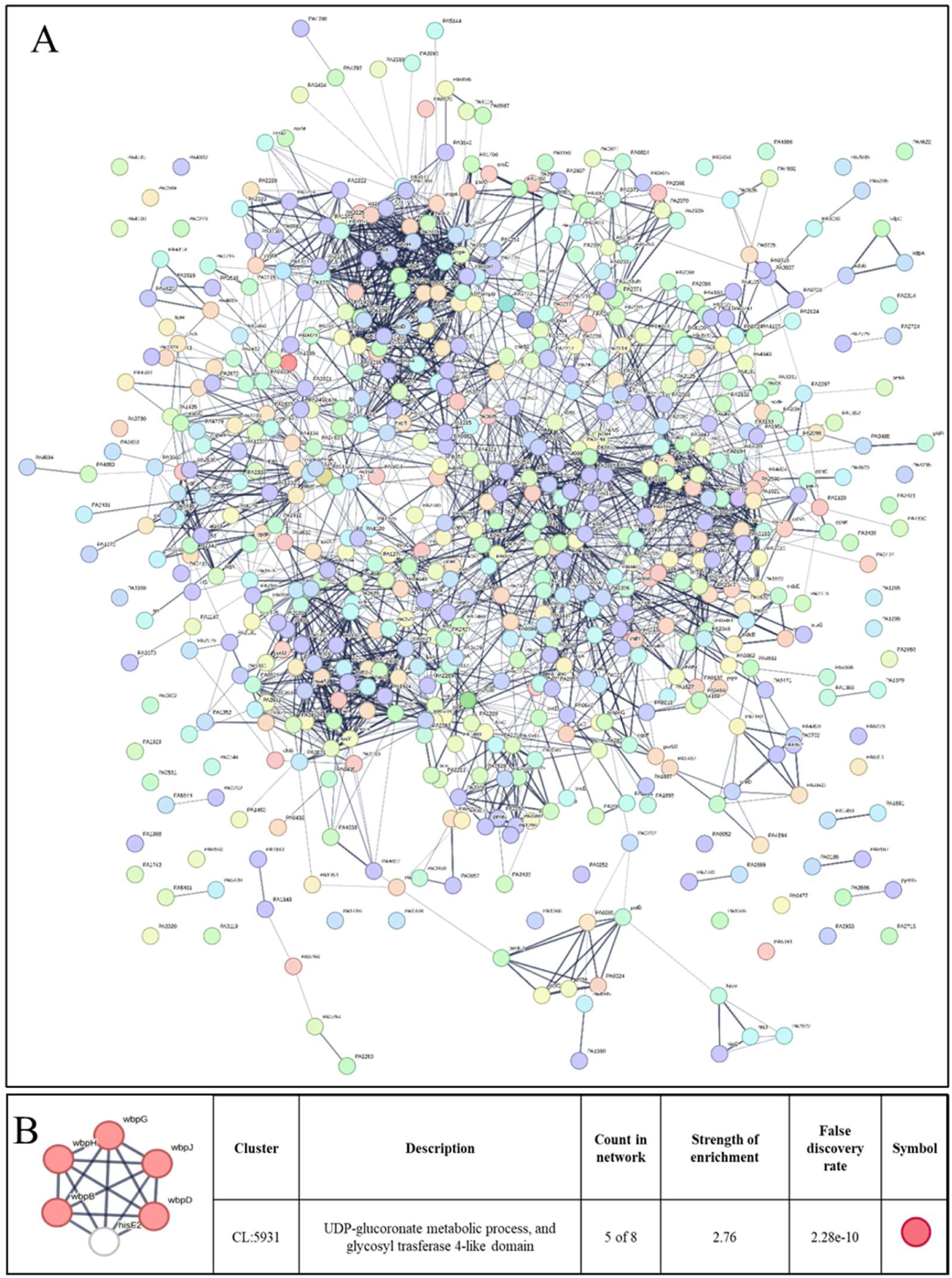
(A) Protein-Protein Interaction (PPI) network of up-regulated genes in Enteropan-exposed *P. aeruginosa*. Edges represents protein-protein associations that are meant to be specific and meaningful, i.e., proteins jointly contribute to a shared function. This does not necessarily mean they are physically binding to each other. Network nodes represents all the proteins produced by a single, protein-coding gene locus. **(B) PPI network of top-ranked genes revealed through cytoHubba among up-regulated DEG in Enteropan-exposed *P. aeruginosa***

**Table 3.** Cooccurrence analysis of genes coding for potential targets in *P. aeruginosa*

| Organism | Potential hubs up-regulated in *P. aeruginosa* |  |  |  |  |  | Potential hubs down-regulated in *P. aeruginosa* |  |  |  |  |  |  |  |  |  |
| --- | --- | --- | --- | --- | --- | --- | --- | --- | --- | --- | --- | --- | --- | --- | --- | --- |
|  | hisF2 | wbpG | wbpD | wbpJ | wbpH | wbpB | rpsL | rplE | rpsD | rpsJ | rplT | rpoA | rpsB | rplN | rplL | rplF |
| *Pseudomonas aeruginosa* | ■ | ■ | ■ | ■ | ■ | ■ | ■ | ■ | ■ | ■ | ■ | ■ | ■ | ■ | ■ | ■ |
| *Homo sapiens* |  |  |  |  |  |  | ■ | ■ |  |  | ■ |  | ■ | ■ | ■ |  |
| *Acinetobacter baumannii* | ■ |  | ■ | ■ | ■ |  | ■ | ■ | ■ | ■ | ■ | ■ | ■ | ■ | ■ | ■ |
| *Enterobacteriaceae* | ■ |  | ■ | ■ |  | ■ | ■ | ■ | ■ | ■ | ■ | ■ | ■ | ■ | ■ | ■ |
| *Salmonella* Serotype typhi | ■ |  |  | ■ |  |  | ■ | ■ | ■ | ■ | ■ | ■ | ■ | ■ | ■ | ■ |
| *Shigella* Spp. | ■ |  | ■ |  |  |  | ■ | ■ | ■ | ■ | ■ | ■ | ■ | ■ | ■ | ■ |
| *Mycobacterium tuberculosis* | ■ |  | ■ |  |  |  | ■ | ■ | ■ | ■ | ■ | ■ | ■ | ■ | ■ | ■ |

The color of the squares denotes, for each gene of interest, the degree of similarity of its best hit in a given STRING genome.
Lowest similarity Highest similarity

PPI network for down-regulated genes is presented in Figure 6A, which shows 327 nodes connected through 3206 edges with an average node degree of 19.6. Since the number of edges in this PPI network is 1.93-fold higher than expected (1658) with a PPI enrichment p-value <1.0e-16, this network can be said to possess significantly more interactions among the member proteins that what can be expected for a random set of proteins of the identical sample size and degree distribution. Such enrichment can be taken as an indication of the member proteins being at least partially biologically connected. When we arranged all the down-regulated DEGs in decreasing order of node degree, 298 nodes were found to have a non-zero score, and we selected top 50 genes with a node degree ≥53 (Table S10) for further ranking by different cytoHubba methods. Then we looked for genes which appear among top-10 ranked candidates by ≥6 cytoHubba methods, and 10 such genes (Table S11) were identified as potential hubs, whose down-regulation can be hypothesized to attenuate *P. aeruginosa* virulence. Interaction map of these 10 potential hubs (Figure 6B) showed them to be strongly networked as the average node degree score was 9. This network possessed 45 edges as against expected (fifteen) for any such random set of proteins. The PPI network showed these 10 genes to be distributed among six different local network clusters, whose strength score ranged from 1.97-2.2 (Figure 6B). Cooccurrence analysis (Table 3) of these ten hub proteins indicated four (rpsD, rpsJ, rpoA, rplF) of them to be absent from humans, and hence they can be said to possess high targetability with respect to discovery of new antibiotics satisfying the criteria of selective toxicity. Since all the 10 predicted hubs are indicated by cooccurrence analysis to be present in other important bacterial pathogens too, antagonists of these proteins are likely to be useful as broad-spectrum antibiotics.

**Figure 6.**
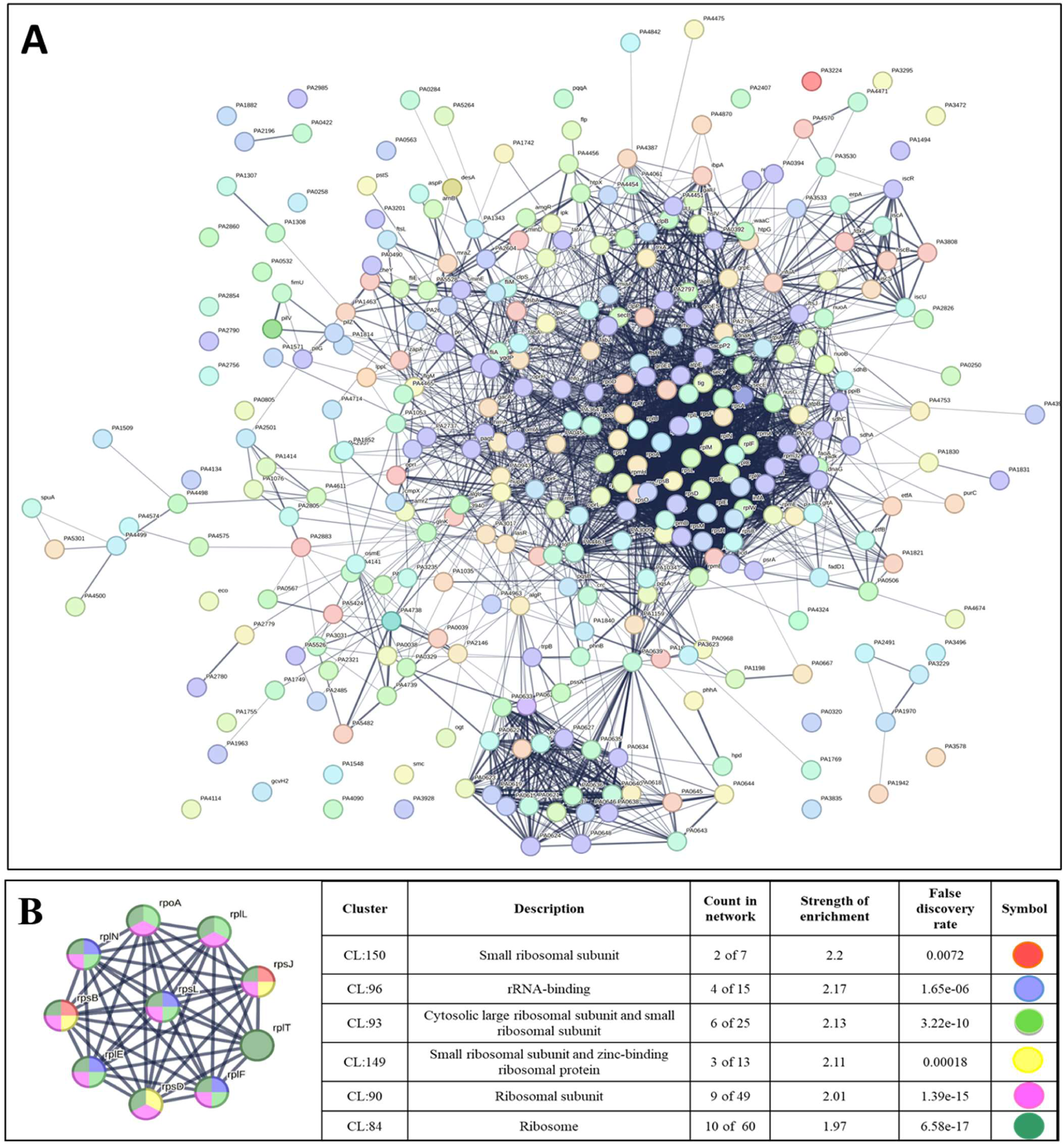
**(A) Protein-Protein Interaction (PPI) network of down-regulated genes in Enteropan-exposed *P. aeruginosa* (B) PPI network of top-ranked genes revealed through cytoHubba among down-regulated DEG in Enteropan-exposed *P. aeruginosa***

#### Target validation through RT-PCR

From the 10 identified hubs among the down-regulated genes, we selected five (*rpoA, tig, rpsB, rpsL, rpsJ*) for further validation through RT-PCR. From the ten genes shown in Figure 6B, four (*rpoA, rpsB, rpsL, rpsJ*) passing the dual criteria of node degree ≥70 (Table S10) and been part of ≥3 clusters were selected for RT-PCR. Though rpsD (node degree 70) also passed these dual criteria, since already three rps genes were selected for PCR, we preferred rpoA (node degree 75) over it. Additionally, we included one gene (*tig*; 3.73-fold↓; node degree 70) for PCR validation, though it was not among the identified hubs, because tig is a trigger factor involved in protein export, and we did observe a heavy increase in extracellular protein content in Enteropan-exposed *P. aeruginosa.* PCR did confirm down regulation of all the selected five genes in Enteropan-exposed *P. aeruginosa* (Figure 7), and thus they can be considered as potential antibacterial targets worthy of attention by drug discovery programmes.

**Figure 7.**
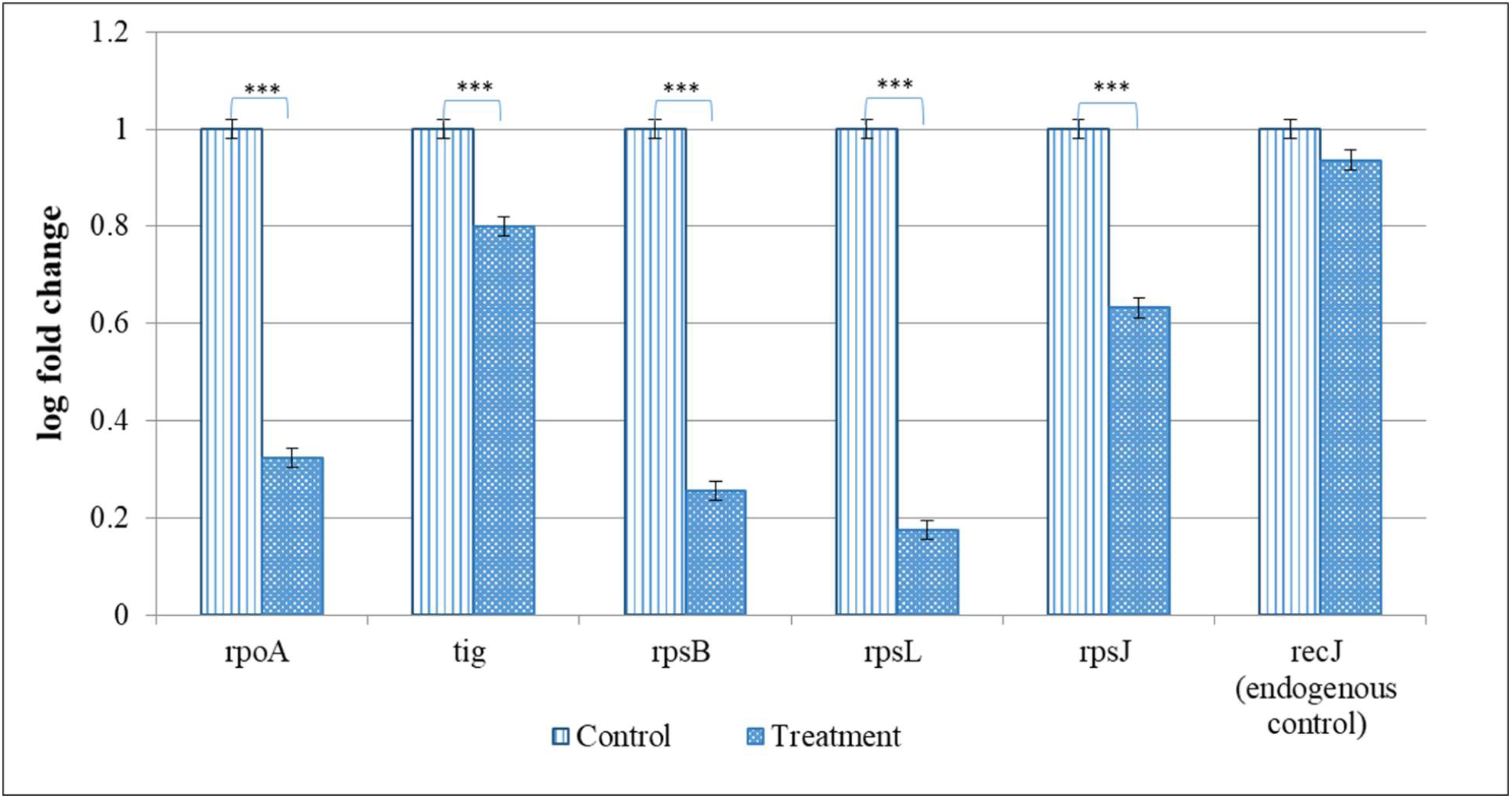
Confirmation of differential expression of selected genes in Enteropan-treated *P. aeruginosa* through RT-PCR. *recJ* selected as an endogenous control was not expressed differently (FDR 1) between control and experimental bacterial cultures. ***p≤0.001

## Conclusion

The polyherbal formulation Enteropan was found to have virulence-attenuating effect against an important gram-negative bacterial pathogen *P. aeruginosa*, without affecting its growth heavily. As can be expected from any multicomponent polyherbal formulation, Enteropan also exerted multiplicity of targets against the test pathogen. A large fraction of the bacterial genome was expressed differently under influence of this anti-pathogenic formulation, which corroborated well with the altered phenotypic traits in extract-exposed bacterial culture. Major mechanisms revealed through various *in vitro/ in vivo* assays and transcriptome analysis through which Enteropan exerted its anti-virulence activity were found to be generation of nitrosative stress, oxidative stress, quorum modulation, disturbance of protein homeostasis and metal homeostasis. A wholistic summary depicting the mechanistic details associated with the anti-pathogenic potential of Enteropan against *P. aeruginosa* is presented in Figure 8, with particular attention on Enteropan’s effect on quorum sensing machinery and virulence regulators of this notorious pathogen. Our results validate the anti-pathogenic potential of Enteropan, and also the concept of polyherbalism and its relevance in combating antimicrobial resistance.

**Figure 8.**
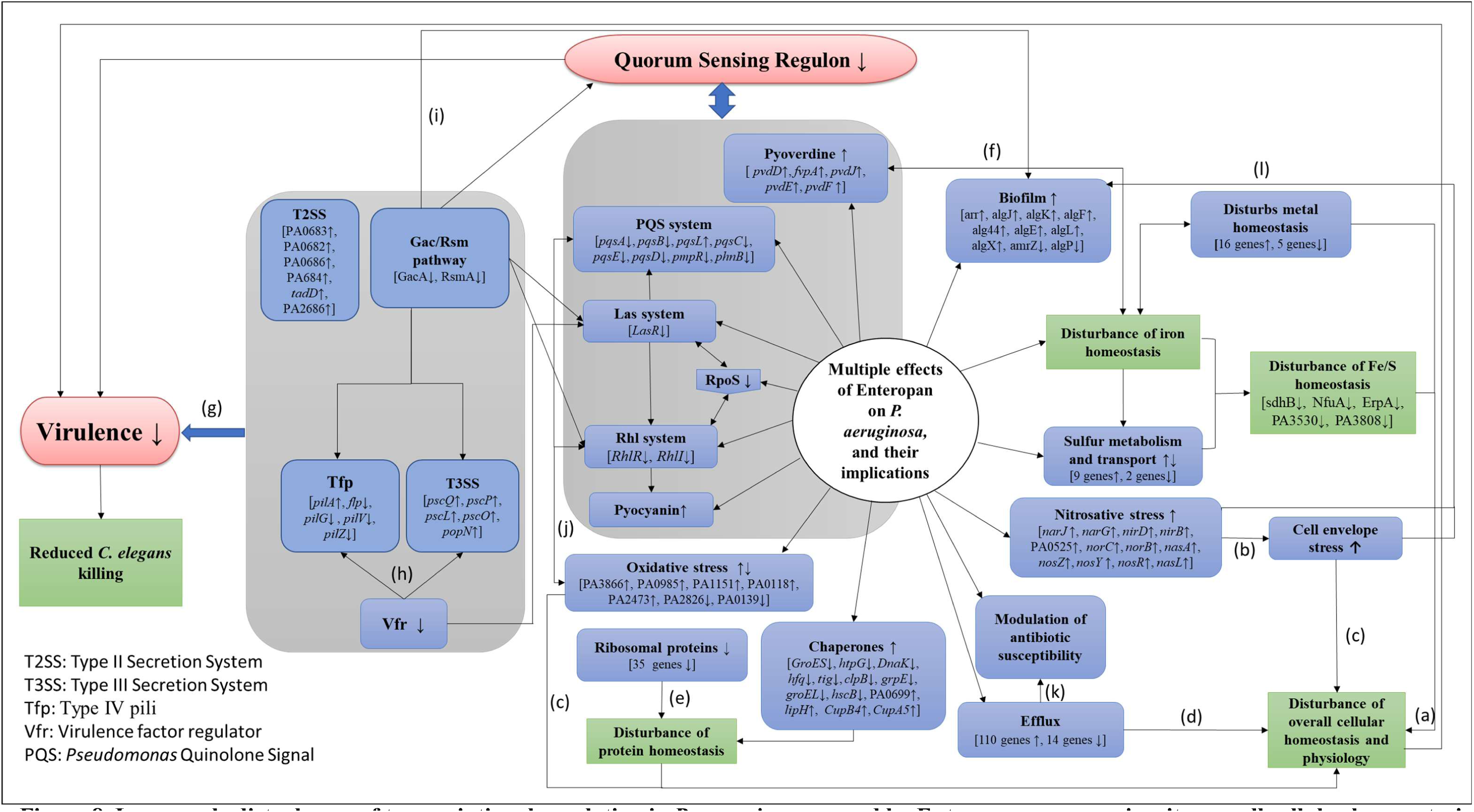
Large scale disturbance of transcriptional regulation in *P. aeruginosa* caused by Enteropan compromises its overall cellular homeostasis and virulence. This figure presents a wholistic summary of multiple effects exerted by Enteropan against *P. aeruginosa.* Various cellular, physiological, and virulence-associated traits of *P. aeruginosa* expressed differently under the influence of Enteropan are depicted. The genes shown with an up or down arrow are those getting differentially expressed with a log fold change of ≥1.5 and FDR≤0.05. The **Gac/Rsm pathway** inversely regulates the expression of virulence factors (T3SS, Tfp, exopolysaccharides) associated with acute and chronic disease (Sultan et al., 2021). **T2SS** is responsible for secreting many secretory proteins like alkaline phosphatase, lipase, exotoxin A, phospholipase, and proteases. **T3SS** is largely involved in secretion of virulence determinants associated with acute infection. *Vfr* is a global regulator of virulence gene expression, which allows coordinated production of related virulence functions (Tfp, T3SS) necessary for adherence to an intoxication of host cells. **LasR** is a transcriptional activator of multiple virulence-associated genes in *P. aeruginosa.* It represents a central checkpoint, with the highest degree of interconnection in the network. **RpoS**, the stationary phase sigma factor, influences the expression of more than one-third of all the quorum-regulated genes. It is a central regulator of many stationary phase-inducible genes and a master stress-response regulator under a variety of stress conditions (Murakami et al., 2015). **Tfp**, a major surface adhesin mechanochemically regulates virulence factors in *P. aeruginosa* (Persat et al., 2015). The *Rhl* system is a quorum sensing system acquired by *P. aeruginosa* through lateral gene transfer. **PQS** is an essential mediator of the shaping of the population structure of *P. aeruginosa* and of its response to and survival in stress conditions. [^a^Myriam et al., 2022; ^b^Chautrand et al., 2022; ^c^Mitchell and Silhavy, 2019; ^d^Hajiagha and Kafil, 2023; ^e^Deuerling et al., 2019; ^f^Hamza et al., 2023; ^g^Coggan and Wolfgang, 2012; ^h^Marsden et al., 2016; ^i^Schuster and Greenberg, 2007, ^j^Häussler and Becker, 2008, ^k^Xu et al., 2020; ^l^Fazeli-Nasab et al., 2022]

## Acknowledgement

Authors thank Nirma Education and Research Foundation (NERF), Ahmedabad for infrastructural support; Virupakshi Soppina (IIT-Gn) for providing worms. SP and GG acknowledge fellowship from Gujarat government under their SHODH scheme. NT acknowledges fellowship from Nirma University.

