## Supplementary File for "Deciphering the molecular mechanisms underlying anti-pathogenic potential of a polyherbal formulation Enteropan^®^ against multi-drug resistant *Pseudomonas aeruginosa*"

^#^Contributed equally

**Table S1. Antibiogram of *P. aeruginosa* generated through Kirby-Bauer Disc Diffusion assay**

| **Antibiotic** | **Concentration (µg/disc)** | **Interpretation** |
| --- | --- | --- |
| Imipenem (IPM) | 10 | Sensitive |
| Ciprofloxacin (CIP) | 5 | Sensitive |
| Tobramycin (TOB) | 10 | Sensitive |
| Moxifloxacin (MO) | 5 | Sensitive |
| Ofloxacin (OF) | 5 | Sensitive |
| Sparfloxacin (SPX) | 5 | Sensitive |
| Levofloxacin (LE) | 5 | Sensitive |
| Norfloxacin (NX) | 10 | Sensitive |
| Co-Trimoxazole (COT) | 25 | Resistant |
| Colistin (CL) | 10 | Sensitive |
| Nalidixic acid (NA) | 30 | Sensitive |
| Augmentin (AMC) | 30 | Resistant |
| Kanamycin (K) | 30 | Intermediate |
| Gatifloxacin (GAT) | 5 | Sensitive |
| Gentamicin (GEN) | 10 | Sensitive |
| Amikacin (AK) | 30 | Sensitive |
| Streptomycin (S) | 25 | Resistant |
| Ceftriaxone (CTR) | 30 | Sensitive |
| Cefpodoxime (CPD) | 10 | Sensitive |
| Ticarcillin (TI) | 75 | Sensitive |

Antibiotic susceptibility profile of the organism was generated using the antibiotic discs- Icosa GI Minus (HiMedia, Mumbai) through disc diffusion assay on cation-adjusted Mueller-Hinton agar (HiMedia) as per CLSI guidelines (https://doi.org/10.1177/001857870403900608). The zones of inhibition were measured and the interpretation (S- sensitive, I - intermediate, R - resistant) was drawn as per zone size interpretative chart provided by the manufacturer

**Table S2. Quantification of extracted RNA**

| **Sr. No.** | **Sample Name** | **ng/μl** | **260/280** | **260/230** | **Quant (ng/µl)** | **RIN Value** | **QC Remark** |
| --- | --- | --- | --- | --- | --- | --- | --- |
| 1 | Control | 3744.4 | 2.15 | 2.22 | 1984 | 7.6 | Pass |
| 2 | Experimental | 2523.3 | 1.94 | 1.75 | 976 | 7.3 | Pass |

**Table S3. Library preparation and quality control**

| **Sr. No.** | **Sample Name** | **ng/μl** | **Quant (ng/µl)** | **Index** | **QC Remark** |
| --- | --- | --- | --- | --- | --- |
| 1 | Control | 4.58 | 271 | 52 | Pass |
| 2 | Experimental | 0.912 | 281 | 53 | Pass |

**Table S4. Temperature profile for the RT-PCR assay**

| **Temperature** (^o^C) | **Time** (Seconds) |
| --- | --- |
| PCR stage (45 Cycles) | |
| 95 | 15 |
| 59 | 60 |
| Melt curve stage | |
| 95 | 15 |
| 60 | 60 |
| 95 | 15 |

**Table S5. Enteropan pre-treatment modulated bacterial susceptibility to some antibiotics**

| **Antibiotic** | **Symbol** | **Concentration** (µg/disc) | **Zone of inhibition**  (mm) | | **% Difference** |
| --- | --- | --- | --- | --- | --- |
|  |  |  | **Control**  (Mean ± SD) | **Experimental**  (Mean ± SD) |  |
| Imipenem | IMP | 10 | 25.6±2 | 29.6±1.5 | 15.58* ± 7.9 |
| Ciprofloxacin | CIP | 5 | 40±2.6 | 39±3.5 | Not significant |
| Tobramycin | TOB | 10 | 31±3 | 29±1.5 |  |
| Moxifloxacin | MO | 5 | 30±0.5 | 31±2 |  |
| Ofloxacin | OF | 5 | 32±2 | 30±2.3 |  |
| Sparfloxacin | SPX | 5 | 29±0.5 | 30±0.5 |  |
| Levofloxacin | LE | 5 | 37±2.5 | 37±2.5 |  |
| Norfloxacin | NX | 10 | 33±1.5 | 34±1 |  |
| Co-Trimoxazole | COT | 25 | 0 | 0 |  |
| Colistin | CL | 10 | 20±0.5 | 20±0.5 |  |
| Nalidixic acid | NA | 30 | 15±0.5 | 17±1.7 |  |
| Augmentin | AMC | 30 | 10 | 0 | 100*** ± 0 |
| Kanamycin | K | 30 | 12±1 | 13±2 | Not significant |
| Gatifloxacin | GAT | 5 | 31±1 | 32±1 |  |
| Gentamicin | GEN | 10 | 22±2 | 21±1 |  |
| Amikacin | AK | 30 | 24.5±1.5 | 23±5 |  |
| Streptomycin | S | 25 | 0 | 0 |  |
| Ceftriaxone | CTR | 30 | 21±0.5 | 23±1 |  |
| Cefpodoxime | CPD | 10 | 22±3 | 22±4 |  |
| Ticarcillin | TI | 15 | 19 | 20 |  |

Antibiotic susceptibility profile of the bacterium was generated using the antibiotic discs- Icosa G-I Minus (HiMedia, Mumbai), through disc diffusion assay performed as per CLSI guidelines. The zones of inhibition were measured and the interpretation (S - sensitive, I - intermediate, R - resistant) was drawn as per zone size interpretative chart provided by the manufacturer. *p<0.05

**Table S6. List of Up-regulated genes in Enteropan exposed *P. aeruginosa* satisfying the dual criteria of log fold change ≥2 and FDR≤0.001**

| **Sr. No.** | **Gene ID** | **Symbol** | **Product name** | **Log FC** | **FDR** |
| --- | --- | --- | --- | --- | --- |
| 1 | PA2399 | *pvdD* | pyoverdine synthetase D | 14.13 | 3.68311E-15 |
| 2 | PA1094 | *fliD* | B-type flagellar hook-associated protein | 13.98 | 5.10836E-15 |
| 3 | PA1091 | *fgtA* | flagellar glycosyl transferase FgtA | 13.75 | 1.25225E-14 |
| 4 | PA3160 | *Wzz* | O-antigen chain length regulator | 13.75 | 1.25225E-14 |
| 5 | PA3145 | *wbpL* | glycosyltransferase WbpL | 13.56 | 3.78409E-14 |
| 6 | PA3153 | *Wzx* | O-antigen translocase | 13.39 | 9.85022E-14 |
| 7 | PA3487 | *pldA* | phospholipase D | 13.32 | 1.32823E-13 |
| 8 | PA2732 | NA | hypothetical protein | 13.26 | 1.56091E-13 |
| 9 | PA2398 | *fpvA* | ferripyoverdine receptor | 13.26 | 1.56091E-13 |
| 10 | PA1095 | NA | B-type flagellar protein FliS | 13.10 | 4.2517E-13 |
| 11 | PA3154 | *Wzy* | B-band O-antigen polymerase | 13.05 | 5.42286E-13 |
| 12 | PA3157 | NA | Acetyltransferase | 12.92 | 1.05141E-12 |
| 13 | PA3488 | NA | hypothetical protein | 12.91 | 1.09728E-12 |
| 14 | PA1087 | *flgL* | flagellar hook-associated protein FlgL | 12.81 | 1.80849E-12 |
| 15 | PA3866 | NA | pyocin protein | 12.78 | 2.0428E-12 |
| 16 | PA2735 | NA | restriction-modification system protein | 12.75 | 2.39147E-12 |
| 17 | PA3159 | *wbpA* | UDP-N-acetyl-d-glucosamine 6-dehydrogenase | 12.74 | 2.40102E-12 |
| 18 | PA1428a | NA | hypothetical protein | 12.53 | 8.39067E-12 |
| 19 | PA2818 | *Arr* | aminoglycoside response regulator | 12.53 | 8.39067E-12 |
| 20 | PA2119 | NA | alcohol dehydrogenase | 12.42 | 1.42681E-11 |
| 21 | PA0561 | NA | hypothetical protein | 12.36 | 2.02829E-11 |
| 22 | PA0985 | *pyoS5* | pyocin S5 | 12.28 | 3.36661E-11 |
| 23 | PA0498 | NA | hypothetical protein | 12.26 | 3.74703E-11 |
| 24 | PA0826 | NA | hypothetical protein | 12.21 | 4.64138E-11 |
| 25 | PA0821 | NA | hypothetical protein | 12.21 | 4.64138E-11 |
| 26 | PA1939 | NA | hypothetical protein | 12.19 | 5.11036E-11 |
| 27 | PA1888 | NA | hypothetical protein | 12.06 | 1.12609E-10 |
| 28 | PA3868 | NA | hypothetical protein | 12.05 | 1.12609E-10 |
| 29 | PA3506 | NA | hypothetical protein | 11.98 | 1.80412E-10 |
| 30 | PA0497 | NA | hypothetical protein | 11.96 | 1.94271E-10 |
| 31 | PA3147 | *wbpJ* | glycosyl transferase WbpJ | 11.91 | 2.42097E-10 |
| 32 | PA4797 | NA | Transposase | 11.91 | 2.42097E-10 |
| 33 | PA3993 | NA | Transposase | 11.88 | 2.53006E-10 |
| 34 | PA2319 | NA | Transposase | 11.85 | 2.80745E-10 |
| 35 | PA0716 | NA | hypothetical protein | 11.80 | 3.72012E-10 |
| 36 | PA0978 | NA | hypothetical protein | 11.78 | 3.8445E-10 |
| 37 | PA3500 | NA | hypothetical protein | 11.78 | 3.87946E-10 |
| 38 | PA2219 | *opdE* | transcriptional regulator OpdE | 11.76 | 4.22853E-10 |
| 39 | PA2734 | NA | hypothetical protein | 11.76 | 4.22853E-10 |
| **Sr. No.** | **Gene ID** | **Symbol** | **Product name** | **Log FC** | **FDR** |
| 40 | PA3146 | *wbpK* | NAD-dependent epimerase/dehydratase | 11.76 | 4.22853E-10 |
| 41 | PA2690 | NA | Transposase | 11.71 | 2.33324E-10 |
| 42 | PA1370 | NA | hypothetical protein | 11.70 | 2.41787E-10 |
| 43 | PA3504 | NA | aldehyde dehydrogenase | 11.68 | 2.44885E-10 |
| 44 | PA2101 | NA | hypothetical protein | 11.68 | 2.44885E-10 |
| 45 | PA0981 | NA | hypothetical protein | 11.67 | 2.44885E-10 |
| 46 | PA0188 | NA | hypothetical protein | 11.67 | 2.47595E-10 |
| 47 | PA3434 | NA | Transposase | 11.67 | 2.50443E-10 |
| 48 | PA1366 | NA | hypothetical protein | 11.64 | 2.75999E-10 |
| 49 | PA1372 | NA | hypothetical protein | 11.63 | 2.92294E-10 |
| 50 | PA2073 | NA | transporter membrane subunit | 11.61 | 3.15239E-10 |
| 51 | PA0445 | NA | Transposase | 11.58 | 3.73749E-10 |
| 52 | PA2459 | NA | hypothetical protein | 11.58 | 3.79803E-10 |
| 53 | PA1093 | NA | hypothetical protein | 11.57 | 3.8445E-10 |
| 54 | PA1368 | NA | hypothetical protein | 11.56 | 3.87946E-10 |
| 55 | PA3497 | NA | hypothetical protein | 11.53 | 4.25297E-10 |
| 56 | PA3549 | *algJ* | alginate o-acetylase AlgJ | 11.53 | 4.25297E-10 |
| 57 | PA2100 | NA | transcriptional regulator | 11.52 | 4.62825E-10 |
| 58 | PA2099 | NA | short-chain dehydrogenase | 11.48 | 5.95777E-10 |
| 59 | PA2091 | NA | hypothetical protein | 11.46 | 6.50932E-10 |
| 60 | PA2566 | NA | hypothetical protein | 11.44 | 7.44011E-10 |
| 61 | PA2220 | NA | transcriptional regulator | 11.38 | 1.00798E-09 |
| 62 | PA3158 | *wbpB* | UDP-N-acetyl-2-amino-2-deoxy-D-glucuronate oxidase | 11.38 | 1.00798E-09 |
| 63 | PA0715 | NA | hypothetical protein | 11.32 | 1.39194E-09 |
| 64 | PA0982 | NA | hypothetical protein | 11.31 | 1.48398E-09 |
| 65 | PA1938 | NA | hypothetical protein | 11.26 | 1.96615E-09 |
| 66 | PA2221 | NA | hypothetical protein | 11.25 | 2.0238E-09 |
| 67 | PA0820 | NA | hypothetical protein | 11.24 | 2.08366E-09 |
| 68 | PA2228 | NA | hypothetical protein | 11.22 | 2.41626E-09 |
| 69 | PA0187 | NA | hypothetical protein | 11.13 | 3.9928E-09 |
| 70 | PA3867 | NA | DNA invertase | 11.13 | 4.12502E-09 |
| 71 | PA3513 | NA | hypothetical protein | 11.12 | 4.2233E-09 |
| 72 | PA1351 | NA | ECF subfamily sigma-70 factor | 11.11 | 4.28719E-09 |
| 73 | PA1088 | NA | hypothetical protein | 11.07 | 5.45271E-09 |
| 74 | PA3149 | *wbpH* | glycosyltransferase WbpH | 11.07 | 5.64638E-09 |
| 75 | PA1380 | NA | transcriptional regulator | 11.06 | 5.79808E-09 |
| 76 | PA3155 | *wbpE* | UDP-2-acetamido-2-deoxy-3-oxo-D-glucuronate aminotransferase | 11.06 | 5.79808E-09 |
| 77 | PA2730 | NA | hypothetical protein | 11.02 | 7.11987E-09 |
| 78 | PA3151 | *hisF2* | imidazole glycerol phosphate synthase subunit HisF | 11.00 | 8.02025E-09 |
| 79 | PA3150 | *wbpG* | LPS biosynthesis protein WbpG | 10.97 | 9.4834E-09 |
| **Sr. No.** | **Gene ID** | **Symbol** | **Product name** | **Log FC** | **FDR** |
| 80 | PA3152 | *hisH2* | imidazole glycerol phosphate synthase subunit HisH | 10.88 | 1.61662E-08 |
| 81 | PA3510 | NA | hypothetical protein | 10.88 | 1.61662E-08 |
| 82 | PA2104 | NA | cysteine synthase | 10.86 | 1.81848E-08 |
| 83 | PA0824 | NA | hypothetical protein | 10.86 | 1.81848E-08 |
| 84 | PA3511 | NA | short-chain dehydrogenase | 10.83 | 2.06736E-08 |
| 85 | PA1379 | NA | short-chain dehydrogenase | 10.83 | 2.121E-08 |
| 86 | PA3514 | NA | ABC transporter ATP-binding protein | 10.82 | 2.20676E-08 |
| 87 | PA0984 | NA | colicin immunity protein | 10.80 | 2.42855E-08 |
| 88 | PA2733 | NA | hypothetical protein | 10.79 | 2.5397E-08 |
| 89 | PA2461 | NA | hypothetical protein | 10.75 | 3.192E-08 |
| 90 | PA2103 | NA | molybdopterin biosynthesis protein MoeB | 10.74 | 3.32203E-08 |
| 91 | PA3142 | NA | hypothetical protein | 10.73 | 3.41347E-08 |
| 92 | PA3143 | NA | hypothetical protein | 10.72 | 3.57971E-08 |
| 93 | PA2223 | NA | hypothetical protein | 10.71 | 3.75532E-08 |
| 94 | PA3148 | *wbpI* | UDP-2,3-diacetamido-2,3-dideoxy-D-glucuronate 2-epimeras | 10.68 | 4.62232E-08 |
| 95 | PA2460 | NA | hypothetical protein | 10.68 | 4.62232E-08 |
| 96 | PA2106 | NA | hypothetical protein | 10.64 | 5.6473E-08 |
| 97 | PA2736 | NA | hypothetical protein | 10.63 | 5.94262E-08 |
| 98 | PA3498 | NA | Oxidoreductase | 10.61 | 6.85628E-08 |
| 99 | PA3381 | NA | transcriptional regulator | 10.58 | 8.06539E-08 |
| 100 | PA3512 | NA | ABC transporter permease | 10.58 | 8.06539E-08 |
| 101 | PA2218 | NA | hypothetical protein | 10.57 | 8.42948E-08 |
| 102 | PA1887 | NA | hypothetical protein | 10.56 | 8.7959E-08 |
| 103 | PA1937 | NA | hypothetical protein | 10.55 | 8.7959E-08 |
| 104 | PA2564 | NA | trans-aconitate 2-methyltransferase | 10.47 | 1.45077E-07 |
| 105 | PA2387 | *fpvI* | RNA polymerase sigma factor | 10.45 | 1.52147E-07 |
| 106 | PA1378 | NA | hypothetical protein | 10.43 | 1.70321E-07 |
| 107 | PA3865a | NA | hypothetical protein | 10.42 | 1.78934E-07 |
| 108 | PA1151 | *imm2* | pyocin-S2 immunity protein | 10.38 | 2.22683E-07 |
| 109 | PA0522 | NA | hypothetical protein | 10.38 | 2.22683E-07 |
| 110 | PA0979 | NA | hypothetical protein | 10.34 | 2.70275E-07 |
| 111 | PA0825 | NA | hypothetical protein | 10.31 | 3.24308E-07 |
| 112 | PA2105 | NA | Acetyltransferase | 10.28 | 3.67943E-07 |
| 113 | PA2427 | NA | hypothetical protein | 10.24 | 4.7244E-07 |
| 114 | PA1089 | NA | hypothetical protein | 10.21 | 5.30195E-07 |
| 115 | PA3501 | NA | hypothetical protein | 10.21 | 5.30195E-07 |
| 116 | PA4823 | NA | hypothetical protein | 10.15 | 7.3674E-07 |
| 117 | PA0986 | NA | hypothetical protein | 10.13 | 8.31928E-07 |
| 118 | PA0983 | NA | hypothetical protein | 10.12 | 8.31928E-07 |
| 119 | PA1090 | NA | hypothetical protein | 10.10 | 9.57461E-07 |
| **Sr. No.** | **Gene ID** | **Symbol** | **Product name** | **Log FC** | **FDR** |
| 120 | PA0823 | NA | hypothetical protein | 10.10 | 9.57461E-07 |
| 121 | PA0209 | NA | 2-(5''-triphosphoribosyl)-3'-dephosphocoenzyme-A synthase | 10.08 | 1.02062E-06 |
| 122 | PA3156 | *wbpD* | UDP-2-acetamido-3-amino-2,3-dideoxy-D-glucuronate N-acetyltransferase | 10.07 | 1.07604E-06 |
| 123 | PA2516 | *xylZ* | toluate 1,2-dioxygenase electron transfer subunit | 10.06 | 1.07604E-06 |
| 124 | PA3508 | NA | transcriptional regulator | 10.03 | 1.33268E-06 |
| 125 | PA0213 | NA | phosphoribosyl-dephospho-CoA transferase | 10.01 | 1.43263E-06 |
| 126 | PA3869 | NA | hypothetical protein | 10.00 | 1.53914E-06 |
| 127 | PA1981 | NA | hypothetical protein | 9.99 | 1.6561E-06 |
| 128 | PA3507 | NA | short-chain dehydrogenase | 9.96 | 1.93577E-06 |
| 129 | PA3505 | NA | L-aspartate dehydrogenase | 9.95 | 2.0662E-06 |
| 130 | PA3502 | NA | hypothetical protein | 9.91 | 2.41059E-06 |
| 131 | PA5417 | *soxD* | sarcosine oxidase subunit delta | 9.77 | 4.98351E-06 |
| 132 | PA2102 | NA | hypothetical protein | 9.72 | 5.77943E-06 |
| 133 | PA3509 | NA | Hydrolase | 9.67 | 7.5136E-06 |
| 134 | PA2161 | NA | hypothetical protein | 9.56 | 1.26957E-05 |
| 135 | PA3380 | NA | hypothetical protein | 9.56 | 1.26957E-05 |
| 136 | PA1369 | NA | hypothetical protein | 9.54 | 1.38061E-05 |
| 137 | PA1371 | NA | hypothetical protein | 9.52 | 1.51726E-05 |
| 138 | PA2565 | NA | hypothetical protein | 9.51 | 1.51726E-05 |
| 139 | PA2224 | NA | hypothetical protein | 9.49 | 1.82008E-05 |
| 140 | PA1702 | NA | hypothetical protein | 9.48 | 1.82008E-05 |
| 141 | PA0210 | *mdcC* | malonate decarboxylase acyl carrier protein | 9.42 | 2.45359E-05 |
| 142 | PA3499 | NA | hypothetical protein | 9.42 | 2.45359E-05 |
| 143 | PA2225 | NA | hypothetical protein | 9.40 | 2.70635E-05 |
| 144 | PA2098 | NA | Esterase | 9.37 | 2.98572E-05 |
| 145 | PA1635 | *kdpC* | potassium-transporting ATPase subunit C | 9.36 | 3.28091E-05 |
| 146 | PA2227 | *vqsM* | HTH-type transcriptional regulator VqsM | 9.30 | 4.51481E-05 |
| 147 | PA2222 | NA | hypothetical protein | 8.84 | 0.0002 |
| 148 | PA1700 | NA | hypothetical protein | 8.62 | 0.0007 |
| 149 | PA2368 | NA | hypothetical protein | 8.58 | 0.0008 |
| 150 | PA1849 | NA | hypothetical protein | 8.41 | 0.0004 |
| 151 | PA2226 | NA | hypothetical protein | 8.30 | 0.0006 |
| 152 | PA2457 | NA | hypothetical protein | 7.58 | 1.3333E-11 |
| 153 | PA4894 | NA | hypothetical protein | 7.40 | 1.25832E-09 |
| 154 | PA4860 | NA | ABC transporter permease | 7.39 | 1.27621E-09 |
| 155 | PA0695 | NA | hypothetical protein | 7.19 | 4.17225E-09 |
| 156 | PA2400 | *pvdJ* | pyoverdine biosynthesis protein PvdJ | 6.95 | 8.39067E-12 |
| **Sr. No.** | **Gene ID** | **Symbol** | **Product name** | **Log FC** | **FDR** |
| 157 | PA0697 | NA | hypothetical protein | 6.83 | 2.97214E-08 |
| 158 | PA3774 | NA | acetylpolyamine aminohydrolase | 6.72 | 5.26668E-08 |
| 159 | PA1352 | NA | hypothetical protein | 6.68 | 6.67308E-08 |
| 160 | PA2472 | NA | major facilitator superfamily transporter | 6.65 | 1.74802E-09 |
| 161 | PA5401 | NA | hypothetical protein | 6.61 | 9.7707E-08 |
| 162 | PA1698 | *popN* | type III secretion outer membrane protein PopN | 6.40 | 2.96117E-07 |
| 163 | PA0021 |  | hypothetical protein | 6.27 | 1.45581E-08 |
| 164 | PA4525 | *pilA* | type 4 fimbrial protein PilA | 6.26 | 3.21053E-10 |
| 165 | PA4088 |  | Aminotransferase | 6.21 | 2.09682E-08 |
| 166 | PA1275 | *cobD* | cobalamin biosynthesis protein CobD | 6.18 | 9.57461E-07 |
| 167 | PA3406 | *hasD* | transporter HasD | 6.14 | 4.25628E-09 |
| 168 | PA2397 | *pvdE* | pyoverdine biosynthesis protein PvdE | 6.10 | 1.11433E-09 |
| 169 | PA4097 | NA | alcohol dehydrogenase | 6.04 | 1.87569E-06 |
| 170 | PA0440 | NA | Oxidoreductase | 6.00 | 9.82392E-09 |
| 171 | PA5392 | NA | hypothetical protein | 5.79 | 3.14228E-08 |
| 172 | PA2336 | NA | hypothetical protein | 5.76 | 3.41347E-08 |
| 173 | PA3379 | NA | carbon-phosphorus lyase complex subunit | 5.73 | 8.60461E-06 |
| 174 | PA4152 | NA | branched-chain alpha-keto acid dehydrogenase subunit E2 | 5.72 | 2.75675E-07 |
| 175 | PA2724 | NA | hypothetical protein | 5.69 | 3.38903E-07 |
| 176 | PA0683 | NA | type II secretion system protein | 5.67 | 3.78438E-07 |
| 177 | PA2090 | NA | hypothetical protein | 5.64 | 1.31979E-05 |
| 178 | PA3375 | NA | ABC transporter ATP-binding protein | 5.60 | 1.59524E-05 |
| 179 | PA2092 | NA | major facilitator superfamily transporter | 5.53 | 1.24899E-07 |
| 180 | PA1265 | NA | hypothetical protein | 5.53 | 1.28024E-07 |
| 181 | PA0112 | NA | hypothetical protein | 5.50 | 1.48874E-07 |
| 182 | PA0881 | NA | hypothetical protein | 5.46 | 1.8238E-07 |
| 183 | PA0514 | *nirL* | heme d1 biosynthesis protein NirL | 5.39 | 4.37275E-05 |
| 184 | PA0842 | NA | glycosyl transferase family protein | 5.38 | 8.72814E-08 |
| 185 | PA2343 | *mtlY* | xylulose kinase | 5.38 | 2.79528E-07 |
| 186 | PA0480 | NA | 3-oxoadipate enol-lactonase | 5.38 | 2.79528E-07 |
| 187 | PA2699 | NA | hypothetical protein | 5.30 | 2.121E-08 |
| 188 | PA4649 | NA | hypothetical protein | 5.30 | 4.46792E-07 |
| 189 | PA4892 | *ureF* | urease accessory protein UreF | 5.27 | 2.91953E-06 |
| 190 | PA0525 | NA | denitrification protein NorD | 5.27 | 1.67072E-07 |
| 191 | PA3543 | *algK* | alginate biosynthesis protein AlgK | 5.27 | 6.235E-08 |
| 192 | PA2257 | *pvcD* | paerucumarin biosynthesis protein PvcD | 5.26 | 7.71862E-05 |
| 193 | PA0513 | NA | heme d1 biosynthesis protein NirG | 5.26 | 7.71862E-05 |
| 194 | PA1486 | NA | hypothetical protein | 5.24 | 5.77796E-07 |
| 195 | PA1021 | NA | enoyl-CoA hydratase | 5.24 | 5.93922E-07 |
| **Sr. No.** | **Gene ID** | **Symbol** | **Product name** | **Log FC** | **FDR** |
| 196 | PA2923 | *hisJ* | histidine ABC transporter substrate-binding protein HisJ | 5.18 | 7.9005E-07 |
| 197 | PA2396 | *pvdF* | pyoverdine synthetase F | 5.18 | 2.70275E-07 |
| 198 | PA4884 | NA | hypothetical protein | 5.16 | 8.70272E-07 |
| 199 | PA2515 | *xylL* | 1,6-dihydroxycyclohexa-2,4-diene-1-carboxylate dehydrogenase | 5.16 | 5.39152E-05 |
| 200 | PA2833 | NA | hypothetical protein | 5.16 | 5.08276E-06 |
| 201 | PA1891 | NA | hypothetical protein | 5.12 | 6.80024E-05 |
| 202 | PA3395 | *nosY* | membrane protein NosY | 5.12 | 1.38018E-07 |
| 203 | PA3905 | NA | hypothetical protein | 5.08 | 7.99753E-05 |
| 204 | PA3560 | *fruA* | PTS system fructose-specific transporter subunit IIBC | 5.08 | 4.59066E-07 |
| 205 | PA2133 | NA | hypothetical protein | 5.07 | 7.69971E-06 |
| 206 | PA4814 | *fadH2* | 2,4-dienoyl-CoA reductase | 5.07 | 3.3482E-08 |
| 207 | PA3132 | NA | Hydrolase | 5.04 | 2.10785E-07 |
| 208 | PA4824 | NA | hypothetical protein | 5.03 | 1.69411E-06 |
| 209 | PA2335 | NA | TonB-dependent receptor | 5.00 | 1.10949E-07 |
| 210 | PA1147 | NA | amino acid permease | 4.99 | 1.17992E-07 |
| 211 | PA1694 | *pscQ* | type III secretion system protein | 4.98 | 7.39362E-07 |
| 212 | PA3405 | *hasE* | metalloprotease secretion protein | 4.93 | 9.75718E-07 |
| 213 | PA1237 | NA | multidrug resistance efflux pump | 4.92 | 2.92315E-06 |
| 214 | PA4149 | NA | hypothetical protein | 4.92 | 1.03146E-06 |
| 215 | PA1412 | NA | hypothetical protein | 4.91 | 3.14367E-06 |
| 216 | PA1143 | NA | hypothetical protein | 4.90 | 1.72207E-05 |
| 217 | PA0279 | NA | transcriptional regulator | 4.89 | 1.2229E-06 |
| 218 | PA0184 | NA | ABC transporter ATP-binding protein | 4.88 | 1.2564E-06 |
| 219 | PA2293 | NA | hypothetical protein | 4.87 | 2.02805E-05 |
| 220 | PA3550 | *algF* | alginate o-acetyltransferase AlgF | 4.86 | 5.55856E-07 |
| 221 | PA5431 | NA | transcriptional regulator | 4.84 | 1.89055E-07 |
| 222 | PA1488 | NA | hypothetical protein | 4.84 | 4.72529E-06 |
| 223 | PA4908 | NA | ornithine cyclodeaminase | 4.83 | 6.73583E-07 |
| 224 | PA2933 | NA | major facilitator superfamily transporter | 4.80 | 1.94085E-06 |
| 225 | PA3873 | *narJ* | respiratory nitrate reductase subunit delta | 4.80 | 2.8433E-05 |
| 226 | PA1924 | NA | hypothetical protein | 4.79 | 2.98572E-05 |
| 227 | PA4096 | NA | major facilitator superfamily transporter | 4.79 | 5.19758E-07 |
| 228 | PA2086 | NA | epoxide hydrolase | 4.79 | 2.98572E-05 |
| 229 | PA4864 | *ureD* | urease accessory protein | 4.78 | 1.5195E-06 |
| 230 | PA1019a | NA | Thioesterase | 4.73 | 0.0003 |
| 231 | PA2125 | NA | aldehyde dehydrogenase | 4.72 | 2.92315E-06 |
| 232 | PA1148 | *toxA* | exotoxin A | 4.71 | 2.11109E-07 |
| 233 | PA4121 | NA | hypothetical protein | 4.70 | 1.24257E-06 |
| **Sr. No.** | **Gene ID** | **Symbol** | **Product name** | **Log FC** | **FDR** |
| 234 | PA2984 | NA | hypothetical protein | 4.70 | 1.03398E-07 |
| 235 | PA1695 | *pscP* | translocation protein in type III secretion | 4.69 | 2.38081E-06 |
| 236 | PA2836 | NA | secretion protein | 4.69 | 3.2167E-06 |
| 237 | PA0682 | NA | HxcX atypical pseudopilin | 4.68 | 9.79558E-06 |
| 238 | PA0724 | NA | phage coat protein A | 4.68 | 8.06539E-08 |
| 239 | PA0686 | NA | type II secretion system protein HxcR | 4.67 | 2.65599E-06 |
| 240 | PA2084 | NA | asparagine synthetase | 4.67 | 2.65599E-06 |
| 241 | PA4818 | NA | hypothetical protein | 4.66 | 6.90797E-07 |
| 242 |  | *mdcD* | malonate decarboxylase subunit beta | 4.65 | 1.17225E-05 |
| 243 | PA0241 | NA | major facilitator superfamily transporter | 4.64 | 4.5535E-07 |
| 244 | PA2474 | NA | hypothetical protein | 4.63 | 3.36902E-06 |
| 245 | PA1262 | NA | major facilitator superfamily transporter | 4.56 | 2.60851E-06 |
| 246 | PA5353 | *glcF* | glycolate oxidase iron-sulfur subunit | 4.55 | 3.36536E-06 |
| 247 | PA3911 | NA | hypothetical protein | 4.54 | 1.88632E-05 |
| 248 | PA1356 | NA | hypothetical protein | 4.54 | 7.00242E-06 |
| 249 | PA0219 | NA | aldehyde dehydrogenase | 4.52 | 1.0321E-06 |
| 250 | PA2701 | NA | major facilitator superfamily transporter | 4.49 | 2.43041E-06 |
| 251 | PA0185 | NA | ABC transporter permease | 4.49 | 1.26305E-06 |
| 252 | PA2060 | NA | ABC transporter permease | 4.48 | 9.30546E-06 |
| 253 | PA5385 | *cdhB* | carnitine dehydrogenase | 4.48 | 2.63877E-05 |
| 254 | PA4819 | NA | glycosyl transferase family protein | 4.46 | 1.08893E-05 |
| 255 | PA0110 | NA | hypothetical protein | 4.45 | 1.08893E-05 |
| 256 | PA0222 | NA | hypothetical protein | 4.45 | 2.04282E-06 |
| 257 | PA3448 | NA | ABC transporter permease | 4.44 | 5.68918E-06 |
| 258 | PA2088 | NA | hypothetical protein | 4.44 | 3.18396E-05 |
| 259 | PA1286 | NA | major facilitator superfamily transporter | 4.44 | 2.16211E-06 |
| 260 | PA4599 | *mexC* | resistance-nodulation-cell division (RND) multidrug efflux membrane fusion protein MexC | 4.43 | 2.36121E-06 |
| 261 | PA0273 | NA | major facilitator superfamily transporter | 4.42 | 5.37164E-06 |
| 262 | PA2350 | NA | methionine ABC transporter ATP-binding protein | 4.40 | 7.1203E-06 |
| 263 | PA2325 | NA | hypothetical protein | 4.39 | 1.4655E-05 |
| 264 | PA0699 | NA | PpiC-type peptidyl-prolyl cis-trans isomerase | 4.39 | 4.07009E-05 |
| 265 | PA1216 | NA | hypothetical protein | 4.39 | 6.4266E-06 |
| 266 | PA0056 | NA | transcriptional regulator | 4.38 | 2.91604E-06 |
| 267 | PA0238 | NA | hypothetical protein | 4.37 | 8.05538E-06 |
| 268 | PA3394 | *nosF* | copper ABC transporter ATP-binding | 4.37 | 1.65505E-05 |
| **Sr. No.** | **Gene ID** | **Symbol** | **Product name** | **Log FC** | **FDR** |
| 269 | PA3442 | NA | aliphatic sulfonates ABC transporter ATP-binding subunit | 4.36 | 4.52001E-05 |
| 270 | PA1214 | NA | hypothetical protein | 4.36 | 1.71322E-05 |
| 271 | PA2214 | NA | major facilitator superfamily transporter | 4.35 | 1.07361E-06 |
| 272 | PA3780 | NA | hypothetical protein | 4.35 | 0.0002 |
| 273 | PA2120 | NA | hypothetical protein | 4.34 | 5.00763E-05 |
| 274 | PA1231 | NA | hypothetical protein | 4.34 | 0.0002 |
| 275 | PA0118 | NA | hypothetical protein | 4.32 | 1.56862E-06 |
| 276 | PA2729 | NA | hypothetical protein | 4.32 | 7.58704E-07 |
| 277 | PA3589 | NA | acetyl-CoA acetyltransferase | 4.31 | 2.1781E-05 |
| 278 | PA0882 | NA | hypothetical protein | 4.28 | 1.28758E-05 |
| 279 | PA3542 | *alg44* | alginate biosynthesis protein Alg44 | 4.27 | 2.15204E-06 |
| 280 | PA2314 | NA | major facilitator superfamily transporter | 4.26 | 5.37164E-06 |
| 281 | PA1855 | NA | hypothetical protein | 4.26 | 0.0003 |
| 282 | PA2213 | NA | Porin | 4.26 | 1.20688E-05 |
| 283 | PA2371 | NA | ClpA/B-type protease | 4.26 | 1.74233E-06 |
| 284 | PA2715 | NA | Ferredoxin | 4.26 | 0.0003 |
| 285 | PA0205 | NA | ABC transporter permease | 4.22 | 3.22807E-06 |
| 286 | PA4099 | NA | hypothetical protein | 4.22 | 2.79473E-06 |
| 287 | PA2473 | NA | glutathione S-transferase | 4.21 | 8.7287E-05 |
| 288 | PA3444 | NA | alkanesulfonate monooxygenase | 4.21 | 7.03044E-06 |
| 289 | PA2863 | *lipH* | lipase chaperone | 4.20 | 1.08071E-05 |
| 290 | PA0136 | NA | ABC transporter ATP-binding protein | 4.17 | 4.25493E-06 |
| 291 | PA1908 | NA | major facilitator superfamily transporter | 4.16 | 9.29948E-06 |
| 292 | PA2462 | NA | hypothetical protein | 4.14 | 7.71393E-07 |
| 293 | PA0194 | NA | hypothetical protein | 4.13 | 7.71769E-06 |
| 294 | PA4859 | NA | ABC transporter permease | 4.13 | 5.20607E-06 |
| 295 | PA3773 | NA | hypothetical protein | 4.13 | 1.51405E-05 |
| 296 | PA3592 | NA | hypothetical protein | 4.12 | 1.10306E-05 |
| 297 | PA2689 | NA | hypothetical protein | 4.11 | 5.62043E-05 |
| 298 | PA1827 | NA | short-chain dehydrogenase | 4.11 | 2.47374E-05 |
| 299 | PA1020 | NA | acyl-CoA dehydrogenase | 4.10 | 1.2209E-05 |
| 300 | PA2922 | NA | Hydrolase | 4.09 | 1.77328E-05 |
| 301 | PA3904 | NA | hypothetical protein | 4.09 | 2.82303E-05 |
| 302 | PA4092 | *hpaC* | 4-hydroxyphenylacetate 3-monooxygenase small subunit | 4.08 | 6.35353E-05 |
| 303 | PA5391 | NA | hypothetical protein | 4.07 | 6.67419E-05 |
| 304 | PA0987 | NA | hypothetical protein | 4.07 | 3.08219E-05 |
| 305 | PA1980 | *eraR* | response regulator EraR | 4.07 | 0.0001 |
| 306 | PA3885 | *tpbA* | protein tyrosine phosphatase TpbA | 4.06 | 3.5273E-05 |
| 307 | PA1497 | NA | Transporter | 4.06 | 2.04756E-05 |
| **Sr. No.** | **Gene ID** | **Symbol** | **Product name** | **Log FC** | **FDR** |
| 308 | PA2141 | NA | hypothetical protein | 4.06 | 0.0007 |
| 309 | PA4103 | NA | hypothetical protein | 4.06 | 5.68918E-06 |
| 310 | PA3591 | NA | enoyl-CoA hydratase | 4.06 | 7.32239E-05 |
| 311 | PA4188 | NA | hypothetical protein | 4.05 | 3.28091E-05 |
| 312 | PA1780 | *nirD* | assimilatory nitrite reductase small subunit | 4.05 | 0.0001 |
| 313 | PA2373 | NA | hypothetical protein | 4.04 | 3.71697E-06 |
| 314 | PA0802 | NA | hypothetical protein | 4.03 | 0.0001 |
| 315 | PA1725 | *pscL* | type III secretion system protein | 4.03 | 7.99753E-05 |
| 316 | PA4083 | *cupB4* | chaperone CupB4 | 4.03 | 2.44802E-05 |
| 317 | PA4038 | NA | hypothetical protein | 4.02 | 1.33829E-05 |
| 318 | PA5419 | *soxG* | sarcosine oxidase subunit gamma | 4.02 | 1.33829E-05 |
| 319 | PA1186 | NA | hypothetical protein | 4.00 | 4.14744E-05 |
| 320 | PA2458 | NA | hypothetical protein | 4.00 | 4.28134E-06 |
| 321 | PA1489 | NA | hypothetical protein | 3.99 | 2.98572E-05 |
| 322 | PA0726 | NA | hypothetical protein | 3.99 | 3.14367E-06 |
| 323 | PA2124 | NA | Dehydrogenase | 3.97 | 1.77152E-05 |
| 324 | PA2036 | NA | hypothetical protein | 3.97 | 1.40401E-05 |
| 325 | PA3436 | NA | hypothetical protein | 3.94 | 2.61211E-05 |
| 326 | PA0752 | NA | hypothetical protein | 3.94 | 1.10118E-05 |
| 327 | PA0521 | NA | cytochrome C oxidase subunit | 3.93 | 1.26229E-05 |
| 328 | PA3535 | NA | serine protease | 3.93 | 5.19922E-06 |
| 329 | PA3871 | NA | PpiC-type peptidyl-prolyl cis-trans isomerase | 3.92 | 4.01341E-05 |
| 330 | PA0725 | NA | hypothetical protein | 3.92 | 0.0001 |
| 331 | PA2924 | *hisQ* | histidine ABC transporter permease HisQ | 3.91 | 6.17218E-05 |
| 332 | PA3037 | NA | hypothetical protein | 3.90 | 7.46055E-05 |
| 333 | PA4107 | NA | hypothetical protein | 3.89 | 4.71561E-05 |
| 334 | PA0221 | NA | Aminotransferase | 3.89 | 2.08372E-05 |
| 335 | PA3396 | *nosL* | acessory protein NosL | 3.88 | 0.0003 |
| 336 | PA0798 | *pmtA* | phospholipid methyltransferase | 3.87 | 7.61222E-05 |
| 337 | PA1917 | NA | hypothetical protein | 3.86 | 0.0001 |
| 338 | PA5144 | NA | hypothetical protein | 3.86 | 0.0001 |
| 339 | PA0144 | NA | hypothetical protein | 3.86 | 5.48473E-05 |
| 340 | PA1274 | NA | 5,6-dimethylbenzimidazole synthase | 3.86 | 9.16933E-05 |
| 341 | PA1298 | NA | hypothetical protein | 3.86 | 9.16933E-05 |
| 342 | PA4105 | NA | hypothetical protein | 3.84 | 9.92969E-05 |
| 343 | PA2324 | NA | hypothetical protein | 3.83 | 4.52001E-05 |
| 344 | PA0684 | NA | type II secretion system protein | 3.83 | 0.0004 |
| 345 | PA3376 | NA | phosphonate C-P lyase system protein PhnK | 3.83 | 0.0004 |
| 346 | PA4820 | NA | hypothetical protein | 3.82 | 4.66203E-05 |
| 347 | PA4918 | NA | hypothetical protein | 3.82 | 4.88946E-06 |
| **Sr. No.** | **Gene ID** | **Symbol** | **Product name** | **Log FC** | **FDR** |
| 348 | PA5418 | *soxA* | sarcosine oxidase subunit alpha | 3.82 | 7.61629E-06 |
| 349 | PA4985 | NA | hypothetical protein | 3.81 | 3.71249E-05 |
| 350 | PA2892 | *atuG* | short-chain dehydrogenase | 3.80 | 5.03768E-05 |
| 351 | PA4593 | NA | ABC transporter permease | 3.80 | 3.91345E-05 |
| 352 | PA4978 | NA | hypothetical protein | 3.80 | 1.67058E-05 |
| 353 | PA2471 | NA | hypothetical protein | 3.79 | 7.21024E-05 |
| 354 | PA3383 | NA | phosphonate ABC transporter substrate-binding protein | 3.79 | 1.68888E-05 |
| 355 | PA1022 | NA | acyl-CoA dehydrogenase | 3.79 | 1.56427E-05 |
| 356 | PA4179 | NA | Porin | 3.79 | 8.9392E-06 |
| 357 | PA3447 | NA | ABC transporter ATP-binding protein | 3.79 | 0.0002 |
| 358 | PA0237 | NA | Oxidoreductase | 3.78 | 5.64353E-05 |
| 359 | PA4392 | NA | hypothetical protein | 3.78 | 5.64353E-05 |
| 360 | PA5400 | NA | electron transfer flavoprotein subunit alpha | 3.77 | 5.97376E-05 |
| 361 | PA2342 | *mtlD* | mannitol dehydrogenase | 3.77 | 4.7284E-05 |
| 362 | PA0212 | *mdcE* | malonate decarboxylase subunit gamma | 3.76 | 0.0002 |
| 363 | PA2255 | *pvcB* | paerucumarin biosynthesis protein PvcB | 3.73 | 0.0001 |
| 364 | PA1876 | NA | ABC transporter ATP-binding protein/permease | 3.73 | 3.18396E-05 |
| 365 | PA1696 | *pscO* | translocation protein in type III secretion | 3.71 | 0.0007 |
| 366 | PA4622 | NA | major facilitator superfamily transporter | 3.71 | 1.32569E-05 |
| 367 | PA1219 | NA | hypothetical protein | 3.71 | 0.0007 |
| 368 | PA4795 | NA | hypothetical protein | 3.71 | 0.0007 |
| 369 | PA1490 | NA | transcriptional regulator | 3.71 | 7.95014E-05 |
| 370 | PA0466 | NA | hypothetical protein | 3.70 | 0.0007 |
| 371 | PA2131 | *cupA4* | fimbrial subunit CupA4 | 3.69 | 8.42475E-05 |
| 372 | PA0526 | NA | hypothetical protein | 3.69 | 0.0001 |
| 373 | PA2295 | NA | ABC transporter permease | 3.68 | 8.9497E-05 |
| 374 | PA2217 | NA | aldehyde dehydrogenase | 3.68 | 5.80724E-05 |
| 375 | PA5341 | NA | hypothetical protein | 3.67 | 9.21863E-05 |
| 376 | PA2078 | NA | hypothetical protein | 3.66 | 3.27673E-05 |
| 377 | PA1023 | NA | short-chain dehydrogenase | 3.66 | 7.75714E-05 |
| 378 | PA2349 | NA | hypothetical protein | 3.65 | 0.0001 |
| 379 | PA5354 | NA | glycolate oxidase FAD binding subunit | 3.65 | 0.0001 |
| 380 | PA2216 | NA | hypothetical protein | 3.64 | 8.18882E-05 |
| 381 | PA5266 | NA | hypothetical protein | 3.64 | 1.67058E-05 |
| 382 | PA4148 | NA | short-chain dehydrogenase | 3.64 | 0.0001 |
| 383 | PA1711 | NA | hypothetical protein | 3.63 | 0.0004 |
| **Sr. No.** | **Gene ID** | **Symbol** | **Product name** | **Log FC** | **FDR** |
| 384 | PA3433 | NA | transcriptional regulator | 3.63 | 3.22656E-05 |
| 385 | PA4802 | NA | hypothetical protein | 3.62 | 2.08372E-05 |
| 386 | PA3358 | NA | hypothetical protein | 3.62 | 7.4063E-05 |
| 387 | PA2163 | NA | 4-alpha-glucanotransferase | 3.61 | 9.37843E-05 |
| 388 | PA0137 | NA | ABC transporter permease | 3.61 | 2.98572E-05 |
| 389 | PA3036 | NA | hypothetical protein | 3.61 | 5.55792E-05 |
| 390 | PA1952 | NA | hypothetical protein | 3.61 | 0.0004 |
| 391 | PA2439 | NA | hypothetical protein | 3.61 | 5.32018E-05 |
| 392 | PA0193 | NA | hypothetical protein | 3.60 | 9.92969E-05 |
| 393 | PA0192 | NA | TonB-dependent receptor | 3.60 | 2.0619E-05 |
| 394 | PA3544 | *algE* | alginate production protein AlgE | 3.60 | 2.96629E-05 |
| 395 | PA3320 | NA | hypothetical protein | 3.60 | 5.41779E-05 |
| 396 | PA2925 | *hisM* | histidine ABC transporter permease HisM | 3.60 | 9.92969E-05 |
| 397 | PA2650 | NA | hypothetical protein | 3.60 | 5.52034E-05 |
| 398 | PA0111 | NA | hypothetical protein | 3.60 | 0.0005 |
| 399 | PA2179 | NA | hypothetical protein | 3.60 | 0.0001 |
| 400 | PA5328 | NA | mono-heme cytochrome C | 3.58 | 0.0005 |
| 401 | PA1743 | NA | hypothetical protein | 3.58 | 0.0005 |
| 402 | PA1893 | NA | hypothetical protein | 3.57 | 3.30626E-05 |
| 403 | PA2589 | NA | hypothetical protein | 3.56 | 4.7776E-05 |
| 404 | PA0523 | *norC* | nitric oxide reductase subunit C | 3.56 | 6.00389E-05 |
| 405 | PA2369 | NA | hypothetical protein | 3.55 | 7.32239E-05 |
| 406 | PA3609 | *potC* | polyamine ABC transporter permease PotC | 3.55 | 7.32239E-05 |
| 407 | PA1346 | NA | hypothetical protein | 3.55 | 8.55758E-05 |
| 408 | PA2370 | NA | hypothetical protein | 3.54 | 0.0006 |
| 409 | PA1983 | *exaB* | cytochrome C550 | 3.54 | 0.0002 |
| 410 | PA4181 | NA | hypothetical protein | 3.53 | 5.62043E-05 |
| 411 | PA3908 | NA | hypothetical protein | 3.53 | 8.18882E-05 |
| 412 | PA4650 | NA | hypothetical protein | 3.52 | 0.0001 |
| 413 | PA4862 | NA | ABC transporter ATP-binding protein | 3.52 | 0.0002 |
| 414 | PA3373 | NA | hypothetical protein | 3.52 | 0.0007 |
| 415 | PA2463 | NA | hypothetical protein | 3.51 | 4.83297E-05 |
| 416 | PA3875 | *narG* | respiratory nitrate reductase subunit alpha | 3.51 | 2.0619E-05 |
| 417 | PA3519 | NA | hypothetical protein | 3.51 | 3.30626E-05 |
| 418 | PA1213 | NA | hypothetical protein | 3.51 | 0.0002 |
| 419 | PA3561 | *fruK* | 1-phosphofructokinase | 3.51 | 0.0002 |
| 420 | PA1954 | NA | hypothetical protein | 3.51 | 3.3813E-05 |
| 421 | PA5282 | NA | major facilitator superfamily transporter | 3.51 | 5.02286E-05 |
| 422 | PA0718 | NA | hypothetical protein | 3.51 | 6.14091E-05 |
| **Sr. No.** | **Gene ID** | **Symbol** | **Product name** | **Log FC** | **FDR** |
| 423 | PA0324 | NA | ABC transporter permease | 3.50 | 4.73741E-05 |
| 424 | PA1212 | NA | major facilitator superfamily transporter | 3.50 | 6.84991E-05 |
| 425 | PA1786 | NA | hypothetical protein | 3.50 | 6.24084E-05 |
| 426 | PA0474 | NA | Esterase | 3.50 | 0.0004 |
| 427 | PA4798 | NA | hypothetical protein | 3.48 | 3.37782E-05 |
| 428 | PA4982 | NA | two-component sensor | 3.48 | 5.79529E-05 |
| 429 | PA1279 | *cobU* | nicotinate-nucleotide--dimethylbenzimidazole phosphoribosyltransferase | 3.47 | 0.0003 |
| 430 | PA2835 | NA | major facilitator superfamily transporter | 3.46 | 6.14091E-05 |
| 431 | PA0242 | NA | hypothetical protein | 3.46 | 5.93033E-05 |
| 432 | PA5386 | *cdhA* | 3-hydroxybutyryl-CoA dehydrogenase | 3.45 | 0.0001 |
| 433 | PA5326 | NA | hypothetical protein | 3.45 | 9.92969E-05 |
| 434 | PA4592 | NA | hypothetical protein | 3.44 | 7.21024E-05 |
| 435 | PA2085 | NA | ring-hydroxylating dioxygenase small subunit | 3.44 | 0.0009 |
| 436 | PA5399 | *dgcB* | dimethylglycine catabolism protein DgcB | 3.43 | 6.46809E-05 |
| 437 | PA5159 | NA | multidrug resistance protein | 3.43 | 9.37843E-05 |
| 438 | PA5470 | NA | peptide chain release factor-like protein | 3.43 | 9.55376E-05 |
| 439 | PA0173 | NA | chemotaxis response regulator protein-glutamate methylesterase | 3.42 | 0.0002 |
| 440 | PA2362 | NA | hypothetical protein | 3.41 | 0.0006 |
| 441 | PA1848 | NA | major facilitator superfamily transporter | 3.41 | 0.0001 |
| 442 | PA0117 | NA | short-chain dehydrogenase | 3.41 | 0.0001 |
| 443 | PA2890 | *atuE* | isohexenylglutaconyl-CoA hydratase | 3.41 | 0.0004 |
| 444 | PA2837 | NA | hypothetical protein | 3.39 | 0.0001 |
| 445 | PA3772 | NA | hypothetical protein | 3.39 | 7.77875E-05 |
| 446 | PA3547 | *algL* | alginate lyase | 3.39 | 8.47958E-05 |
| 447 | PA0883 | NA | acyl-CoA lyase subunit beta | 3.38 | 0.0003 |
| 448 | PA4920 | NA*dE* | NAD synthetase | 3.37 | 5.13913E-05 |
| 449 | PA1236 | NA | major facilitator superfamily transporter | 3.35 | 0.0001 |
| 450 | PA0244 | NA | shikimate 5-dehydrogenase | 3.34 | 0.0003 |
| 451 | PA1634 | *kdpB* | potassium-transporting ATPase subunit B | 3.34 | 0.0001 |
| 452 | PA5115 | NA | hypothetical protein | 3.33 | 0.0003 |
| 453 | PA2348 | NA | hypothetical protein | 3.33 | 0.0004 |
| 454 | PA0786 | NA | Transporter | 3.33 | 0.0008 |
| 455 | PA2347 | NA | hypothetical protein | 3.33 | 0.0008 |
| 456 | PA3546 | *algX* | alginate biosynthesis protein AlgX | 3.33 | 0.0001 |
| **Sr. No.** | **Gene ID** | **Symbol** | **Product name** | **Log FC** | **FDR** |
| 457 | PA3518 | NA | hypothetical protein | 3.33 | 0.0001 |
| 458 | PA2356 | *msuD* | methanesulfonate monooxygenase | 3.32 | 0.0004 |
| 459 | PA4861 | NA | ABC transporter ATP-binding protein | 3.32 | 0.0001 |
| 460 | PA2155 | NA | cardiolipin synthase 2 | 3.32 | 0.0002 |
| 461 | PA2061 | NA | ABC transporter ATP-binding protein | 3.32 | 0.0002 |
| 462 | PA1218 | NA | hypothetical protein | 3.32 | 0.0002 |
| 463 | PA4299 | *tadD* | type II secretion system protein TadD | 3.32 | 0.0009 |
| 464 | PA3870 | *moaA1* | molybdenum cofactor biosynthesis protein A | 3.32 | 0.0002 |
| 465 | PA5395 | NA | hypothetical protein | 3.31 | 0.0003 |
| 466 | PA1929 | NA | hypothetical protein | 3.31 | 0.0009 |
| 467 | PA2284 | NA | hypothetical protein | 3.30 | 0.0002 |
| 468 | PA2346 | NA | hypothetical protein | 3.30 | 0.0002 |
| 469 | PA0058 | NA | hypothetical protein | 3.29 | 0.0004 |
| 470 | PA2039 | NA | hypothetical protein | 3.28 | 9.41752E-05 |
| 471 | PA0175 | NA | chemotaxis protein methyltransferase | 3.28 | 0.0001 |
| 472 | PA2803 | NA | hypothetical protein | 3.28 | 0.0003 |
| 473 | PA1435 | NA | resistance-nodulation-cell division (RND) efflux membrane fusion protein | 3.28 | 0.0002 |
| 474 | PA0029 | NA | sulfate transporter | 3.27 | 0.0001 |
| 475 | PA0150 | NA | transmembrane sensor | 3.27 | 0.0001 |
| 476 | PA2596 | NA | hypothetical protein | 3.27 | 0.0002 |
| 477 | PA3360 | NA | secretion protein | 3.27 | 0.0003 |
| 478 | PA4098 | NA | short-chain dehydrogenase | 3.27 | 0.0003 |
| 479 | PA3291 | NA | hypothetical protein | 3.26 | 0.0003 |
| 480 | PA2676 | NA | type II secretion system protein | 3.26 | 0.0001 |
| 481 | PA4586 | NA | hypothetical protein | 3.25 | 0.0005 |
| 482 | PA3416 | NA | pyruvate dehydrogenase E1 component subunit beta | 3.25 | 0.0003 |
| 483 | PA3608 | *potB* | polyamine ABC transporter permease PotB | 3.25 | 0.0002 |
| 484 | PA4189 | NA | aldehyde dehydrogenase | 3.25 | 0.0002 |
| 485 | PA3907 | NA | hypothetical protein | 3.25 | 0.0008 |
| 486 | PA0103 | NA | sulfate transporter | 3.24 | 0.0001 |
| 487 | PA3133 | NA | transcriptional regulator | 3.24 | 0.0002 |
| 488 | PA1270 | NA | hypothetical protein | 3.24 | 0.0001 |
| 489 | PA3912 | NA | hypothetical protein | 3.24 | 0.0006 |
| 490 | PA2421 | NA | hypothetical protein | 3.23 | 0.0001 |
| 491 | PA4883 | NA | hypothetical protein | 3.22 | 0.0006 |
| 492 | PA0511 | *nirJ* | heme d1 biosynthesis protein NirJ | 3.22 | 0.0002 |
| 493 | PA4137 | NA | Porin | 3.21 | 0.0001 |
| 494 | PA0236 | NA | transcriptional regulator | 3.21 | 0.0002 |
| 495 | PA1251 | NA | chemotaxis transducer | 3.21 | 0.0001 |
| **Sr. No.** | **Gene ID** | **Symbol** | **Product name** | **Log FC** | **FDR** |
| 496 | PA4830 | NA | hypothetical protein | 3.21 | 0.0003 |
| 497 | PA2132 | *cupA5* | chaperone CupA5 | 3.20 | 0.0009 |
| 498 | PA2289 | NA | hypothetical protein | 3.20 | 0.0001 |
| 499 | PA0214 | NA | acyl transferase | 3.19 | 0.0006 |
| 500 | PA1313 | NA | major facilitator superfamily transporter | 3.18 | 0.0002 |
| 501 | PA1782 | NA | serine/threonine-protein kinase | 3.18 | 0.0002 |
| 502 | PA2181 | NA | glutamate--cysteine ligase | 3.18 | 0.0003 |
| 503 | PA2229 | NA | hypothetical protein | 3.18 | 0.0003 |
| 504 | PA5420 | *purU2* | formyltetrahydrofolate deformylase | 3.17 | 0.0002 |
| 505 | PA0197 | *tonB2* | transporter TonB | 3.17 | 0.0008 |
| 506 | PA0702 | NA | hypothetical protein | 3.17 | 0.0005 |
| 507 | PA0166 | NA | Transporter | 3.17 | 0.0002 |
| 508 | PA4095 | NA | hypothetical protein | 3.17 | 0.0006 |
| 509 | PA4167 | NA | 2,5-diketo-D-gluconate reductase B | 3.16 | 0.0001 |
| 510 | PA3119 | NA | hypothetical protein | 3.16 | 0.0003 |
| 511 | PA0052 | NA | hypothetical protein | 3.15 | 0.0004 |
| 512 | PA3607 | *potA* | polyamine transporter ATP-binding protein PotA | 3.15 | 0.0002 |
| 513 | PA0252 | NA | hypothetical protein | 3.15 | 0.0003 |
| 514 | PA2056 | NA | transcriptional regulator | 3.15 | 0.0007 |
| 515 | PA4087 | NA | hypothetical protein | 3.15 | 0.0003 |
| 516 | PA4652 | NA | hypothetical protein | 3.14 | 0.0001 |
| 517 | PA2057 | NA | hypothetical protein | 3.14 | 0.0001 |
| 518 | PA5352 | NA | hypothetical protein | 3.13 | 0.0007 |
| 519 | PA2243 | *pslM* | FAD-binding dehydrogenase | 3.13 | 0.0003 |
| 520 | PA2431 | NA | hypothetical protein | 3.13 | 0.0002 |
| 521 | PA3443 | NA | ABC transporter permease | 3.13 | 0.0005 |
| 522 | PA0326 | NA | ABC transporter ATP-binding protein | 3.12 | 0.0002 |
| 523 | PA3415 | NA | branched-chain alpha-keto acid dehydrogenase subunit E2 | 3.12 | 0.0002 |
| 524 | PA4027a | NA | hypothetical protein | 3.12 | 0.0002 |
| 525 | PA0875 | NA | hypothetical protein | 3.11 | 0.0001 |
| 526 | PA4343 | NA | major facilitator superfamily transporter | 3.11 | 0.0002 |
| 527 | PA1215 | NA | hypothetical protein | 3.10 | 0.0004 |
| 528 | PA3884 | NA | hypothetical protein | 3.10 | 0.0007 |
| 529 | PA2783 | NA | hypothetical protein | 3.09 | 0.0002 |
| 530 | PA4120 | NA | transcriptional regulator | 3.08 | 0.0005 |
| 531 | PA3384 | *phnC* | phosphonate ABC transporter ATP-binding protein | 3.08 | 0.0005 |
| 532 | PA0693 | *exbB2* | transporter ExbB | 3.08 | 0.0004 |
| 533 | PA2076 | NA | transcriptional regulator | 3.07 | 0.0009 |
| 534 | PA3323 | NA | hypothetical protein | 3.06 | 0.0005 |
| **Sr. No.** | **Gene ID** | **Symbol** | **Product name** | **Log FC** | **FDR** |
| 535 | PA3936 | NA | taurine ABC transporter permease | 3.06 | 0.0007 |
| 536 | PA1092 | *fliC* | B-type flagellin | 3.06 | 0.0001 |
| 537 | PA0477 | NA | transcriptional regulator | 3.05 | 0.0002 |
| 538 | PA4191 | NA | iron/ascorbate oxidoreductase | 3.04 | 0.0005 |
| 539 | PA1975 | NA | hypothetical protein | 3.04 | 0.0006 |
| 540 | PA5132 | NA | hypothetical protein | 3.03 | 0.0003 |
| 541 | PA3521 | NA | hypothetical protein | 3.02 | 0.0004 |
| 542 | PA1633 | *kdpA* | potassium-transporting ATPase subunit A | 3.02 | 0.0004 |
| 543 | PA0311 | NA | hypothetical protein | 3.02 | 0.0005 |
| 544 | PA5539 | NA | GTP cyclohydrolase | 3.02 | 0.0005 |
| 545 | PA4903 | NA | major facilitator superfamily transporter | 3.02 | 0.0006 |
| 546 | PA4073 | NA | aldehyde dehydrogenase | 3.02 | 0.0004 |
| 547 | PA2670 | NA | hypothetical protein | 3.02 | 0.0003 |
| 548 | PA5387 | *cdhC* | carnitine dehydrogenase | 3.01 | 0.0006 |
| 549 | PA3749 | NA | major facilitator superfamily transporter | 3.01 | 0.0002 |
| 550 | PA2520 | *czcA* | resistance-nodulation-cell division (RND) divalent metal cation efflux transporter CzcA | 3.01 | 0.0003 |
| 551 | PA3750 | NA | hypothetical protein | 3.01 | 0.0009 |
| 552 | PA1281 | *cobV* | adenosylcobinamide-GDP ribazoletransferase | 3.00 | 0.0004 |
| 553 | PA2097 | NA | flavin-binding monooxygenase | 3.00 | 0.0006 |
| 554 | PA2598 | NA | hypothetical protein | 3.00 | 0.0006 |
| 555 | PA3937 | NA | taurine ABC transporter ATP-binding protein | 3.00 | 0.0009 |
| 556 | PA0189 | NA | Porin | 2.99 | 0.0005 |
| 557 | PA1017 | *pauA* | pimeloyl-CoA synthetase | 2.99 | 0.0003 |
| 558 | PA1172 | NA*pC* | cytochrome C protein NapC | 2.98 | 0.0007 |
| 559 | PA0685 | NA | type II secretion system protein | 2.97 | 0.0003 |
| 560 | PA3393 | *nosD* | copper-binding periplasmic protein | 2.95 | 0.0006 |
| 561 | PA1232 | NA | hypothetical protein | 2.95 | 0.0004 |
| 562 | PA3296 | *phoA* | alkaline phosphatase | 2.95 | 0.0003 |
| 563 | PA1417 | NA | hypothetical protein | 2.95 | 0.0006 |
| 564 | PA0057 | NA | hypothetical protein | 2.95 | 0.0006 |
| 565 | PA0153 | *pcaH* | protocatechuate 3,4-dioxygenase subunit beta | 2.94 | 0.0005 |
| 566 | PA1783 | NA*sA* | nitrate transporter | 2.94 | 0.0007 |
| 567 | PA1781 | *nirB* | assimilatory nitrite reductase large subunit | 2.94 | 0.0004 |
| 568 | PA4911 | NA | branched-chain amino acid ABC transporter permease | 2.93 | 0.0005 |
| 569 | PA3391 | *nosR* | regulatory protein NosR | 2.93 | 0.0003 |
| **Sr. No.** | **Gene ID** | **Symbol** | **Product name** | **Log FC** | **FDR** |
| 570 | PA0433 | NA | hypothetical protein | 2.93 | 0.0009 |
| 571 | PA5158 | NA | hypothetical protein | 2.93 | 0.0005 |
| 572 | PA1138 | NA | transcriptional regulator | 2.92 | 0.0007 |
| 573 | PA3409 | NA | transmembrane sensor | 2.91 | 0.0006 |
| 574 | PA4037 | NA | ABC transporter ATP-binding protein | 2.91 | 0.0006 |
| 575 | PA0051 | *phzH* | phenazine-modifying protein | 2.90 | 0.0005 |
| 576 | PA2283 | NA | hypothetical protein | 2.90 | 0.0005 |
| 577 | PA3954 | NA | hypothetical protein | 2.90 | 0.0005 |
| 578 | PA4166 | NA | Acetyltransferase | 2.90 | 0.0009 |
| 579 | PA2691 | NA | hypothetical protein | 2.89 | 0.0007 |
| 580 | PA4898 | *opdK* | vanillate porin OpdK | 2.89 | 0.0007 |
| 581 | PA4190 | *pqsL* | Monooxygenase | 2.89 | 0.0008 |
| 582 | PA5544 | NA | hypothetical protein | 2.88 | 0.0004 |
| 583 | PA1923 | NA | cobaltochelatase subunit CobN | 2.88 | 0.0006 |
| 584 | PA1922 | NA | TonB-dependent receptor | 2.88 | 0.0006 |
| 585 | PA0528 | NA | transcriptional regulator | 2.87 | 0.0004 |
| 586 | PA1027 | NA | aldehyde dehydrogenase | 2.87 | 0.0006 |
| 587 | PA3176 | *gltS* | glutamate/sodium ion symporter GltS | 2.87 | 0.0007 |
| 588 | PA4342 | NA | Amidase | 2.87 | 0.0007 |
| 589 | PA1360 | NA | hypothetical protein | 2.86 | 0.0008 |
| 590 | PA0516 | *nirF* | heme d1 biosynthesis protein NirF | 2.86 | 0.0007 |
| 591 | PA4136 | NA | major facilitator superfamily transporter | 2.86 | 0.0005 |
| 592 | PA1373 | *fabF2* | 3-oxoacyl-ACP synthase | 2.86 | 0.0006 |
| 593 | PA4100 | NA | Dehydrogenase | 2.86 | 0.0008 |
| 594 | PA5294 | NA | multidrug efflux protein NorA | 2.85 | 0.0007 |
| 595 | PA1400 | NA | pyruvate carboxylase | 2.85 | 0.0005 |
| 596 | PA3392 | *nosZ* | nitrous-oxide reductase | 2.85 | 0.0005 |
| 597 | PA3619 | NA | hypothetical protein | 2.84 | 0.0005 |
| 598 | PA0435 | NA | hypothetical protein | 2.84 | 0.0008 |
| 599 | PA3532 | NA | hypothetical protein | 2.82 | 0.0005 |
| 600 |  | *pfeA* | ferric enterobactin receptor | 2.82 | 0.0005 |
| 601 | PA0143 | *nuh* | nonspecific ribonucleoside hydrolase | 2.82 | 0.0007 |
| 602 | PA1019 | *mucK* | cis,cis-muconate transporter MucK | 2.81 | 0.0008 |
| 603 | PA0781 | NA | hypothetical protein | 2.81 | 0.0006 |
| 604 | PA3638 | NA | tRNA (Ile)-lysidine synthase | 2.80 | 0.0006 |
| 605 | PA4540 | NA | hypothetical protein | 2.80 | 0.0007 |
| 606 | PA4779 | NA | hypothetical protein | 2.80 | 0.0009 |
| 607 | PA3562 | *fruI* | PTS system fructose-specific transporter subunit FruI | 2.79 | 0.0007 |
| 608 | PA1450 | NA | hypothetical protein | 2.79 | 0.0007 |
| 609 | PA4648 | NA | hypothetical protein | 2.79 | 0.0008 |
| 610 | PA5393 | NA | hypothetical protein | 2.77 | 0.0009 |
| **Sr. No.** | **Gene ID** | **Symbol** | **Product name** | **Log FC** | **FDR** |
| 611 | PA5160 | NA | drug efflux transporter | 2.74 | 0.0007 |
| 612 | PA3760 | NA | N-acetyl-D-glucosamine phosphotransferase system transporter | 2.73 | 0.0008 |
| 613 | PA0183 | *atsA* | Arylsulfatase | 2.73 | 0.0008 |
| 614 | PA0524 | *norB* | nitric oxide reductase subunit B | 2.73 | 0.0009 |
| 615 | PA5430 | NA | hypothetical protein | 2.71 | 0.0009 |
| 616 | PA5471 | NA | hypothetical protein | 2.68 | 0.0009 |

Genes are arranged in decreasing order of Fold Change; Databases consulted for gene functions were: NCBI gene database (https://www.ncbi.nlm.nih.gov/nuccore/NC_002516); KEGG (Kyoto Encyclopedia of Genes and Genomes: https://www.genome.jp/kegg/); Uniprot (<https://www.uniprot.org/>). NA: Not Applicable; FDR: False Discovery Rate

**Table S7. Node degree score of the top up-regulated genes**

| **No.** | **Gene ID/Symbol** | **Identifier** | **Node degree** |
| --- | --- | --- | --- |
| 1 | *wbpI* | 208964.PA3148 | 27 |
| 2 | PA0192 | 208964.PA0192 | 26 |
| 3 | PA2730 | 208964.PA2730 | 26 |
| 4 | PA3501 | 208964.PA3501 | 25 |
| 5 | *wbpJ* | 208964.PA3147 | 25 |
| 6 | PA2090 | 208964.PA2090 | 24 |
| 7 | *wbpB* | 208964.PA3158 | 24 |
| 8 | *wbpD* | 208964.PA3156 | 24 |
| 9 | *wbpG* | 208964.PA3150 | 24 |
| 10 | *wbpH* | 208964.PA3149 | 24 |
| 11 | *wbpK* | 208964.PA3146 | 24 |
| 12 | PA1786 | 208964.PA1786 | 23 |
| 13 | PA3498 | 208964.PA3498 | 23 |
| 14 | *hisF2* | 208964.PA3151 | 23 |
| 15 | *hisH2* | 208964.PA3152 | 23 |
| 16 | *norB* | 208964.PA0524 | 23 |
| 17 | *nosZ* | 208964.PA3392 | 23 |
| 18 | *wbpE* | 208964.PA3155 | 23 |
| 19 | PA0184 | 208964.PA0184 | 22 |
| 20 | PA0185 | 208964.PA0185 | 22 |
| 21 | PA0521 | 208964.PA0521 | 22 |
| 22 | PA1372 | 208964.PA1372 | 22 |
| 23 | PA3513 | 208964.PA3513 | 22 |
| 24 | PA3514 | 208964.PA3514 | 22 |
| 25 | *nirF* | 208964.PA0516 | 22 |
| 26 | *nosD* | 208964.PA3393 | 22 |
| 27 | *nosR* | 208964.PA3391 | 22 |
| 28 | *wbpA* | 208964.PA3159 | 22 |
| 29 | *wbpL* | 208964.PA3145 | 22 |
| 30 | PA1400 | 208964.PA1400 | 21 |
| 31 | PA3504 | 208964.PA3504 | 21 |
| 32 | PA3506 | 208964.PA3506 | 21 |
| 33 | *nirG* | 208964.PA0513 | 21 |
| 34 | *nosL* | 208964.PA3396 | 21 |
| 35 | *wzx* | 208964.PA3153 | 21 |
| 36 | PA2324 | 208964.PA2324 | 20 |
| 37 | PA2325 | 208964.PA2325 | 20 |
| 38 | PA2596 | 208964.PA2596 | 20 |
| 39 | PA3512 | 208964.PA3512 | 20 |
| 40 | PA3912 | 208964.PA3912 | 20 |
| 41 | NA*rG* | 208964.PA3875 | 20 |
| 42 | *nirJ* | 208964.PA0511 | 20 |
| 43 | *pvdJ* | 208964.PA2400 | 20 |
| 44 | PA2347 | 208964.PA2347 | 19 |
| 45 | PA3157 | 208964.PA3157 | 19 |
| 46 | PA3535 | 208964.PA3535 | 19 |
| 47 | PA3760 | 208964.PA3760 | 19 |
| 48 | *nirL* | 208964.PA0514 | 19 |
| 49 | *norC* | 208964.PA0523 | 19 |
| 50 | *pscQ* | 208964.PA1694 | 19 |
| 51 | *wzz* | 208964.PA3160 | 19 |
| 52 | *xylZ* | 208964.PA2516 | 19 |

Rest 558 genes with node degree score ‘≤18’ are not listed.

**Table S8. Top six cytoHubba ranked up-regulated genes from among the top-52 in Table S7**

| **No.** | **Gene ID** | **Number of methods ranking this protein among top 10** | **Names of 12 ranking methods of CytoHubba and rank score provided by them** | | | | | | | | | | | |
| --- | --- | --- | --- | --- | --- | --- | --- | --- | --- | --- | --- | --- | --- | --- |
|  |  |  | **Degree** | **MNC** | **DMNC** | **MCC** | **Bottleneck** | **EcCentricity** | **Closeness** | **Radiality** | **Betweenness** | **Stress** | **CC** | **EPC** |
| 1 | PA3156 | 8 | 17 | 17 | 1.01 | 1.87E+11 | 3 | - | 23.76 | 3.68 | - | - | - | 25.32 |
| 2 | PA3158 | 7 | 18 | 18 | - | 1.87E+11 | - | - | 24.26 | 3.70 | - | 1390 | - | 25.48 |
| 3 | PA3150 | 7 | 18 | 18 | - | 1.87E+11 | - | - | 24.26 | 3.70 | - | 1390 | - | 25.44 |
| 4 | PA3149 | 7 | 17 | 17 | 1.01 | 1.87E+11 | - | - | 23.76 | 3.68 | - | - | - | 25.33 |
| 5 | PA3147 | 6 | 18 | 18 | - | 1.87E+11 | - | - | 24.26 | 3.70 | - | 1390 | - | - |
| 6 | PA3151 | 6 | 16 | 16 | 1.01 | 1.87E+11 | - | - | - | - | - | - | 0.97 | 25.35 |

"-": This method did not rank the shown protein among top 10.

MNC: Maximum Neighborhood Component; DMNC: Density of Maximum Neighborhood Component; MCC: Maximal Clique Centrality; CC: Clustering Co-efficient; EPC: Edge Percolated Component

**Table S9. List of down regulated genes in Enteropan-exposed *P. aeruginosa* satisfying the dual criteria of log fold-change ≥2 and FDR≤0.001**

| **Sr. No** | **Gene ID** | **Symbol** | **Product name** | **Log FC** | **FDR** |
| --- | --- | --- | --- | --- | --- |
| 1 | PA3266 | *capB* | major cold shock protein CspA | 6.63 | 7.761E-13 |
| 2 | PA0905 | *rsmA* | carbon storage regulator | 6.48 | 1.8058E-12 |
| 3 | PA3126 | *ibpA* | heat-shock protein IbpA | 6.32 | 4.361E-12 |
| 4 | PA2853 | *oprI* | outer membrane lipoprotein OprI | 6.26 | 5.52E-12 |
| 5 | PA4747 | *secG* | preprotein translocase subunit SecG | 6.11 | 1.3333E-11 |
| 6 | PA1159 | NA | cold-shock protein | 6.08 | 1.788E-11 |
| 7 | PA4432 | *rpsI* | 30S ribosomal protein S9 | 5.93 | 3.7893E-11 |
| 8 | PA4945 | *miaA* | tRNA delta (2)-isopentenylpyrophosphate transferase | 5.83 | 6.3081E-11 |
| 9 | PA0456 | NA | cold-shock protein | 5.76 | 9.1501E-11 |
| 10 | PA4463 | NA | hypothetical protein | 5.75 | 1.0075E-10 |
| 11 | PA3530 | NA | hypothetical protein | 5.73 | 1.3508E-10 |
| 12 | PA5526 | NA | hypothetical protein | 5.64 | 2.4488E-10 |
| 13 | PA2619 | *infA* | translation initiation factor IF-1 | 5.63 | 2.421E-10 |
| 14 | PA2604 | NA | hypothetical protein | 5.57 | 2.4488E-10 |
| 15 | PA3031 | NA | hypothetical protein | 5.54 | 2.6678E-10 |
| 16 | PA2966 | *acpP* | acyl carrier protein | 5.46 | 3.8445E-10 |
| 17 | PA3496 | NA | hypothetical protein | 5.40 | 6.7265E-10 |
| 18 | PA1802 | *clpX* | ATP-dependent protease ATP-binding subunit ClpX | 5.34 | 7.0703E-10 |
| 19 | PA0665 | NA | iron-sulfur cluster insertion protein ErpA | 5.32 | 9.4678E-10 |
| 20 | PA4669 | *Ipk* | 4-diphosphocytidyl-2C-methyl-D-erythritol kinase | 5.31 | 9.033E-10 |
| 21 | PA3601 | NA | 50S ribosomal protein L31 type B | 5.30 | 1.1507E-09 |
| 22 | PA2738 | *himA* | integration host factor subunit alpha | 5.26 | 1.2062E-09 |
| 23 | PA1804 | *hupB* | DNA-binding protein HU | 5.25 | 1.1942E-09 |
| 24 | PA3229 | NA | hypothetical protein | 5.23 | 1.893E-09 |
| 25 | PA3822 | NA | preprotein translocase subunit YajC | 5.21 | 1.748E-09 |
| 26 | PA5049 | *rpmE* | 50S ribosomal protein L31 | 5.17 | 1.893E-09 |
| 27 | PA1343 | NA | hypothetical protein | 5.17 | 1.9576E-09 |
| 28 | PA0621 | NA | hypothetical protein | 5.14 | 2.6641E-09 |
| 29 | PA1178 | *oprH* | PhoP/Q and low Mg2+ inducible outer membrane protein H1 | 5.11 | 2.4212E-09 |
| 30 | PA5053 | *hslV* | ATP-dependent protease peptidase subunit | 5.03 | 4.2563E-09 |
| **Sr. No** | **Gene ID** | **Symbol** | **Product name** | **Log FC** | **FDR** |
| 31 | PA0623 | NA | bacteriophage protein | 5.00 | 5.2599E-09 |
| 32 | PA0622 | NA | bacteriophage protein | 4.95 | 6.4156E-09 |
| 33 | PA4876 | *osmE* | OsmE family transcriptional regulator | 4.94 | 7.5021E-09 |
| 34 | PA3623 | NA | hypothetical protein | 4.90 | 8.839E-09 |
| 35 | PA4386 | *groES* | co-chaperonin GroES | 4.82 | 1.4969E-08 |
| 36 | PA2621 | NA | ATP-dependent Clp protease adapter protein Clp | 4.80 | 1.8185E-08 |
| 37 | PA4741 | *rpsO* | 30S ribosomal protein S15 | 4.78 | 1.8185E-08 |
| 38 | PA4387 | NA | phage exclusion suppressor FxsA | 4.77 | 2.0181E-08 |
| 39 | PA4406 | *lpxC* | UDP-3-O-[3-hydroxymyristoyl] N-acetylglucosamine deacetylase | 4.76 | 2.0531E-08 |
| 40 | PA2830 | *htpX* | protease HtpX | 4.75 | 2.121E-08 |
| 41 | PA2637 | *nuoA* | NADH-quinone oxidoreductase subunit A | 4.70 | 3.0696E-08 |
| 42 | PA4264 | *rpsJ* | 30S ribosomal protein S10 | 4.69 | 2.7571E-08 |
| 43 | PA4268 | *rpsL* | 30S ribosomal protein S12 | 4.67 | 3.3193E-08 |
| 44 | PA4550 | *fimU* | type 4 fimbrial biogenesis protein FimU | 4.67 | 3.3482E-08 |
| 45 | PA4241 | *rpsM* | 30S ribosomal protein S13 | 4.58 | 6.235E-08 |
| 46 | PA0805 | NA | hypothetical protein | 4.58 | 5.2105E-08 |
| 47 | PA5288 | *glnK* | nitrogen regulatory protein P-II 2 | 4.58 | 5.4587E-08 |
| 48 | PA1985 | *pqqA* | coenzyme PQQ synthesis protein A | 4.57 | 8.1886E-08 |
| 49 | PA2883 | NA | hypothetical protein | 4.54 | 6.7776E-08 |
| 50 | PA0635 | NA | hypothetical protein | 4.44 | 1.3231E-07 |
| 51 | PA1596 | *htpG* | chaperone protein HtpG | 4.42 | 1.2852E-07 |
| 52 | PA1847 | NA | Fe/S biogenesis protein NfuA | 4.40 | 1.4521E-07 |
| 53 | PA0634 | NA | hypothetical protein | 4.40 | 1.7148E-07 |
| 54 | PA2826 | NA | glutathione peroxidase | 4.39 | 1.8238E-07 |
| 55 | PA3745 | *rpsP* | 30S ribosomal protein S16 | 4.38 | 1.7728E-07 |
| 56 | PA4563 | *rpsT* | 30S ribosomal protein S20 | 4.38 | 1.699E-07 |
| 57 | PA0633 | NA | hypothetical protein | 4.33 | 2.068E-07 |
| 58 | PA1777 | *oprF* | outer membrane porin F | 4.32 | 2.1424E-07 |
| 59 | PA3235 | NA | hypothetical protein | 4.29 | 3.8138E-07 |
| 60 | PA1198 | NA | hypothetical protein | 4.25 | 3.3656E-07 |
| 61 | PA4249 | *rpsH* | 30S ribosomal protein S8 | 4.21 | 4.3194E-07 |
| 62 | PA3245 | *minE* | cell division topological specificity factor MinE | 4.21 | 5.8917E-07 |
| 63 | PA1592 | NA | hypothetical protein | 4.18 | 4.5957E-07 |
| 64 | PA0624 | NA | hypothetical protein | 4.18 | 5.1763E-07 |
| 65 | PA3815 | *iscR* | HTH-type transcriptional regulator | 4.17 | 5.2217E-07 |
| **Sr. No** | **Gene ID** | **Symbol** | **Product name** | **Log FC** | **FDR** |
| 66 | PA2741 | *rplT* | 50S ribosomal protein L20 | 4.17 | 5.2629E-07 |
| 67 | PA2207 | NA | bacteriophage protein | 4.16 | 5.3019E-07 |
| 68 | PA0038 | NA | hypothetical protein | 4.15 | 5.7104E-07 |
| 69 | PA4752 | *ftsJ* | cell division protein FtsJ | 4.13 | 6.6714E-07 |
| 70 | PA4567 | *rpmA* | 50S ribosomal protein L27 | 4.12 | 6.7979E-07 |
| 71 | PA4251 | *rplE* | 50S ribosomal protein L5 | 4.11 | 7.7139E-07 |
| 72 | PA4761 | *d*NA*K* | molecular chaperone DnaK | 4.10 | 7.2619E-07 |
| 73 | PA1831 | NA | hypothetical protein | 4.09 | 8.2392E-07 |
| 74 | PA3812 | *iscA* | iron-binding protein IscA | 4.08 | 1.0485E-06 |
| 75 | PA0258 | NA | hypothetical protein | 4.08 | 1.0765E-06 |
| 76 | PA4568 | *rplU* | 50S ribosomal protein L21 | 4.05 | 9.5746E-07 |
| 77 | PA4237 | *rplQ* | 50S ribosomal protein L17 | 4.05 | 1.0021E-06 |
| 78 | PA1584 | *sdhB* | succinate dehydrogenase iron-sulfur subunit | 4.04 | 1.0485E-06 |
| 79 | PA0736a | NA | hypothetical protein | 4.03 | 1.0987E-06 |
| 80 | PA1179 | *phoP* | two-component response regulator PhoP | 4.02 | 1.2631E-06 |
| 81 | PA0645 | NA | hypothetical protein | 4.01 | 1.5332E-06 |
| 82 | PA2970 | *rpmF* | 50S ribosomal protein L32 | 4.00 | 1.4326E-06 |
| 83 | PA1053 | NA | hypothetical protein | 3.97 | 1.4326E-06 |
| 84 | PA5332 | *Crc* | catabolite repression control protein | 3.96 | 1.6483E-06 |
| 85 | PA1610 | *fabA* | 3-hydroxydecanoyl-ACP dehydratase | 3.92 | 2.0256E-06 |
| 86 | PA3049 | *Rmf* | ribosome modulation factor | 3.91 | 1.9734E-06 |
| 87 | PA4944 | *Hfq* | RNA-binding protein Hfq | 3.90 | 2.047E-06 |
| 88 | PA5570 | *rpmH* | 50S ribosomal protein L34 | 3.89 | 2.2916E-06 |
| 89 | PA3928 | NA | hypothetical protein | 3.88 | 3.0977E-06 |
| 90 | PA4451 | NA | hypothetical protein | 3.87 | 2.6749E-06 |
| 91 | PA3808 | NA | hypothetical protein | 3.86 | 2.9232E-06 |
| 92 | PA0996 | *pqsA* | anthranilate--CoA ligase | 3.82 | 3.1437E-06 |
| 93 | PA0779 | NA | ATP-dependent protease | 3.82 | 3.1073E-06 |
| 94 | PA2638 | *nuoB* | NADH-quinone oxidoreductase subunit B | 3.82 | 3.395E-06 |
| 95 | PA1581 | *sdhC* | succinate dehydrogenase subunit C | 3.79 | 3.7159E-06 |
| 96 |  | NA | hypothetical protein | 3.79 | 3.9236E-06 |
| 97 | PA3299 | *fadD1* | long-chain-fatty-acid--CoA ligase | 3.78 | 3.9538E-06 |
| 98 | PA0039 | NA | hypothetical protein | 3.77 | 4.1824E-06 |
| 99 | PA1970 | NA | hypothetical protein | 3.77 | 6.2279E-06 |
| 100 | PA2501 | NA | hypothetical protein | 3.75 | 8.0269E-06 |
| 101 | PA2860 | NA | hypothetical protein | 3.73 | 5.529E-06 |
| **Sr. No** | **Gene ID** | **Symbol** | **Product name** | **Log FC** | **FDR** |
| 102 | PA1800 | *Tig* | trigger factor | 3.73 | 5.085E-06 |
| 103 | PA1755 | NA | hypothetical protein | 3.73 | 6.0821E-06 |
| 104 | PA3009 | NA | hypothetical protein | 3.73 | 5.5826E-06 |
| 105 | PA4542 | *clpB* | chaperone protein ClpB | 3.73 | 5.1376E-06 |
| 106 | PA4475 | NA | hypothetical protein | 3.72 | 5.5826E-06 |
| 107 | PA4456 | NA | ABC transporter ATP-binding protein | 3.72 | 5.5826E-06 |
| 108 | PA4395 | NA | nucleotide-binding protein | 3.71 | 6.0559E-06 |
| 109 | PA4751 | *ftsH* | cell division protein FtsH | 3.71 | 5.5826E-06 |
| 110 | PA4575 | NA | hypothetical protein | 3.71 | 5.9754E-06 |
| 111 | PA4762 | *grpE* | heat shock protein GrpE | 3.69 | 6.1602E-06 |
| 112 | PA0320 | NA | hypothetical protein | 3.68 | 7.7555E-06 |
| 113 | PA2586 | *gacA* | response regulator GacA | 3.67 | 7.0246E-06 |
| 114 | PA0995 | *ogt* | methylated-DNA--protein-cysteine methyltransferase | 3.67 | 7.7555E-06 |
| 115 | PA5240 | *trxA* | Thioredoxin | 3.66 | 7.0875E-06 |
| 116 | PA4306 | *flp* | type IVb pilin Flp | 3.66 | 8.6046E-06 |
| 117 | PA5424 | NA | hypothetical protein | 3.66 | 7.6997E-06 |
| 118 | PA1776 | *sigX* | RNA polymerase sigma factor SigX | 3.65 | 7.8723E-06 |
| 119 | PA4739 | NA | hypothetical protein | 3.64 | 1.0396E-05 |
| 120 | PA3533 | NA | hypothetical protein | 3.63 | 8.2688E-06 |
| 121 | PA3385 | *amrZ* | alginate and motility regulator Z | 3.62 | 8.6046E-06 |
| 122 | PA4971 | *aspP* | adenosine diphosphate sugar pyrophosphatase | 3.62 | 9.6659E-06 |
| 123 | PA2960 | *pilZ* | type 4 fimbrial biogenesis protein PilZ | 3.60 | 1.2687E-05 |
| 124 | PA2851 | *efp* | elongation factor P | 3.60 | 1.0377E-05 |
| 125 | PA0490 | NA | hypothetical protein | 3.58 | 1.2505E-05 |
| 126 | PA4245 | *rpmD* | 50S ribosomal protein L30 | 3.58 | 1.7314E-05 |
| 127 | PA0628 | NA | hypothetical protein | 3.58 | 1.1723E-05 |
| 128 | PA1769 | NA | phosphoenolpyruvate synthase regulatory protein | 3.58 | 1.1783E-05 |
| 129 | PA3201 | NA | intracellular septation protein A | 3.57 | 1.2666E-05 |
| 130 | PA4935 | *rpsF* | 30S ribosomal protein S6 | 3.55 | 1.2832E-05 |
| 131 | PA4385 | *groEL* | molecular chaperone GroEL | 3.54 | 1.2823E-05 |
| 132 | PA4714 | NA | hypothetical protein | 3.54 | 1.3806E-05 |
| 133 | PA3162 | *rpsA* | 30S ribosomal protein S1 | 3.53 | 1.3389E-05 |
| 134 | PA0619 | NA | bacteriophage protein | 3.52 | 1.6798E-05 |
| 135 | PA4276 | *secE* | preprotein translocase subunit SecE | 3.51 | 1.6245E-05 |
| 136 | PA5561 | *atpI* | ATP synthase subunit I | 3.50 | 1.9018E-05 |
| **Sr. No** | **Gene ID** | **Symbol** | **Product name** | **Log FC** | **FDR** |
| 137 | PA4090 | NA | hypothetical protein | 3.49 | 1.8936E-05 |
| 138 | PA4499 | NA | transcriptional regulator | 3.48 | 1.7788E-05 |
| 139 | PA3809 | *fdx2* | (2Fe-2S) ferredoxin | 3.47 | 2.1541E-05 |
| 140 | PA0646 | NA | hypothetical protein | 3.47 | 1.9778E-05 |
| 141 | PA3647 | NA | hypothetical protein | 3.47 | 2.2257E-05 |
| 142 | PA1035 | NA | hypothetical protein | 3.45 | 2.5059E-05 |
| 143 | PA4061 | NA | Thioredoxin | 3.44 | 2.3019E-05 |
| 144 | PA0643 | NA | hypothetical protein | 3.43 | 2.4912E-05 |
| 145 | PA3351 | *flgM* | protein FlgM | 3.42 | 2.3889E-05 |
| 146 | PA0640 | NA | bacteriophage protein | 3.42 | 2.6587E-05 |
| 147 | PA1830 | NA | hypothetical protein | 3.41 | 2.6521E-05 |
| 148 | PA0644 | NA | hypothetical protein | 3.40 | 3.8331E-05 |
| 149 | PA2957 | NA | transcriptional regulator | 3.39 | 3.0822E-05 |
| 150 | PA0618 | NA | bacteriophage protein | 3.39 | 3.0334E-05 |
| 151 | PA4661 | *pagL* | lipid A 3-O-deacylase | 3.39 | 2.9857E-05 |
| 152 | PA1544 | *Anr* | transcriptional regulator Anr | 3.38 | 2.9857E-05 |
| 153 | PA0625 | NA | hypothetical protein | 3.37 | 3.0878E-05 |
| 154 | PA1882 | NA | Transporter | 3.36 | 3.8331E-05 |
| 155 | PA0762 | *algU* | RNA polymerase sigma factor AlgU | 3.35 | 3.3711E-05 |
| 156 | PA0408 | *pilG* | pilus biosynthesis/twitching motility protein PilG | 3.35 | 3.5749E-05 |
| 157 | PA2623 | *Icd* | isocitrate dehydrogenase | 3.34 | 3.4664E-05 |
| 158 | PA0329 | NA | hypothetical protein | 3.34 | 3.5624E-05 |
| 159 | PA2196 | NA | transcriptional regulator | 3.32 | 4.8798E-05 |
| 160 | PA1414 | NA | hypothetical protein | 3.32 | 3.8315E-05 |
| 161 | PA0139 | *ahpC* | alkyl hydroperoxide reductase | 3.31 | 3.9996E-05 |
| 162 | PA0506 | NA | acyl-CoA dehydrogenase | 3.30 | 4.2353E-05 |
| 163 | PA5528 | NA | hypothetical protein | 3.30 | 4.2743E-05 |
| 164 | PA0615 | NA | hypothetical protein | 3.29 | 5.3202E-05 |
| 165 | PA0636 | NA | hypothetical protein | 3.29 | 4.7284E-05 |
| 166 | PA4248 | *rplF* | 50S ribosomal protein L6 | 3.28 | 5.0377E-05 |
| 167 | PA0286 | *desA* | delta-9 fatty acid desaturase DesA | 3.27 | 5.0229E-05 |
| 168 | PA4261 | *rplW* | 50S ribosomal protein L23 | 3.27 | 6.0809E-05 |
| 169 | PA1963 | NA | hypothetical protein | 3.26 | 6.3188E-05 |
| 170 | PA0639 | NA | hypothetical protein | 3.26 | 5.9738E-05 |
| 171 | PA1100 | *fliE* | flagellar hook-basal body complex protein FliE | 3.25 | 6.0542E-05 |
| 172 | PA5276 | *lppL* | lipopeptide LppL | 3.24 | 6.323E-05 |
| 173 | PA1801 | *clpP* | ATP-dependent Clp protease proteolytic subunit | 3.24 | 5.9579E-05 |
| **Sr. No** | **Gene ID** | **Symbol** | **Product name** | **Log FC** | **FDR** |
| 174 | PA3600 | NA | 50S ribosomal protein L36 | 3.24 | 0.0001 |
| 175 | PA0577 | *dna*G | DNA primase | 3.24 | 5.9075E-05 |
| 176 | PA1775 | *cmpX* | hypothetical protein | 3.23 | 6.0807E-05 |
| 177 | PA2485 | NA | hypothetical protein | 3.23 | 7.0464E-05 |
| 178 | PA4765 | *omlA* | outer membrane lipoprotein OmlA | 3.23 | 6.321E-05 |
| 179 | PA4453 | NA | hypothetical protein | 3.22 | 6.8002E-05 |
| 180 | PA0003 | *recF* | DNA replication and repair protein RecF | 3.22 | 6.5082E-05 |
| 181 | PA4870 | NA | hypothetical protein | 3.22 | 6.903E-05 |
| 182 | PA0626 | NA | hypothetical protein | 3.21 | 6.9739E-05 |
| 183 | PA5559 | *atpE* | ATP synthase subunit C | 3.20 | 8.2487E-05 |
| 184 | PA2797 | NA | hypothetical protein | 3.19 | 8.3721E-05 |
| 185 | PA2667 | NA | hypothetical protein | 3.19 | 7.5932E-05 |
| 186 | PA1749 | NA | hypothetical protein | 3.18 | 7.9156E-05 |
| 187 | PA4724.1 | NA | hypothetical protein | 3.17 | 0.0001 |
| 188 | PA1430 | *lasR* | transcriptional regulator LasR | 3.17 | 8.3473E-05 |
| 189 | PA4421 | NA | cell division protein MraZ | 3.16 | 8.4673E-05 |
| 190 | PA2755 | *eco* | Ecotin | 3.15 | 9.1806E-05 |
| 191 | PA1548 | NA | hypothetical protein | 3.15 | 0.0001 |
| 192 | PA0865 | *hpd* | 4-hydroxyphenylpyruvate dioxygenase | 3.15 | 8.8442E-05 |
| 193 | PA0576 | *rpoD* | RNA polymerase sigma factor RpoD | 3.15 | 8.5466E-05 |
| 194 | PA1840 | NA | hypothetical protein | 3.15 | 0.0001 |
| 195 | PA4462 | *rpoN* | RNA polymerase factor sigma-54 | 3.14 | 9.2754E-05 |
| 196 | PA3814 | *iscS* | cysteine desulfurase | 3.14 | 9.2754E-05 |
| 197 | PA0648 | NA | hypothetical protein | 3.13 | 0.0001 |
| 198 | PA3006 | *psrA* | transcriptional regulator PsrA | 3.13 | 9.6962E-05 |
| 199 | PA4611 | NA | hypothetical protein | 3.11 | 0.0001 |
| 200 | PA5239 | *rho* | transcription termination factor Rho | 3.10 | 0.0001 |
| 201 | PA3014 | *faoA* | fatty acid oxidation complex subunit alpha | 3.10 | 0.0001 |
| 202 | PA3645 | *fabZ* | 3-hydroxyacyl-[acyl-carrier-protein] dehydratase FabZ | 3.10 | 0.0001 |
| 203 | PA1112b | NA | hypothetical protein | 3.09 | 0.0001 |
| 204 | PA1580 | *gltA* | citrate synthase | 3.09 | 0.0001 |
| 205 | PA4500 | NA | ABC transporter | 3.08 | 0.0001 |
| 206 | PA4454 | NA | hypothetical protein | 3.08 | 0.0001 |
| 207 | PA3811 | *hscB* | co-chaperone HscB | 3.08 | 0.0001 |
| 208 | PA4238 | *rpoA* | DNA-directed RNA polymerase subunit alpha | 3.07 | 0.0001 |
| **Sr. No** | **Gene ID** | **Symbol** | **Product name** | **Log FC** | **FDR** |
| 209 | PA0638 | NA | bacteriophage protein | 3.07 | 0.0001 |
| 210 | PA0627 | NA | hypothetical protein | 3.07 | 0.0001 |
| 211 | PA2952 | *etfB* | electron transfer flavoprotein subunit beta | 3.07 | 0.0001 |
| 212 | PA3622 | *rpoS* | RNA polymerase sigma factor RpoS | 3.07 | 0.0001 |
| 213 | PA1869 | NA | acyl carrier protein | 3.07 | 0.0001 |
| 214 | PA3224 | NA | hypothetical protein | 3.06 | 0.0001 |
| 215 | PA1793 | *ppiB* | peptidyl-prolyl cis-trans isomerase B | 3.06 | 0.0001 |
| 216 | PA5068 | *tatA* | twin-arginine translocation protein TatA | 3.05 | 0.0001 |
| 217 | PA0968 | NA | hypothetical protein | 3.04 | 0.0001 |
| 218 | PA5560 | *atpB* | ATP synthase subunit A | 3.04 | 0.0001 |
| 219 | PA0579 | *rpsU* | 30S ribosomal protein S21 | 3.04 | 0.0001 |
| 220 | PA1742 | NA | Amidotransferase | 3.03 | 0.0001 |
| 221 | PA2023 | *galU* | UTP-glucose-1-phosphate uridylyltransferase | 3.03 | 0.0001 |
| 222 | PA2798 | NA | two-component response regulator | 3.02 | 0.0001 |
| 223 | PA4275 | *nusG* | transcription antitermination protein NusG | 3.02 | 0.0001 |
| 224 | PA2743 | *infC* | translation initiation factor IF-3 | 3.01 | 0.0001 |
| 225 | PA4693 | *pssA* | phosphatidylserine synthase | 3.01 | 0.0001 |
| 226 | PA0336 | *ygdP* | RNA pyrophosphohydrolase | 3.01 | 0.0001 |
| 227 | PA4253 | *rplN* | 50S ribosomal protein L14 | 3.00 | 0.0001 |
| 228 | PA0532 | NA | hypothetical protein | 3.00 | 0.0001 |
| 229 | PA4753 | NA | hypothetical protein | 2.99 | 0.0001 |
| 230 | PA2971 | NA | hypothetical protein | 2.99 | 0.0001 |
| 231 | PA0872 | *phhA* | phenylalanine 4-monooxygenase | 2.98 | 0.0002 |
| 232 | PA0392 | NA | hypothetical protein | 2.98 | 0.0002 |
| 233 | PA4465 | NA | hypothetical protein | 2.98 | 0.0002 |
| 234 | PA1623 | NA | hypothetical protein | 2.97 | 0.0002 |
| 235 | PA5253 | *algP* | alginate regulatory protein AlgP | 2.97 | 0.0002 |
| 236 | PA2407 | NA | adhesion protein | 2.96 | 0.0002 |
| 237 | PA2985 | NA | hypothetical protein | 2.96 | 0.0002 |
| 238 | PA3295 | NA | HIT family protein | 2.96 | 0.0002 |
| 239 | PA5054 | *hslU* | ATP-dependent protease ATP-binding subunit HslU | 2.96 | 0.0002 |
| 240 | PA4433 | *rplM* | 50S ribosomal protein L13 | 2.95 | 0.0002 |
| 241 | PA2780 | NA | hypothetical protein | 2.95 | 0.0002 |
| 242 | PA1814 | NA | hypothetical protein | 2.95 | 0.0002 |
| **Sr. No** | **Gene ID** | **Symbol** | **Product name** | **Log FC** | **FDR** |
| 243 | PA2951 | *etfA* | electron transfer flavoprotein subunit alpha | 2.94 | 0.0002 |
| 244 | PA0997 | *pqsB* | hypothetical protein | 2.93 | 0.0002 |
| 245 | PA5489 | *dsbA* | thiol:disulfide interchange protein DsbA | 2.92 | 0.0002 |
| 246 | PA5011 | *waaC* | heptosyltransferase I | 2.92 | 0.0002 |
| 247 | PA3244 | *minD* | cell division inhibitor MinD | 2.92 | 0.0002 |
| 248 | PA0637 | NA | hypothetical protein | 2.92 | 0.0003 |
| 249 | PA4963 | NA | hypothetical protein | 2.92 | 0.0002 |
| 250 | PA0297 | *spuA* | glutamine amidotransferase | 2.92 | 0.0002 |
| 251 | PA1852 | NA | hypothetical protein | 2.91 | 0.0002 |
| 252 | PA2756 | NA | hypothetical protein | 2.91 | 0.0002 |
| 253 | PA1463 | NA | hypothetical protein | 2.91 | 0.0002 |
| 254 | PA0284 | NA | hypothetical protein | 2.91 | 0.0002 |
| 255 | PA3552 | *arnB* | UDP-4-amino-4-deoxy-L-arabinose--oxoglutarate aminotransferase | 2.90 | 0.0003 |
| 256 | PA4471 | NA | hypothetical protein | 2.90 | 0.0003 |
| 257 | PA1509 | NA | hypothetical protein | 2.90 | 0.0003 |
| 258 | PA4263 | *rplC* | 50S ribosomal protein L3 | 2.90 | 0.0002 |
| 259 | PA1002 | *phnB* | anthranilate synthase component II | 2.90 | 0.0003 |
| 260 | PA2805 | NA | hypothetical protein | 2.89 | 0.0002 |
| 261 | PA0970 | *tolR* | translocation protein TolR | 2.89 | 0.0003 |
| 262 | PA4324 | NA | hypothetical protein | 2.88 | 0.0003 |
| 263 | PA4243 | *secY* | preprotein translocase subunit SecY | 2.88 | 0.0003 |
| 264 | PA1456 | *cheY* | chemotaxis protein CheY | 2.88 | 0.0003 |
| 265 | PA4574 | NA | hypothetical protein | 2.88 | 0.0003 |
| 266 | PA0036 | *trpB* | tryptophan synthase subunit beta | 2.88 | 0.0003 |
| 267 | PA2747a | NA | hypothetical protein | 2.87 | 0.0003 |
| 268 | PA1583 | *sdhA* | succinate dehydrogenase flavoprotein subunit | 2.87 | 0.0003 |
| 269 | PA3017 | NA | hypothetical protein | 2.86 | 0.0003 |
| 270 | PA2146 | NA | hypothetical protein | 2.86 | 0.0003 |
| 271 | PA3257 | *prc* | tail-specific protease | 2.86 | 0.0003 |
| 272 | PA5301 | NA | transcriptional regulator | 2.85 | 0.0003 |
| 273 | PA0363 | *coaD* | phosphopantetheine adenylyltransferase | 2.85 | 0.0003 |
| 274 | PA1076 | NA | hypothetical protein | 2.84 | 0.0003 |
| 275 | PA1797a | NA | hypothetical protein | 2.84 | 0.0003 |
| 276 | PA4551 | *pilV* | type 4 fimbrial biogenesis protein PilV | 2.84 | 0.0003 |
| **Sr. No** | **Gene ID** | **Symbol** | **Product name** | **Log FC** | **FDR** |
| 277 | PA1942 | NA | hypothetical protein | 2.84 | 0.0004 |
| 278 | PA1307 | NA | hypothetical protein | 2.82 | 0.0004 |
| 279 | PA4842 | NA | hypothetical protein | 2.82 | 0.0004 |
| 280 | PA2790 | NA | hypothetical protein | 2.82 | 0.0004 |
| 281 | PA2614 | *lolA* | outer-membrane lipoprotein carrier protein | 2.80 | 0.0004 |
| 282 | PA0943 | NA | hypothetical protein | 2.80 | 0.0004 |
| 283 | PA0567 | NA | hypothetical protein | 2.80 | 0.0005 |
| 284 | PA4419 | *ftsL* | cell division protein FtsL | 2.80 | 0.0006 |
| 285 | PA1494 | NA | hypothetical protein | 2.80 | 0.0004 |
| 286 | PA4259 | *rpsS* | 30S ribosomal protein S19 | 2.80 | 0.0004 |
| 287 | PA0563 | NA | hypothetical protein | 2.79 | 0.0004 |
| 288 | PA4239 | *rpsD* | 30S ribosomal protein S4 | 2.79 | 0.0004 |
| 289 | PA2742 | *rpmI* | 50S ribosomal protein L35 | 2.79 | 0.0004 |
| 290 | PA4748 | *tpiA* | triosephosphate isomerase | 2.79 | 0.0004 |
| 291 | PA4114 | NA | spermidine acetyltransferase | 2.78 | 0.0004 |
| 292 | PA4674 | NA | hypothetical protein | 2.78 | 0.0004 |
| 293 | PA2491 | NA | Oxidoreductase | 2.78 | 0.0004 |
| 294 | PA0422 | NA | hypothetical protein | 2.78 | 0.0005 |
| 295 | PA0667 | NA | hypothetical protein | 2.78 | 0.0004 |
| 296 | PA5369 | *pstS* | phosphate ABC transporter substrate-binding protein | 2.77 | 0.0005 |
| 297 | PA3940 | NA | DNA binding protein | 2.76 | 0.0005 |
| 298 | PA4738 | NA | hypothetical protein | 2.76 | 0.0006 |
| 299 | PA5482 | NA | hypothetical protein | 2.75 | 0.0007 |
| 300 | PA0951a | NA | hypothetical protein | 2.75 | 0.0007 |
| 301 | PA1013 | *purC* | phosphoribosylaminoimidazole-succinocarboxamide synthase | 2.75 | 0.0005 |
| 302 | PA0394 | NA | hypothetical protein | 2.74 | 0.0005 |
| 303 | PA5128 | *secB* | preprotein translocase subunit SecB | 2.74 | 0.0005 |
| 304 | PA1308 | NA | hypothetical protein | 2.74 | 0.0006 |
| 305 | PA2312a | NA | hypothetical protein | 2.73 | 0.0007 |
| 306 | PA2737 | NA | hypothetical protein | 2.72 | 0.0006 |
| 307 | PA4498 | NA | Metallopeptidase | 2.72 | 0.0006 |
| 308 | PA1821 | NA | enoyl-CoA hydratase | 2.71 | 0.0007 |
| 309 | PA4271 | *rplL* | 50S ribosomal protein L7/L12 | 2.71 | 0.0006 |
| 310 | PA0250 | NA | hypothetical protein | 2.70 | 0.0007 |
| 311 | PA4141 | NA | hypothetical protein | 2.69 | 0.0007 |
| 312 | PA3835 | NA | hypothetical protein | 2.69 | 0.0007 |
| 313 | PA3472 | NA | hypothetical protein | 2.68 | 0.0007 |
| **Sr. No** | **Gene ID** | **Symbol** | **Product name** | **Log FC** | **FDR** |
| 314 | PA2446 | *gcvH2* | glycine cleavage system protein H | 2.67 | 0.0008 |
| 315 | PA3578 | NA | hypothetical protein | 2.67 | 0.0007 |
| 316 | PA4257 | *rpsC* | 30S ribosomal protein S3 | 2.67 | 0.0007 |
| 317 | PA3085 | NA | hypothetical protein | 2.67 | 0.0008 |
| 318 | PA3656 | *rpsB* | 30S ribosomal protein S2 | 2.67 | 0.0007 |
| 319 | PA3686 | *adk* | adenylate kinase | 2.67 | 0.0008 |
| 320 | PA2746a | NA | hypothetical protein | 2.67 | 0.0008 |
| 321 | PA2779 | NA | hypothetical protein | 2.66 | 0.0009 |
| 322 | PA3813 | *iscU* | scaffold protein | 2.66 | 0.0008 |
| 323 | PA1443 | *fliM* | flagellar motor switch protein FliM | 2.66 | 0.0008 |
| 324 | PA4570 | NA | hypothetical protein | 2.65 | 0.0008 |
| 325 | PA0973 | *oprL* | peptidoglycan associated lipoprotein OprL | 2.65 | 0.0008 |
| 326 | PA1034 | NA | hypothetical protein | 2.64 | 0.0009 |
| 327 | PA2321 | NA | Gluconokinase | 2.64 | 0.0009 |
| 328 | PA1527 | NA | hypothetical protein | 2.64 | 0.0008 |
| 329 | PA1571 | NA | hypothetical protein | 2.64 | 0.0009 |
| 330 | PA5227 | NA | hypothetical protein | 2.63 | 0.0009 |
| 331 | PA1455 | *fliA* | flagellar biosynthesis sigma factor FliA | 2.63 | 0.0009 |
| 332 | PA4134 | NA | hypothetical protein | 2.63 | 0.001 |
| 333 | PA2854 | NA | hypothetical protein | 2.63 | 0.0009 |
| 334 | PA5200 | *amgR* | osmolarity response regulator | 2.62 | 0.0009 |
| 335 | PA5264 | NA | hypothetical protein | 2.62 | 0.0009 |
| 336 | PA4671 | NA | 50S ribosomal protein L25/general stress protein Ctc | 2.62 | 0.0009 |

Genes are arranged in decreasing order of Fold Change; Databases consulted for gene functions were: NCBI gene database (https://www.ncbi.nlm.nih.gov/nuccore/NC_002516); KEGG (Kyoto Encyclopedia of Genes and Genomes: https://www.genome.jp/kegg/); Uniprot (<https://www.uniprot.org/>). NA: Not Applicable; FDR: False Discovery Rate

**Table S10. Node degree score of the top down-regulated genes**

| **No.** | **Gene ID/Symbol** | **Identifier** | **Node degree** |
| --- | --- | --- | --- |
| 1 | *rpsB* | 208964.PA3656 | 79 |
| 2 | *rplM* | 208964.PA4433 | 76 |
| 3 | *rpsL* | 208964.PA4268 | 76 |
| 4 | *rpoA* | 208964.PA4238 | 75 |
| 5 | *rplU* | 208964.PA4568 | 74 |
| 6 | *ftsH* | 208964.PA4751 | 72 |
| 7 | *rplT* | 208964.PA2741 | 72 |
| 8 | *rpmF* | 208964.PA2970 | 72 |
| 9 | *rplC* | 208964.PA4263 | 70 |
| 10 | *rpsD* | 208964.PA4239 | 70 |
| 11 | *rpsJ* | 208964.PA4264 | 70 |
| 12 | *tig* | 208964.PA1800 | 70 |
| 13 | *rplN* | 208964.PA4253 | 69 |
| 14 | *rpsO* | 208964.PA4741 | 69 |
| 15 | *rpmA* | 208964.PA4567 | 67 |
| 16 | *rpsF* | 208964.PA4935 | 67 |
| 17 | *rpsM* | 208964.PA4241 | 67 |
| 18 | *rpsA* | 208964.PA3162 | 66 |
| 19 | *secY* | 208964.PA4243 | 66 |
| 20 | *rplF* | 208964.PA4248 | 65 |
| 21 | *rplL* | 208964.PA4271 | 65 |
| 22 | *rpsT* | 208964.PA4563 | 65 |
| 23 | *efp* | 208964.PA2851 | 64 |
| 24 | *infC* | 208964.PA2743 | 64 |
| 25 | *rplQ* | 208964.PA4237 | 64 |
| 26 | *rpoD* | 208964.PA0576 | 64 |
| 27 | *dnaK* | 208964.PA4761 | 63 |
| 28 | *groEL* | 208964.PA4385 | 63 |
| 29 | *rpmI* | 208964.PA2742 | 63 |
| 30 | *rpsP* | 208964.PA3745 | 63 |
| 31 | *rplE* | 208964.PA4251 | 62 |
| 32 | *rpmD* | 208964.PA4245 | 62 |
| 33 | *acpP* | 208964.PA2966 | 61 |
| 34 | *infA* | 208964.PA2619 | 60 |
| 35 | *rplY* | 208964.PA4671 | 60 |
| 36 | *rpsH* | 208964.PA4249 | 60 |
| 37 | *secE* | 208964.PA4276 | 60 |
| 38 | *nusG* | 208964.PA4275 | 59 |
| 39 | *rpoS* | 208964.PA3622 | 59 |
| 40 | *rpsC* | 208964.PA4257 | 59 |
| 41 | *rpsI* | 208964.PA4432 | 59 |
| 42 | *rplW* | 208964.PA4261 | 58 |
| 43 | *rpsS* | 208964.PA4259 | 58 |
| 44 | *groES* | 208964.PA4386 | 57 |
| 45 | *atpE* | 208964.PA5559 | 56 |
| 46 | *rpsU* | 208964.PA0579 | 55 |
| 47 | PA4463 | 208964.PA4463 | 54 |
| 48 | *hfq* | 208964.PA4944 | 53 |
| 49 | *rpmE* | 208964.PA5049 | 53 |
| 50 | *rpmJ2* | 208964.PA3600 | 53 |

Rest 277 genes with node degree score ‘≤52’ are not listed

**Table S11. Top ten cytoHubba ranked down-regulated genes from among the top-50 in Table S10**

| **No.** | **Gene ID** | **Gene**  **Name** | **Number of methods ranking this protein among top 10** | **Names of 12 ranking methods of CytoHubba and rank score provided by them** | | | | | | | | | | | |
| --- | --- | --- | --- | --- | --- | --- | --- | --- | --- | --- | --- | --- | --- | --- | --- |
|  |  |  |  | **Degree** | **MNC** | **DMNC** | **MCC** | **Bottleneck** | **EcCentricity** | **Closeness** | **Radiality** | **Betweenness** | **Stress** | **CC** | **EPC** |
| 1 | PA3656 | *rpsB* | 10 | 49 | 49 | - | 1.06E+45 | 1 | 1 | 49 | 2.06 | 14.07 | 336 | - | 26.06 |
| 2 | PA4239 | *rpsD* | 10 | 49 | 49 | - | 1.06E+45 | 2 | 1 | 49 | 2.06 | 14.07 | 336 | - | 25.76 |
| 3 | PA4268 | *rpsL* | 9 | 49 | 49 | - | 1.06E+45 | - | 1 | 49 | 2.06 | 14.07 | 336 | - | 25.55 |
| 4 | PA4251 | *rplE* | 8 | 47 | 47 | - | 1.06E+45 | - | - | 48 | 2.02 | 10.08 | 250 | - | 26.3 |
| 5 | PA4238 | *rpoA* | 7 | 48 | 48 | - | - | - | - | 48.5 | 2.04 | 12.73 | 304 | - | 25.65 |
| 6 | PA2741 | *rplT* | 7 | 47 | 47 | - | - | - | - | 48 | 2.02 | 9.83 | 246 | - | 25.65 |
| 7 | PA4264 | *rpsJ* | 7 | 47 | 47 | - | 1.06E+45 | - | - | 48 | 2.02 | - | 234 | - | 25.73 |
| 8 | PA4253 | *rplN* | 7 | 47 | 47 | - | 1.06E+45 | - | - | 48 | 2.02 | 9.21 | 238 | - | - |
| 9 | PA4271 | *rplL* | 7 | 47 | 47 | - | - | - | - | 48 | 2.02 | 11.89 | 278 | - | 25.32 |
| 10 | PA4248 | *rplF* | 6 | 47 | 47 | - | - | - | - | 48 | 2.02 | 9.88 | 248 | - | - |

"-": This method did not rank the shown protein among top 10

MNC: Maximum Neighborhood Component; DMNC: Density of Maximum Neighborhood Component; MCC: Maximal Clique Centrality; CC: Clustering Co-efficient;EPC: Edge Percolated Component

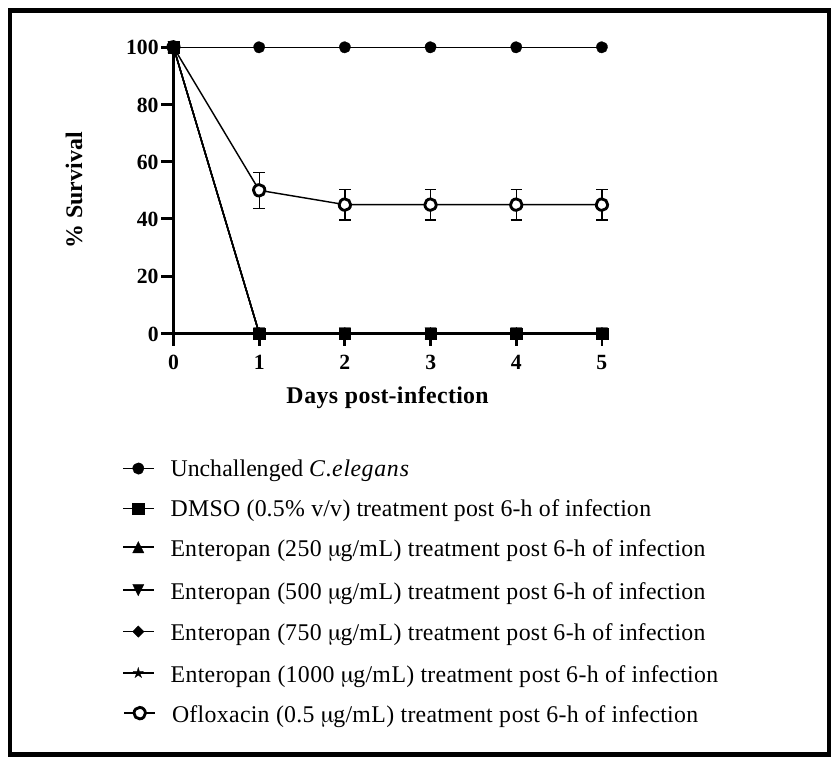

**Figure S1. Enteropan could not rescue the host worm post 6-h of *P. aeruginosa* infection.** Ofloxacin (0.5 µg/mL) employed as positive control post 6-h infection conferred 45% ± 5.47 (*p*<0.001) survival benefit on host worm. Neither DMSO nor Enteropan showed any toxicity towards the worm population at tested concentrations.

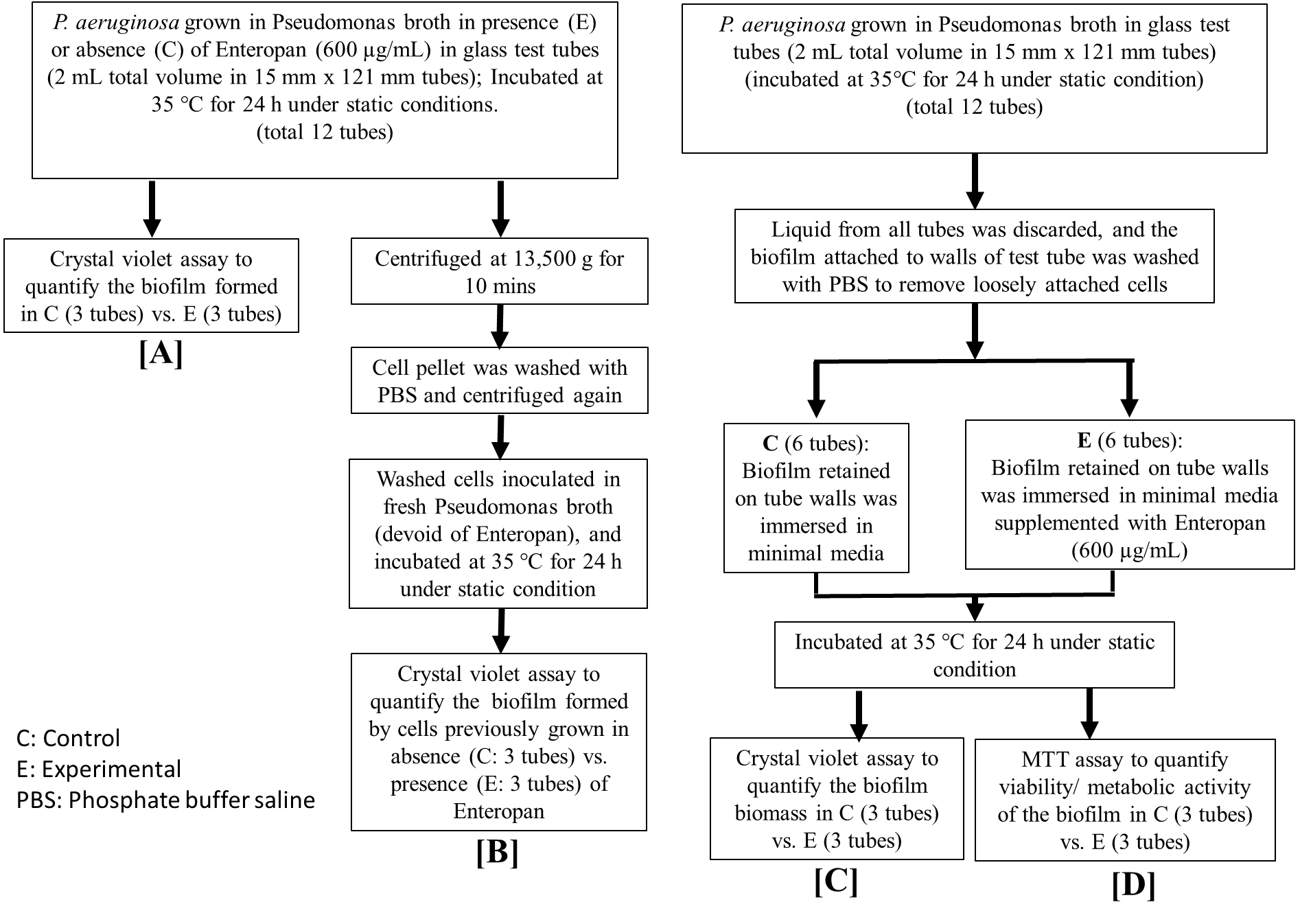

**Figure S2: Flowchart depicting schematic of all biofilm assays**

(A) Quantification of biofilm formation in presence or absence of Enteropan; **(B)** Quantification of biofilm formation by Enteropan-pre-treated vs. non-pre-treated *P. aeruginosa* cells; **(C)** Quantification of biofilm eradication after adding Enteropan onto pre-formed biofilm; **(D)** Quantification of Enteropan’s effect on metabolic activity of pre-formed biofilm

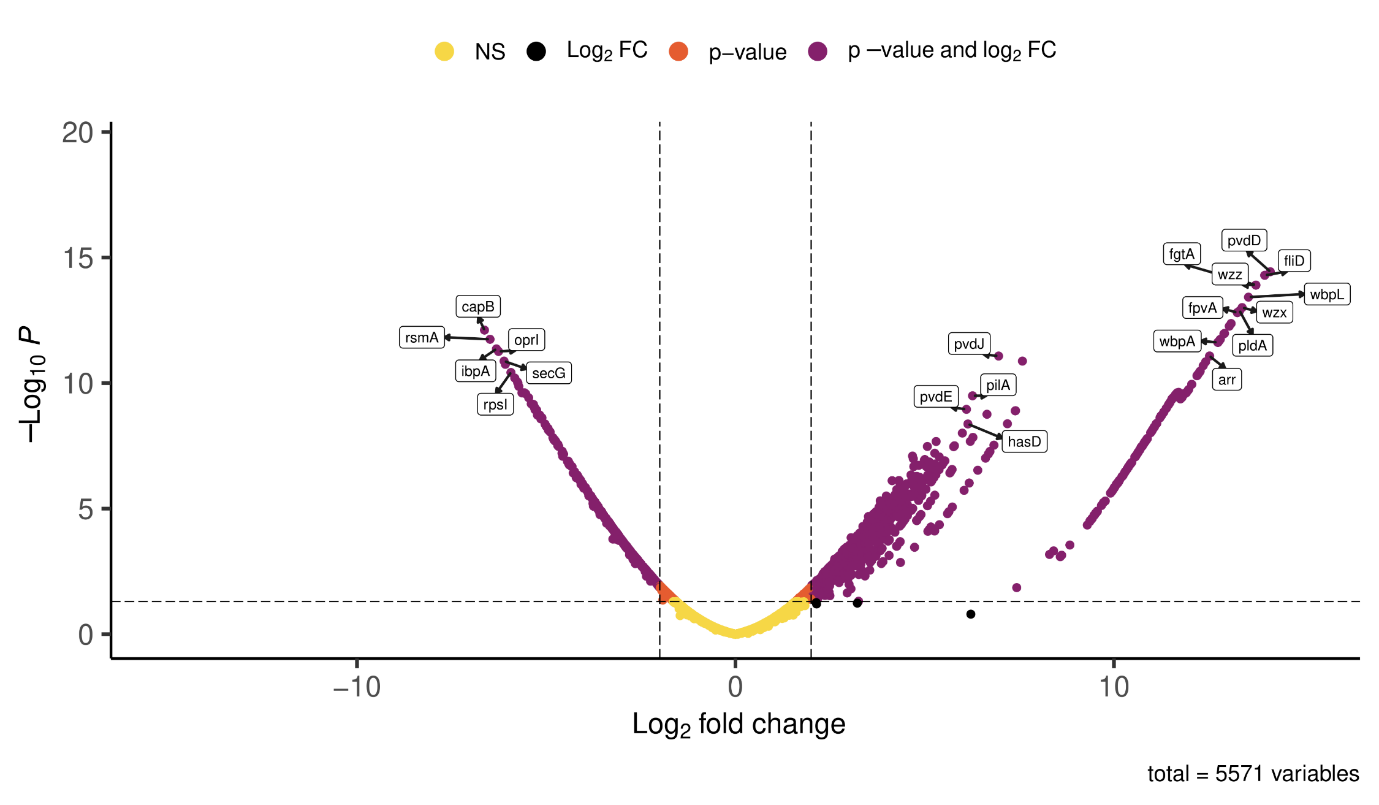

**Figure S3. Volcano plot of experimental versus control samples.** Volcano plot of expressed genes of experimental culture compared to control culture. The y- axis illustrates −log 10 p values, and the x-axis corresponds to a log 2-fold change of gene expression between both cultures. The purple points represent differently expressed genes satisfying the dual criteria of FDR< 0.05 and log fold change ≥2.
